# Comparative analysis of the neural and muscle systems in the subumbrella of hydrozoan jellyfish

**DOI:** 10.64898/2026.08.30.748097

**Authors:** Tigran P. Norekian, Leonid L. Moroz

**Affiliations:** Whitney Laboratory, University of Florida, St. Augustine, FL, USA; Department of Neuroscience and McKnight Brain Institsute, University of Florida, Gainesville, FL, USA

**Keywords:** Hydrozoa, Cnidarians, Nervous System, Muscle System, Evolution

## Abstract

Hydrozoa is a group of relatively simple animals with a well-developed nervous system. The nervous system in all hydrozoan medusae is highly conserved and includes outer and inner nerve rings at the bell margin, a neural network in the manubrium, and radial neural pathways that connect them. However, one element of the nervous system shows substantial variability among species – the subumbrella neural network. We examined the structure of the nervous and muscular systems in the subumbrella of 14 species of hydrozoan medusae. The main conclusion of this study is that the distribution of neural networks in the subumbrella strongly correlates with the distribution of smooth radial muscles. This correlation suggests that smooth radial muscles are the primary target of the subumbrella nervous system. Most species in the order Anthoathecata show a clear tendency to lose the neural networks and radial smooth muscle fibers in the subumbrella region and to concentrate neural elements and smooth muscles only in the radial pathways along the radial canals. All studied species in the order Leptothecata, by contrast, have neural networks in the subumbrella area, as well as radial smooth muscle fibers spread throughout the entire subumbrella region. The correlation between radial smooth muscles and the nervous system is also observed in the radial pathways along the radial canals. All species that have thick bundles of smooth radial muscles along the radial canals have clearly defined and dense neural pathways running along or even embedded within the smooth muscle bundles.

## 1 INTRODUCTION

Hydrozoans are relatively simple yet highly active animals that feed on a variety of organisms. Their complex behavioral repertoire is supported by a well-developed nervous system. The general morphology of hydrozoan medusae and the overall structure of their nervous system are highly conserved across all studied species, and their basic description hasn’t changed since the earlier studies such as Hertwig and Hertwig (1878). The nervous systems of two hydrozoan medusae have been studied most extensively over the years, both electrophysiologically and morphologically. The first species is the anthomedusa *Polyorchis penicillatus* (Singla, 1978a; Anderson, 1979; Anderson & Mackie, 1977; Arkett & Spencer, 1986a, b; Spencer, 1978, 1979, 1981; Spencer & Satterlie, 1980, 1981; Spencer & Arkett, 1984; Satterlie & Spencer, 1983; Weber, 1982; Lin, Gallin & Spencer, 2001). The second species is the trachymedusa *Aglantha digitale* (Singla, 1978b, 1983; Donaldson et al., 1980; Mackie, 1980, 2004; Roberts & Mackie, 1980; Weber et al., 1982; Kerfoot et al., 1985; Mackie & Meech, 1985, 1995a, b, 2000, 2008; Arkett et al., 1988; Meech & Mackie, 1993a, b, 1995; Mackie et al., 2003; Norekian & Moroz, 2020b). To a lesser extent, such hydrozoan species as *Aequorea*, *Sarsia*, *Euphysa*, *Stomotoca*, and *Clytia* (*Phialidium*) have also been investigated to understand the structure and function of their neural systems (Passano, 1965, 1973; Mackie, 1970, 1971, 1975; Mackie & Singla, 1975; Jha & Mackie, 1967; Satterlie & Spencer, 1983; Satterlie, 1985a, b, 2002, 2008).

All hydrozoan medusae have a bell-shaped body, whether elongated and narrow (prolate) or broad and shallow (oblate), and consist of two major parts - the exumbrella and the subumbrella (Fig. 2). Their nervous system is largely concentrated at the margin of the bell, where it consists of two nerve rings - the outer and inner nerve rings. Predominantly, the inner nerve ring contains neurons that trigger rhythmic swimming activity and activate the circular striated swim muscles, which form a myoepithelial sheet in the subumbrella. There are also neural networks in the tentacles and manubrium, as well as radial nerves or neural tracts along the radial canals that connect the marginal nerve rings to the manubrium (Fig. 2). In addition to nervous conduction, hydrozoans have an excitable epithelium capable of transmitting electrical impulses, including both the external epithelium and the subumbrella myoepithelium layer (Mackie & Passano 1968; Mackie, 1965, 1970; Satterlie & Spencer, 1983).

Despite many similarities, there are also considerable differences in neural innervation of the subumbrella. For example, in *Polyorchis* and some other Anthomedusae, the subumbrella consists only of a myoepithelial layer without any neurons or neurites (Singla, 1978a, b; Spencer, 1979; Satterlie & Spencer, 1983). Innervation of the myoepithelial sheet occurs at the bell margin, directly from the inner nerve ring. However, in the Leptomedusae *Aequorea* and *Phialidium*, and in the anthomedusa *Neoturris*, there is also a pronounced neural network covering the entire subumbrella region (Satterlie & Spencer, 1983; Satterlie, 2008; Mackie, Singla & Stell, 1985; Mackie & Meech, 2008). Satterlie (2008) suggested that in *Aequorea* this subumbrella nerve net is involved in the spread of swim contractions, while a comparative investigation of similar subumbrella neural networks (showing FMRFamide immunoreactivity) in several other hydromedusae suggested that they are associated with radial muscles and not with the circular swimming system (Mackie, Singla & Stell, 1985). In the unique trachymedusa *Aglantha*, the subumbrella region contains a system of giant axons with a number of lateral interneurons, which innervate the circular swim muscles and generate the fast escape response (Singla, 1978b; Donaldson, Mackie & Roberts, 1980; Roberts and Mackie, 1980; Kerfoot et al., 1985; Weber et al., 1982; Mackie & Meech, 1985). The goal of the current investigation is to summarize and extend our understanding of the structure of the nervous and muscular systems in hydrozoan jellyfish and to identify possible correlations and differences through extensive analysis of 14 jellyfish species from Northwest Pacific coastal waters off North America.

## 2 MATERIALS AND METHODS

### 2.1 Animals

Adult specimens of 14 hydrozoan jellyfish species were collected from the breakwater in the Northwest Pacific Ocean and held in 1-gal glass jars within large tanks with continuously circulating seawater at 10°C–12°C. For species identification, we used Wrobel and Mills (1998) and other sources. Experiments were conducted at Friday Harbor Laboratories, the University of Washington, during the spring, summer, and fall seasons of 2019–2023.

### 2.2 Immunocytochemistry and Phalloidin staining

Adult animals were fixed overnight (10-12 hours) in 4% paraformaldehyde in 0.1 M phosphate-buffered saline (PBS) at +5°C and washed for 2 hours in PBS. Fixed animals were then dissected into several pieces to make the tissue manageable for confocal microscopy. The dissected tissues were pre-incubated overnight in a blocking solution of 6% goat serum in PBS containing 0.02% Triton X-100 (PBT). The samples were then incubated for 48 hours at +5°C with primary antibodies in 6% goat serum at a final dilution 1:100. We used the rat monoclonal antibody (AbD Serotec Cat# MCA77G, RRID: AB_325003), which recognizes the alpha subunit of tubulin and specifically binds tyrosylated tubulin (Wehland & Willingham, 1983; Wehland et al., 1983). Following a series of PBS washes for 6-8 hours, the dissected tissues were incubated for 24 hours with secondary goat anti-rat IgG antibodies conjugated to Alexa Fluor 488 (Molecular Probes, Invitrogen, Cat# A11006, RRID: AB_141373) at a final dilution 1:50.

To label the muscle fibers, we used well-known marker phalloidin (Alexa Fluor 568 phalloidin from Molecular Probes), which binds to F-actin (Wulf et al., 1979). After washing in PBS following the secondary antibody treatment, the samples were incubated in phalloidin solution (in PBS) for 6 to 8 hours at +5° C, at a final dilution 1:80, and then washed in several PBS rinses for 8 hours.

To stain the nuclei, the tissue was mounted in VECTASHIELD Hard-Set Mounting Medium with DAPI (Cat# H-1500) or Antifade Mounting Medium with DAPI (Cat#H-2000). Some of the larger adult tissues were also mounted in Fluorescent Mounting Media (KPL) on glass microscope slides. The slides were viewed using a Nikon Research Microscope Eclipse E800 with Epi-fluorescence via standard TRITC and FITC filters and recorded using Nikon C1 confocal microscope.

### 2.2 Antibody specificity

Rat monoclonal antityrosinated alpha-tubulin antibody is raised against yeast tubulin, clone YL1/2, isotype IgG2a (Serotec Cat #MCA77G; RRID: AB_325003). The epitope recognized by this antibody has been extensively studied, including details about antibody specificity and relevant assays (Wehland et al., 1983; Wehland & Willingham, 1983). As reported by Wehland et al. (1983) this rat monoclonal antibody “reacts specifically with the tyrosylated form of brain alpha-tubulin from different species” (Wehland et al., 1983). We have successfully used this specific anti-α-tubulin antibody before on several species of ctenophores as well as hydrozoan *Aglantha* and siphonophore *Nanomia* to label their nervous systems (Norekian & Moroz, 2016, 2019a, 2019b, 2020a, 2020b; Norekian & Meech, 2020, 2026). We also tested the specificity of immunostaining by omitting either the primary or the secondary antibody from the procedure. In both cases, no labeling was detected.

## 3 RESULTS

### 3.1 Hydrozoan Diversity

We used 14 species of hydrozoan jellyfish from five different orders, all pelagic animals collected in the Pacific Ocean waters off the American Northwest (Fig. 1, 2). In the following paragraphs, we will describe the structure of the nervous and muscular systems, focusing primarily on the subumbrella region, which shows the greatest variability among jellyfish. Later in the Results, we will present a detailed description of our findings, including many additional details for each of the 14 hydrozoan species. The species are presented in the following order:

Subclass Hydroidolina

<u>Order Anthoathecata</u>

- family Corynidae

***Polyorchis penicillatus***
***Sarsia sp.***
- family Corymorphidae

***Euphysa sp.***
- family Bougainvilliidae

***Bougainvillia multitentaculata***
- family Pandeidae

***Stomotoca atra***
***Neoturris breviconis***
***Leuckartiara longicalcar***
<u>Order Leptothecata</u>

- family Aequoreidae

***Aequorea victoria***
- family Mitrocomidae

***Mitrocoma cellularia*** (also known as ***Halistaura cellularia***)
- family Campanulariidae

***Clytia gregaria*** (formerly known as ***Phialidium gregarium***)
- family Eirenidae

***Eutonina indicans***

Subclass Trachykinae

<u>Order Limnomedusae</u>

- family Olindiidae

***Eperetmus typus***
<u>Order Trachymedusae</u>

- family Rhopalonematidae

***Aglantha digitale***
<u>Order Narcomedusae</u>
- family Solmundaeginidae

***Solmundaegina nematophora*** (previously described as ***Aegina citrea***)
*Aegina* species at FHL has been separated off and given a new name, *Solmunaegina nematophora* (Lindsay et al, 2017).

**Figure 1.**
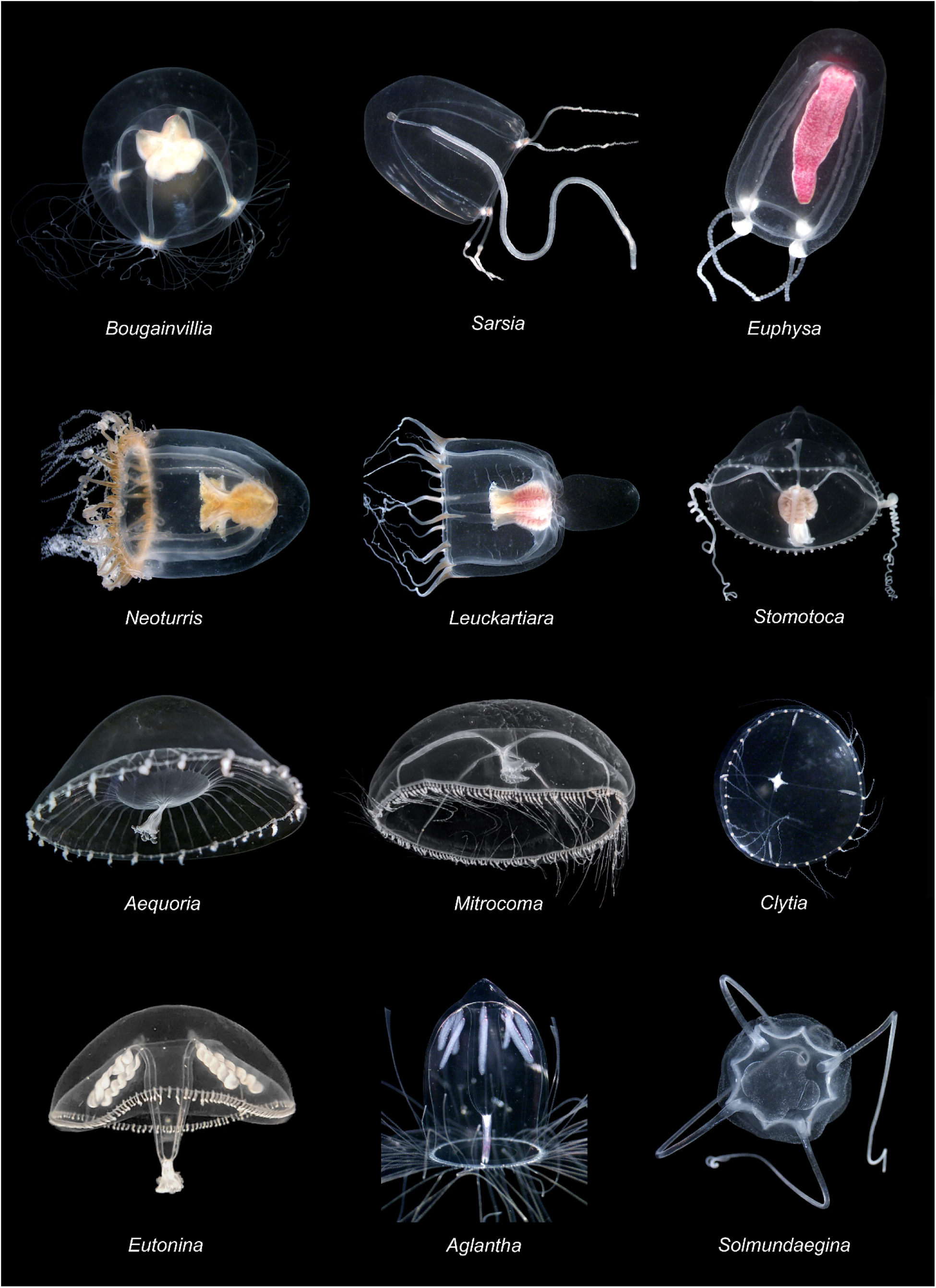
Photos of live mature animals representing different species used in this study.

**Figure 2.**
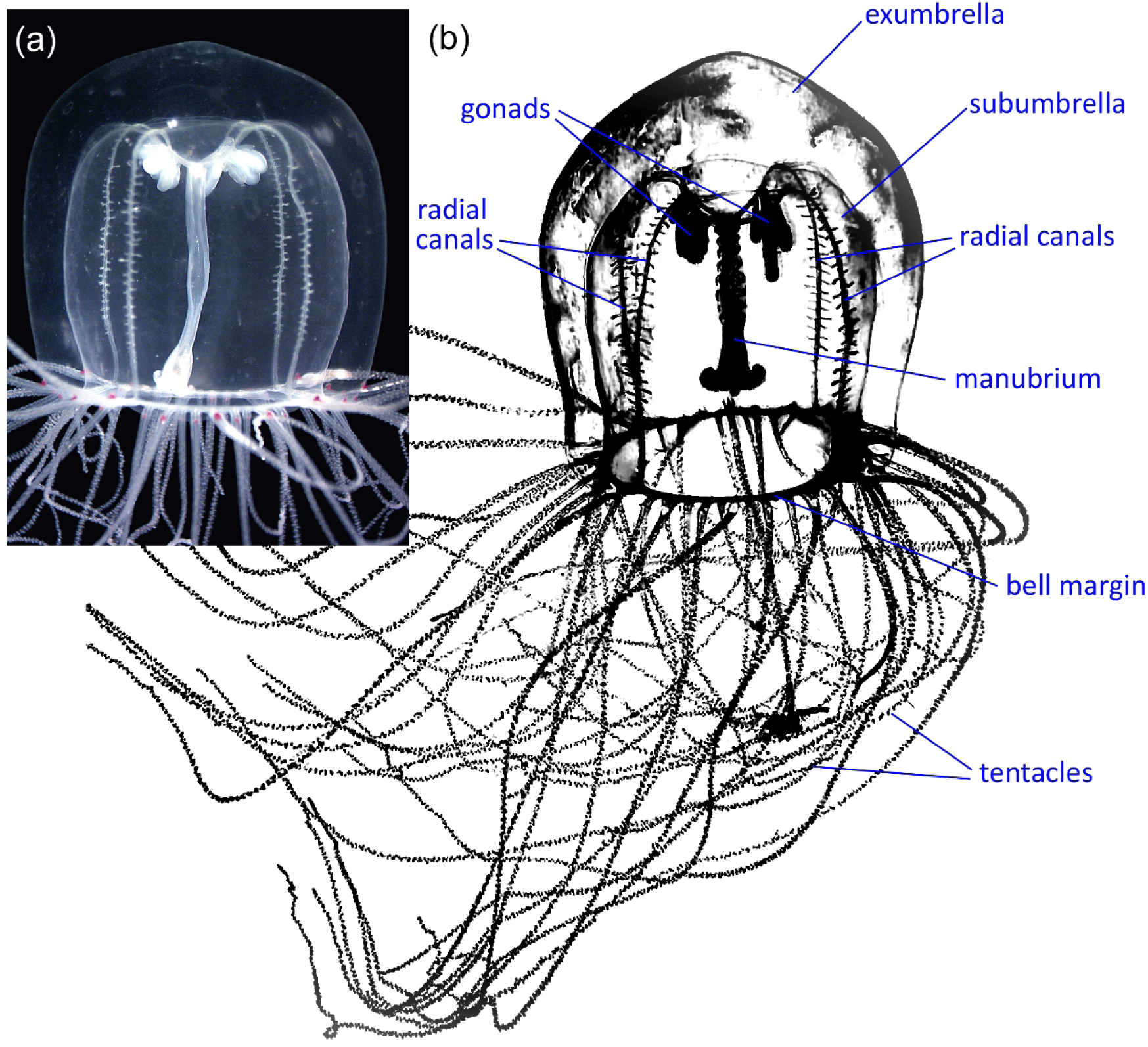
Schematic representation of a hydrozoan jellyfish and its main organs. **a** - Photo of live mature *Polyorchis*. **b** - Schematics based on a photograph of *Polyorchis penicillatus*.

### 3.2 Common elements in all hydrozoans

Hydrozoan medusae have a highly conservative nervous system structure, similar across all studied species. The nerve ring, which encircles the umbrella margin, is the central component of the nervous system in all hydrozoan jellyfish (Satterlie & Spencer, 1983; Satterlie, 2002). All species have a nerve ring consisting of outer and inner nerves (Fig. 3b; 5a; 6a, b, f; 7a; 9a, e; 11a, b, f; 13a; 15a, e; 17a, i; 20a-d; 22a, f; 25a; 26e, f; 27a, d; 29a; 30a, b). In some species, the outer and inner nerves are very closely located and difficult to separate, whereas in others the separation is very clear. All species have a myoepithelial layer of circular striated muscles in the subumbrella region responsible for swimming (Fig. 3g; 6d; 8c; 9f; 12d; 27b; 29e). The striated muscle fibers are nearly fused, forming a single sheet that functions as one unit. All species have radial canals running from the bell margin to the apical end through the entire subumbrella region (Fig. 2; 13a; 15a; 16d; 17a).

Most species have radial neural and muscular pathways that run from the bell margin to the apical end of the subumbrella along the radial canals (Table 1; Fig. 3a-e; 5a-g; 7b-f; 8a, b; 9a-d; 11a-c; 15a, b; 18a, b; 22a-c; 23a; 25a, b; 27d-f; 30a-d; 31a, b). One notable exception is *Neoturris* from the order Anthoathecata. In *Neoturris*, there are no radial nerves along the radial canals, and the entire subumbrella is covered by a neural network (Fig. 13b). However, the density of the neural network is higher along the radial canals (Fig. 14a-c). In *Aglantha* (order Trachymedusae), individual giant motor axons run from the bell margin to the apical end of the subumbrella instead of neural tracts (Fig. 29a-c). There are also no radial bundles of smooth muscle fibers in *Neoturris* and *Aglantha*. In *Neoturris*, radial muscle fibers are evenly distributed across the entire subumbrella next to the circular striated muscles, without forming radial bundles (Fig. 13b, c; 14b, c), while *Aglantha* completely lacks smooth radial muscles in the subumbrella region (some are present in the exumbrella). *Aglantha* does not do defensive “crumpling”; instead, it has a powerful escape response. Thus, all species that have thick bundles of smooth radial muscles along the radial canals have clearly defined neural pathways running along or even embedded within the smooth muscle bundles and crossing the entire subumbrella from the ring nerve to the apical end of the bell (Table 1).

**Table 1.** Distribution of Neural and Muscle elements across hydrozoan species.

|  |  |  |  |
| --- | --- | --- | --- |
| Radial pathways are present along the radial canals. They contain radial neural tracts and bundles of radial smooth muscle fibers. Outside of the radial pathways there are no neural elements and radial smooth muscles – only circular striated muscles. | Neural elements and radial smooth muscle fibers are present throughout the entire subumbrella region. They are spread all over the area. | Highly developed radial pathways, which include bundles of thick smooth radial muscle fibers and neural tracts. But there are also numerous neural processes crossing the entire subumbrella. | Instead of the radial neural tracts, there are motor giant axons. Lateral interneurons spread over the subumbrella region. No radial smooth muscle fibers in the subumbrella and no radial bundles of muscle fibers along the radial canals under axons. |
| *Polyorchis*<br>*Sarsia*<br>*Euphysa*<br>*Stomotoca*<br>*Bougainvillia*<br>*Leuckartiara* – does not have a tight bundle of thick radial smooth muscles as others, but a very wide lane that contains thin smooth radial muscle fibers. | *Neoturris* – has fully developed polygonal neural network.<br>*Aequorea*, *Mitrocoma*, *Clytia*, *Eutonina* – all have neural networks spread throughout the subumbrella. Not very dense radial neural tracts might be present along the radial canals. | *Eperetmus* – looks similar to the first group, but has long neural processes across the subumbrella. No smooth radial muscles outside of the radial pathways.<br>*Solmundaegina* – very different. Radial pathways (tentacular nerves) lead to the tentacles located not at the bell margin but higher on the bell. There are neural cells and processes across the subumbrella. | *Aglantha* – very different and unique. Their nervous system in subumbrella is dedicated to the control of swim striated muscles, while in other species it looks like it controls smooth radial muscles. |

In four species from the order Leptothecata – *Aequorea, Mitrocoma, Eutonina* and *Clytia* – radial smooth muscle fibers are evenly distributed across the entire subumbrella without forming tight radial muscle bundles, at least in the lower and central parts of the subumbrella. In *Aequorea*, radial smooth muscle bundles begin to form near the apical end of the bell and continue into the peduncle (Fig. 19b-e). In three species – *Aequorea, Eutonina* and *Clytia* – radial neural tracts are clearly present in the form of a denser neural network along the radial canals but are not organized into fully formed nerves (Fig. 18a, b; 22a-c; 23a; 25a-c). In *Mitrocoma*, tubulin IR brightly labels numerous small non-neuronal cells on the surface of the radial canal, completely masking any possible radial neural tracts.

*Eperetmus*, the single species studied in the order Limnomedusae, has smooth muscle bundles and dense neural pathways running along the radial canals (Fig. 27d-f; 28a, b). Interestingly, upon reaching the firmly attached subumbrella gonad, each muscle bundle and neural pathway bifurcates to run around the gonad on both sides (Fig. 27e).

### 3.3 Neural networks in subumbrella

In addition to relatively conservative neural elements among all studied hydrozoan species, which include the nerve ring and radial neural pathways, there is tremendous variety and some remarkable differences in the distribution of neural elements in the subumbrella region. *Polyorchis, Sarsia, Stomotoca, Euphysa*, and *Bougainvillia* (five species from the order Anthoathecata) lack neural fibers or neural cells in the subumbrella regions between the radial neural pathways (Fig. 3b, c; 4f; 5b; 8b; 9b, e; 11a, b, e). The myoepithelial layer of circular striated muscles and the neighboring layers in the subumbrella do not have innervation in those areas. Importantly, those areas also lack radial smooth muscle fibers (Fig. 3a; 5d; 6c; 7b; 9b, e; 11a, b). All radial smooth muscle fibers are confined to the radial smooth muscle bundles along the radial canals. And all these species have clearly defined neural pathways running along those smooth muscle bundles (Fig. 3b-f; 5b, c, e-g; 7c-g; 8a, b; 9a-d; 11a-d). It seems that activity in the myoepithelial layer of circular striated muscles (swim muscles) is triggered by the ring nerve, but the signal spreads through the muscle layer without any neural network involvement (see Mackie & Passano 1968; Mackie, 1965, 1970; Satterlie & Spencer, 1983).

The only clear exception among the 7 studied species from the order Anthoathecata is *Neoturris*. In *Neoturris*, the entire subumbrella region is covered by a fully formed polygonal neural network (Fig. 13a-f; 14a-c). The subumbrella region also contains evenly distributed radial smooth muscle fibers (Fig. 13b, c; 14b, c). There are no radial smooth muscle bundles or radial neural pathways – no concentration of the elements; all are evenly distributed. *Leuckartiara*, a close relative of *Neoturris* and a member of the same family, Pandeidae, has a different distribution of muscle and neural networks. Large areas of the subumbrella between the radial canals lack radial muscle fibers and have no neural processes or neural cells (Fig. 15a, e). Only the circular striated muscle layer is present. A wide tract (three times wider than the radial canal) of muscle layer lies next to each radial canal and, in addition to striated muscles, contains radial smooth muscle fibers (Fig. 15a-d). It does not resemble the thick, tight bundle of radial smooth muscle found in other species but rather resembles the subumbrella of *Neoturris*. The polygonal neural network is located only within that wide radial tract (Fig. 15a-d). The neural network, in the form of a wide band, also runs next to the ring nerve and at the apical end of the subumbrella, connecting neighboring radial tracts (Fig. 15e, f; 16a-c). In other words, *Leuckartiara* resembles *Neoturris* but has four large voids between radial canals that lack radial smooth muscle fibers and the neural network.

*Stomotoca* belongs to the same family, Pandeidae, as *Leuckartiara* and *Neoturris*. It also has a very similar polygonal neural network in the subumbrella region (Fig. 11a-d). However, there is a key difference. As mentioned earlier, *Neoturris* has the entire subumbrella surface covered with such a polygonal network, whereas *Leuckartiara* has a polygonal neural network but also four large voids in the middle of the subumbrella between the radial pathways, with no neural network present. In*Stomotoca*, there is also a polygonal network, but it covers only a relatively narrow strip along the radial canals halfway through the bell, extending to the peduncle base (Fig. 11a-c; 12a). Most of the subumbrella surface lacks the network (peduncle has neuronal elements, but not a polygonal network). Thus, there is a clear transition from the *Neoturris* scheme, with a neural network spread across the entire subumbrella, through *Leuckartiara*, to *Stomotoca*, with its typical organization of other Anthoathecata species, without any subumbrella neural networks. This transition is observed in three related species belonging to the same family.

Thus, the studied species from the order Anthoathecata show a clear tendency to lose neural networks and radial smooth muscle fibers in the subumbrella region, with neural elements and smooth muscle fibers concentrated only in the radial pathways along the radial canals (Table 1).

Four studied species from the order Leptothecata - *Aequorea, Mitrocoma, Clytia*, and *Eutonina* - all have radial smooth muscle fibers distributed throughout the entire subumbrella region (Fig. 18a-f; 21c, d; 23a-d; 25b-f). They also have neural networks in the subumbrella area between the radial neural tracts. These networks consist of irregular polygonal units of varying sizes and are significantly elongated radially (Fig. 18c-f; 21a-d; 23b-d; 25d-f). In much younger adults of the *Aequorea* and *Eutonina* species, the subumbrella neurons do not form a fully developed polygonal network but instead represent a meshwork of individual neural cells with long branching processes (Fig. 18c, d; 25d). Some neighboring neurons overlap with their processes, suggesting possible contact, while others are more isolated. In larger adults reaching their medium and maximum sizes, the neural network is fully formed, and neural cell bodies are embedded in the threads of the network (Fig. 18e, f; 25e-f). All four species also have a narrow stretch of a denser network of neurons and neurites attached to the ring nerve in the margin area (Fig. 18a; 21a, b; 22d-f; 23a; 25b). Thus, unlike most Anthoathecata species, all four studied species from the order Leptothecata have a developed subumbrella neural network, as well as radial smooth muscles spread over the entire subumbrella region.

*Eperetmus*, from the order Limnomedusae, has a very wide band of circular smooth muscle fibers in the subumbrella region next to the ring nerve (Fig. 27a-d). This circular smooth muscle band is densely innervated (Fig. 27a, c, d). There are no individual radial smooth muscle fibers in the subumbrella between the radial bundles – only the myoepithelial layer of circular striated muscles (Fig. 27b; 28a-d). However, numerous neural processes cross those areas (Fig. 28a-d). This is the only exception (along with *Aglantha*) to the general correlation that neural elements are found only where smooth muscles are located (Table 1). However, the neural processes that cross the area do not branch much and appear to be passing through.

*Aglantha*, from the order Trachymedusae, is highly distinctive. First, there are no radial smooth muscle fibers in the subumbrella region at all – neither separate radial fibers throughout the subumbrella nor bundles of smooth muscle along the radial canals beneath the giant motor axons (Fig. 29a-e; Norekian and Moroz, 2020b). Second, the only neural elements present in the subumbrella region are *Aglantha*’s unique giant motor axons and lateral interneurons (Fig. 29a-d; Norekian and Moroz, 2020b). All these neural elements specifically target the circular striated swim muscles (Donaldson et al., 1980; Kerfoot et al., 1985; Mackie & Meech, 1985, 2000; Roberts & Mackie, 1980; Singla, 1978b; Weber et al., 1982). Apparently, the speed of signal propagation is essential for *Aglantha*, and therefore the neural system gets involved.

The structure of the nervous and muscular systems in *Solmundaegina*, from the order Narcomedusae, is also unique among hydrozoans. Primarily, because of four very thick tentacular nerves that run from the nerve ring to the base of each tentacle and subsequent dense innervation of the four muscular tentacles (Fig. 30a-h). This innervation reflects the unusual for hydrozoans use of tentacles as a mechanical instrument to grab a large prey, rather than immobilization by numerous nematocysts. The neural network and longitudinal smooth muscle fibers are concentrated mostly on one inward side of the tentacles responsible for powerful bending and grabbing movements (Fig. 30e-h). The dense innervation of the mouth area also reflects the processing of large, active prey (Fig. 31e-g).

The circular striated muscle area is relatively narrow because it is squeezed between the bell margin and the large digestive pouches unique to *Solmundaegina* (Fig. 31a). As an active predator, *Solmundaegina* is a very fast swimmer, second only to *Aglantha*. The network of neurites, which originates from the tentacular nerve and extends to the band of striated circular muscles, runs mostly in parallel with the swim muscle fibers and may target those muscles, facilitating the propagation of signals to the swim musculature (Fig. 31b-d). While in *Aglantha* the role of motor giant axons and lateral interneurons in swimming has been confirmed electrophysiologically, there is no such data for *Solmundaegina*.

### 3.4 Radial canals and Peduncle

In addition to the neural pathways along the radial canals and the neural networks in the subumbrella muscle layer, neural elements are also present in the walls of the radial canals themselves (Table 2). These may include a few large, tripolar neurons with long processes running the length of each radial canal, as in *Euphysa* and *Aglantha* – they look very similar in those two species (Fig. 8d-f; 29f). Alternatively, they may consist of several neural processes formed by small bipolar neurons, also longitudinally oriented, as in *Mitrocoma* (Fig. 21f). They may also form a dense meshwork of numerous small multipolar neurons, as in *Stomotoca*, *Neoturris* and *Leuckartiara* – very similar among these three closely related species (Fig. 12b, c; 14d, e; 16d-f). Finally, they may include a fully formed neural network with thick threads covering the walls of all radial canals and their diverticula, as in *Polyorchis* (Fig. 4a-f).

**Table 2.**
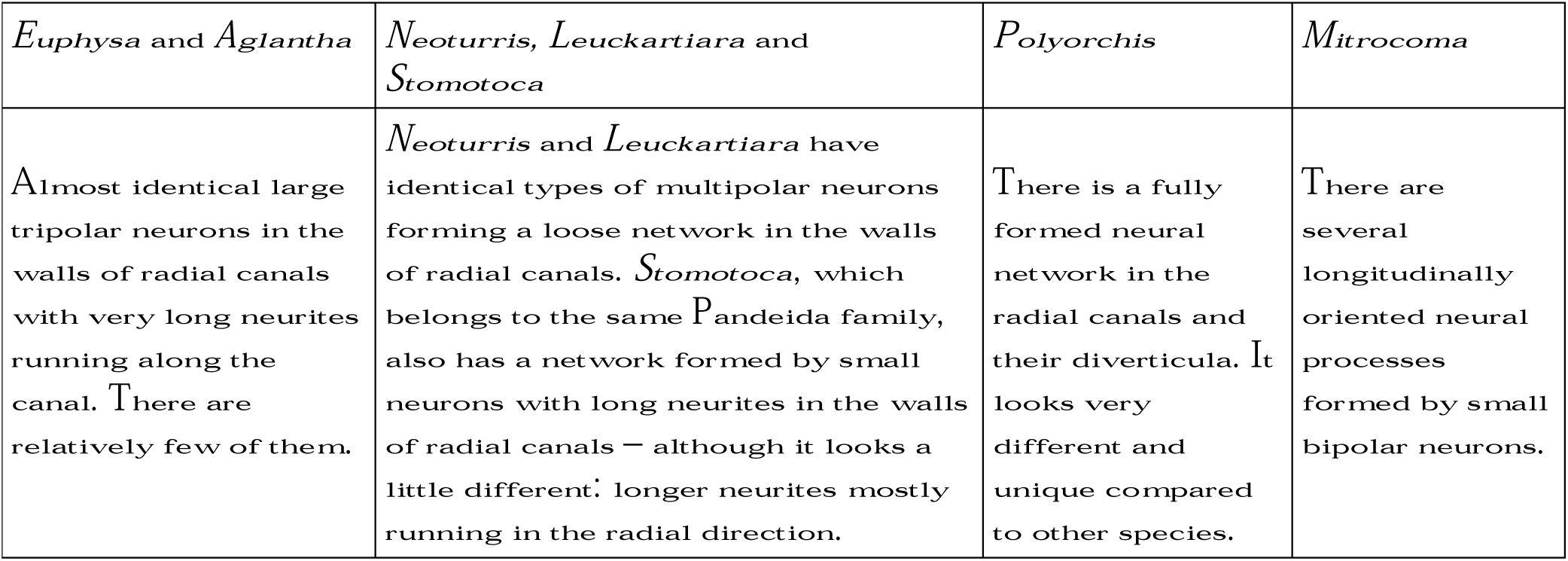
Neurons in the walls of the radial canals.

At the apical end of the bell, above the proper subumbrella area, peduncles in all studied species always had a clear innervation, either a fully formed polygonal neural network, as in *Aequorea*, *Mitrocoma* and *Eutonina* (19d-f; 21e; 26a, b), or a mesh of multipolar neurons with long, branching neurites connecting neighboring cells, as in *Stomotoca* (Fig. 12f-i). This innervation was present in species that have subumbrella neural networks, such as *Aequorea*, *Mitrocoma*, and *Eutonina*, and in species that lack subumbrella neural elements outside the radial neural pathways, as in *Stomotoca*.

### 3.5 Detailed description of the studied hydrozoan species

#### *Polyorchis penicillatus* (order Anthoathecata, family Corynidae)

There are four radial canals, each branching into several lateral diverticula. Tubular gonads hang from the short peduncle, as well as a long, thin manubrium. There are many thin and long tentacles tightly packed on the margin of the bell, with ocelli surrounded by red pigment at the base of the tentacles.

Four thick bundles of smooth radial muscles run along the radial canals and cross the circular striated muscle sheet in the subumbrella (Fig. 3a). There are also four radial neural tracts in the same area, traveling along the radial canals with the smooth radial muscle bundles (Fig. 3b; see also Lin, Gallin & Spencer, 2001). The neural tracts originate at the nerve ring, which is brightly labeled by tubulin AB and encircles the umbrella at the bell margin (Fig. 3b). Each neural tract includes two thick nerves consisting of several neurites that run on both sides of the radial smooth muscle bundle from the ring nerve to the apical end of the bell (Fig. 3c-e). Many short perpendicular branches connect the two longitudinal nerves along the way. Upon entering a short, wide peduncle at the top of the subumbrella, the two nerves on both sides of each radial canal split into separate neurites and terminate without proceeding further (Fig. 3d). The nerves and neurites appear to “hug” the band of smooth radial muscles that runs along the radial canals, suggesting that this band is their target.

**Figure 3.**
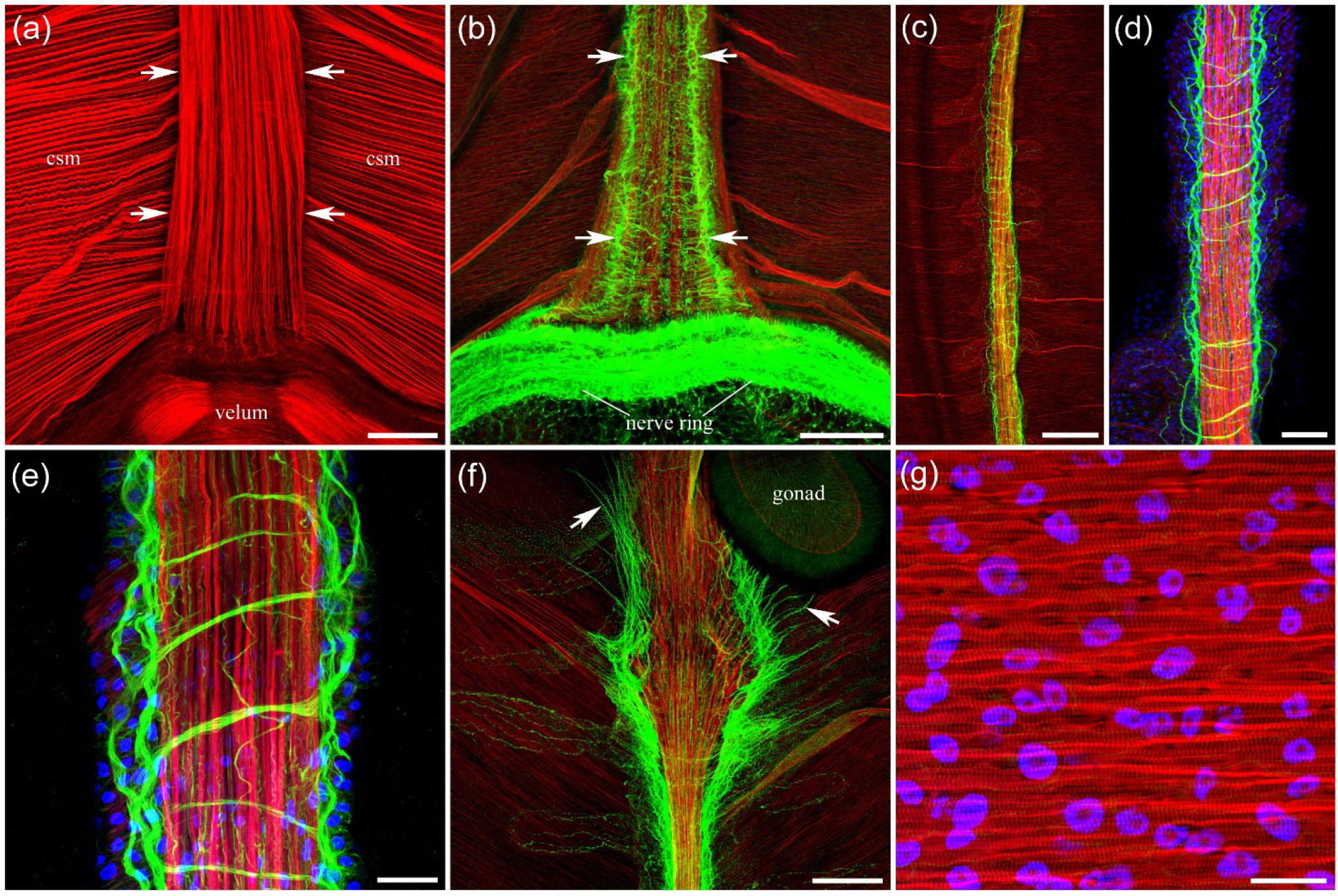
The structure of the *Polyorchis* neural and muscle systems in the subumbrella. Tubulin antibody is green, phalloidin is red, and nuclear DAPI staining is blue – here and in all subsequent figures. The orientation of all images here and in all subsequent figures follows the same pattern: the margin of the bell is down, while the apical end of the bell is up. **a** - Phalloidin labels the thick band of radial smooth muscle fibers (arrows) running along the radial canals, the myoepithelial layer of circular striated muscles (*csm*) controlling swimming, and muscles in the velum. **b** - Preparation with similar orientation and magnification as in (a), but with tubulin AB brightly labeling the nerve ring at the margin of the bell. There is also a radial neural tract (arrows) originating from the nerve ring and traveling with the smooth radial muscle bundle. **c, d, e** - Each radial neural tract embedded with the radial smooth muscle bundle runs across the entire subumbrella area from the nerve ring to the apical end of the bell and includes two thick nerves, each consisting of several smaller neurites and running on both sides of the radial smooth muscle bundle. Many short perpendicular branches also connect the nerves along the way. **f** - Upon entering a wide, short peduncle at the top of the subumbrella, these two nerves on both sides of each radial muscle bundle split into separate neurites (arrows) and terminate their advance. **g** - Circular striated muscles in the subumbrella controlling swimming, with their striation clearly visible at higher magnification. Scale bars: **a, b** - 100 µm; **c** - 200 µm; **d** - 50 µm; **e** - 20 µm; **f** - 200 µm; **g** - 15 µm.

A neural network also exists in the walls of the radial canals (Fig. 4; see also Lin, Gallin & Spencer, 2001). These radial canal neurites and individual neurons are located below the radial neural tracts and smooth muscles – in different optical focal planes. When the radial canal and the band of radial smooth muscle fibers shift relative to each other, the neural network in the wall of the main radial canal becomes clearly visible (Fig. 4a). These neurons and neurites extend from the main trunk of each radial canal into the body wall of diverticula, where they are clearly visible and unobstructed by the radial neural tracts (Fig. 4b). The radial canal network is fully formed, with thick threads extending mostly in the longitudinal direction and bipolar or tripolar neuronal cell bodies incorporated within the threads (Fig. 4c). Small, young animals have very short diverticula and small neural networks (Fig. 4d-e), whereas older, larger animals have very long diverticula, each containing extended neural networks throughout their entire length (Fig. 4f).

**Figure 4.**
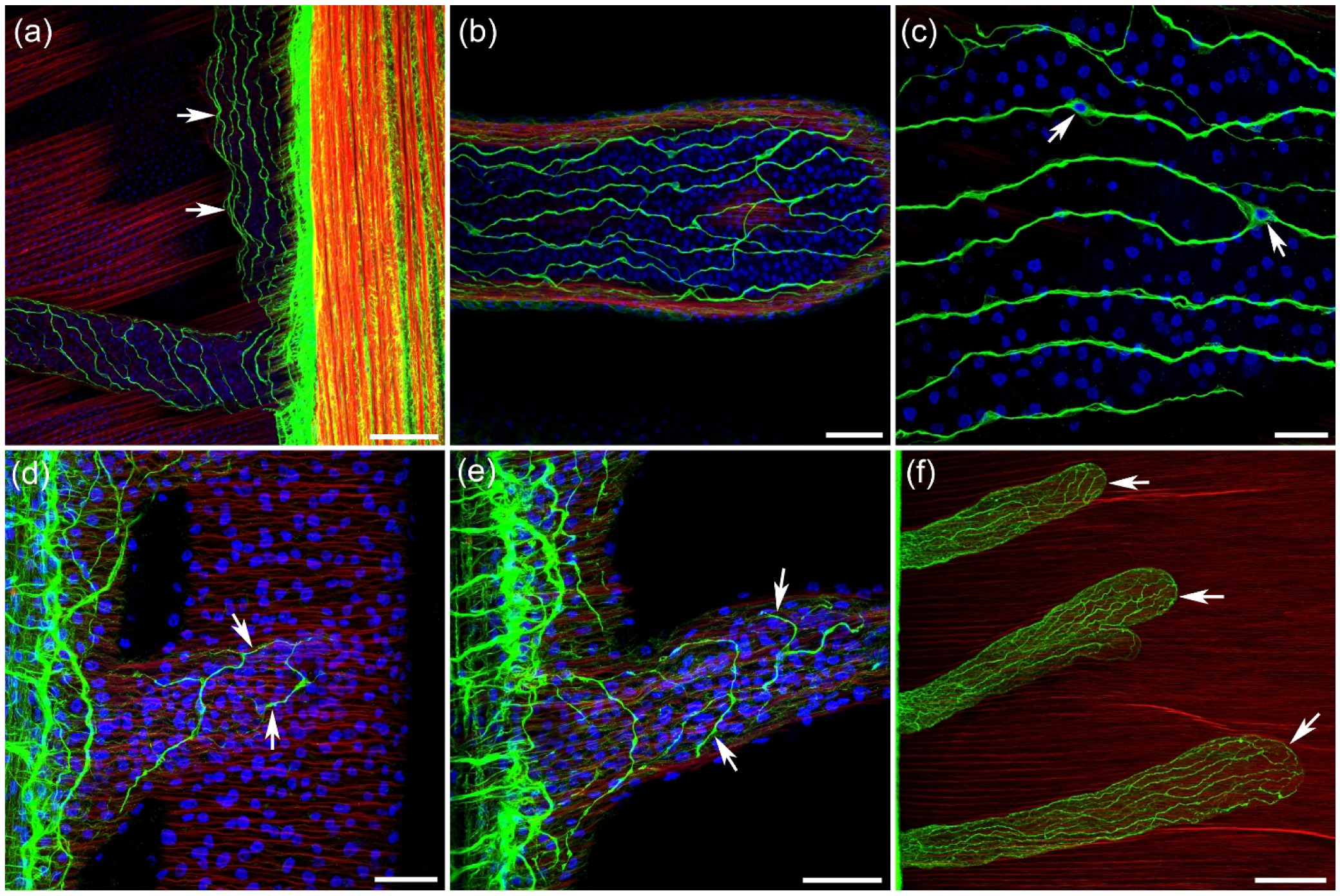
Innervation of the radial canals and their diverticula in *Polyorchis*. **a** - When the radial canal and the bundle of radial smooth muscles are shifted relative to each other, the neural network (arrows) in the wall of the main radial canal becomes visible. **b** - These neurons and neurites extend from the main trunk of each radial canal into the body wall of the diverticula, where they are unobstructed by the radial neural tracts. **c** - Bipolar and tripolar neurons (arrows) that form the radial canal network are clearly visible at higher magnification with DAPI staining. **d, e, f** - Small, young animals have very short diverticula with small neural networks (arrows in d, e), whereas older, larger animals have very long diverticula (f) that contain developed neural networks throughout their entire length (arrows). Scale bars: **a** - 100 µm; **b** - 50 µm; **c** - 20 µm; **d, e** - 30 µm; **f** - 200 µm.

There are no neurons or neurites in the swim striated muscle layer outside the radial neural tracts. There are also no smooth radial muscle fibers among the swim circular striated muscles outside the tracts.

#### *Sarsia sp.* (order Anthoathecata, family Corynidae)

There are four radial canals. Four tentacles are attached to the bell margin – each at the end of a radial canal. Each tentacle has prominent ocelli at its base. The manubrium is thin and very long – extending well beyond the bell and resembling another tentacle. Gonads are located on the manubrium.

The nerve ring, labeled with tubulin AB, encircles the bell margin (Fig. 5a). Along the radial canals, a band of smooth radial muscles crosses the circular striated muscle layer (Fig. 5a, d). This radial pathway, which includes the band of smooth radial muscles, also contains a neural tract that runs from the ring nerve at the margin to the manubrium at the apical end of the bell (Fig. 5b, c). The radial neural tract consists mostly of thick, longitudinally oriented neurites running radially along the tract (Fig. 5e-g). It usually contains 3-5 large axons, plus a few smaller neurites. Short branches also cross the tract, connecting neighboring longitudinal neurites. Bipolar and tripolar neuronal cell bodies that form those axons are clearly visible at high magnification with DAPI staining (Fig. 5g).

**Figure 5.**
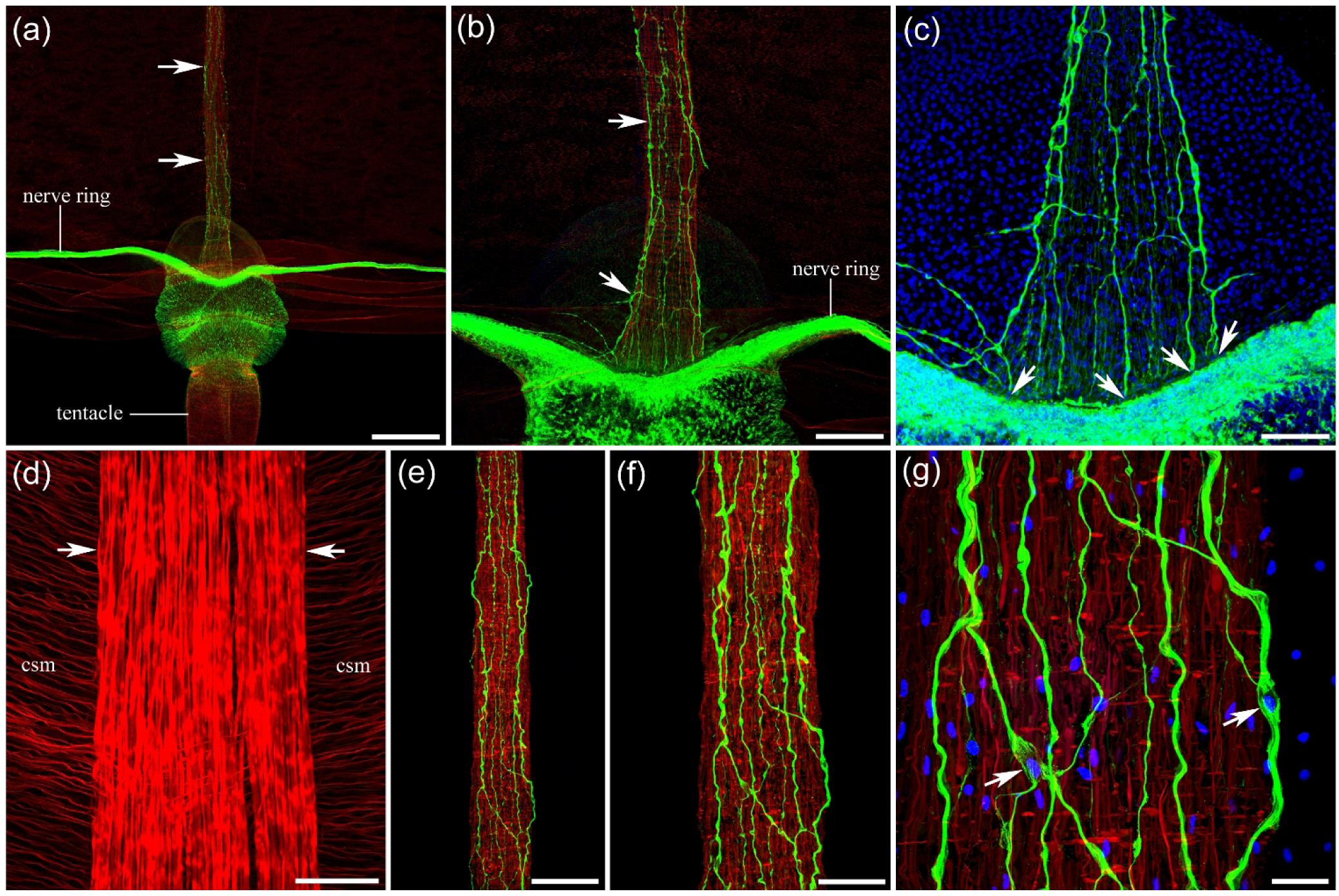
Neural and muscle systems in the subumbrella of *Sarsia*. **a,b** - Each of the four radial smooth muscle bundles and radial neural tracts (arrows) originates at the base of the four tentacles. **c** - Neurites that form the radial neural tract connect to the nerve ring (arrows). **d** - Phalloidin labels a band of smooth radial muscles (arrows) crossing the swim circular striated muscle (*csm*) layer. **e, f, g** - The band of smooth radial muscles and embedded neural tract cross the umbrella from the nerve ring to the apical end of the bell. The tract contains 4-5 large longitudinally oriented axons, plus a few smaller neurites. Neural cell bodies that give rise to these axons are identified by DAPI staining (arrows). Scale bars: **a** - 500 µm; **b** - 200 µm; **c** - 100 µm; **d** - 40 µm; **e** - 200 µm; **f** - 100 µm; **g** - 25 µm.

The nerve ring is clearly divided into the outer and inner nerve rings (Fig. 6a, b). A narrow band of smooth circular muscle fibers lies around the margin, next to the nerve ring (Fig. 6c). There are no neurons or neurites in the swim circular striated muscle layer outside the narrow radial neural tracts along the radial canals. There are also no smooth radial muscle fibers among the swim circular striated muscles outside those radial tracts (Fig. 6d). However, smooth radial muscle fibers are clearly present among the circular striated muscle fibers of the velum (Fig. 6e). These data raise the same questions – does the radial neural tract innervate and activate the smooth radial muscles, and is the lack of neural elements in the rest of the subumbrella directly connected to the absence of smooth radial muscles in that region?

**Figure 6.**
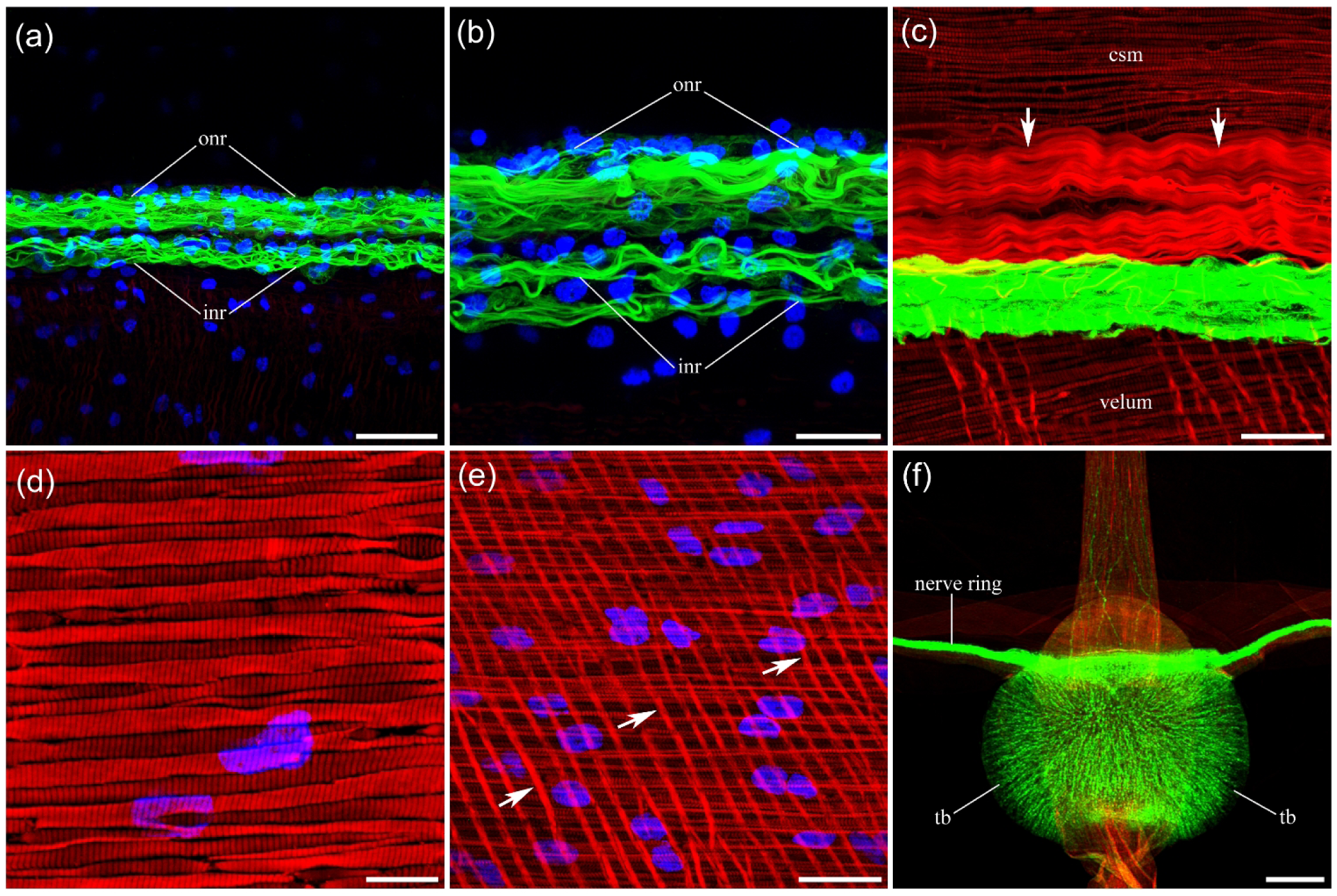
Nerve ring and muscle types in *Sarsia*. **a,b** - Labeled with tubulin AB nerve ring encircles the bell margin, with the outer nerve ring (*onr*) and inner nerve ring (*inr*) clearly separated. **c** - A narrow band of smooth circular muscle fibers (arrows) is located around the margin, next to the nerve ring. **d** - Circular striated muscles (*csm*) at higher magnification, labeled with phalloidin. **e** - The velum contains striated circular muscle fibers and smooth radial muscles (arrows). **f** - The round-shaped tentacle base (*tb*) contains numerous radial filaments labeled with tubulin AB, which are probably not part of the nervous system. Scale bars: **a** - 40 µm; **b** - 20 µm; **c** - 30 µm; **d** - 7 µm; **e** - 20 µm; **f** - 250 µm.

#### *Euphysa sp.* (order Anthoathecata, family Corymorphidae)

There are four radial canals and four thick, relatively short tentacles. The manubrium resembles a long, thick, red sausage attached to the apical end of the subumbrella.

The nerve ring is brightly labeled with the tubulin antibody and clearly divided into inner and outer nerve rings (Fig. 7a). A tight bundle of thick radial smooth muscle fibers crosses the circular striated muscle layer within each radial canal (Fig. 7b). This pathway of radial muscle fibers also contains a narrow neural tract originating from the nerve ring and extending toward the apical end of the bell (Fig. 7c). Each radial neural tract consists of two thick nerves, sometimes three, and several smaller neurites running longitudinally along the radial muscle fibers (Fig. 7d, e). Closer to the apical end of the umbrella, the radial neural tract narrows but produces short lateral branches that extend into the neighboring tissue (Fig. 7f, g). The longitudinal neurites that form the radial nerves arise from bipolar neurons, which are clearly visible at higher magnification with DAPI staining (Fig. 8a, b).

**Figure 7.**
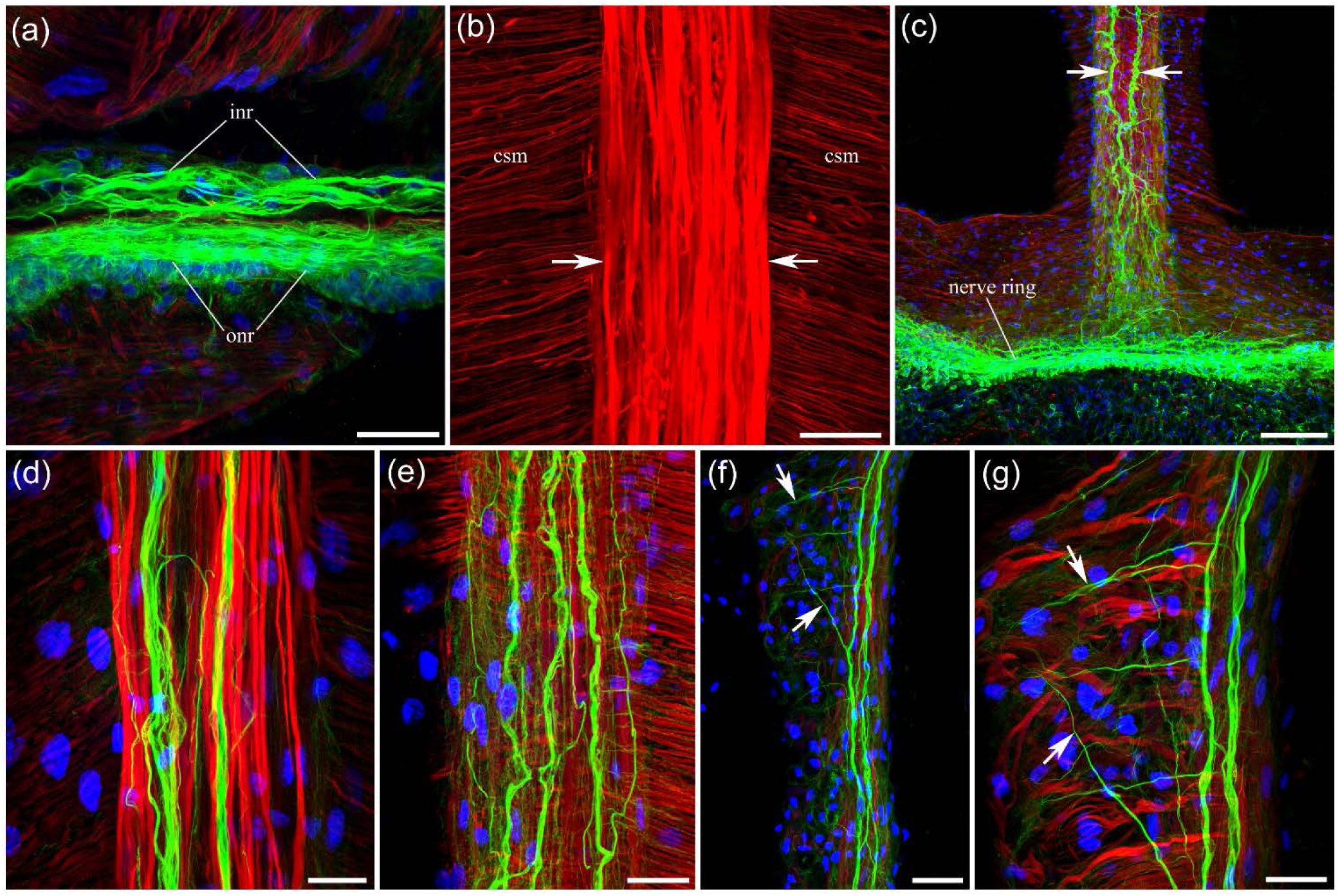
Neural and muscle systems in *Euphysa*. **a** - The nerve ring is brightly labeled with tubulin AB and clearly divided into the inner nerve ring (*inr*) and the outer nerve ring (*onr*). **b** - A bundle of radial smooth muscle fibers (arrows) crosses the circular striated muscle (*csm*) layer along each radial canal. **c** - This region of radial smooth muscle fibers contains a narrow neural tract (arrows) originating from the nerve ring and extending toward the apical end of the bell. **d, e** - Each radial neural tract consists of two thick nerves, sometimes transforming into three, and several smaller neurites running longitudinally along the radial muscle fibers. **f, g** - Closer to the apical end of the umbrella, the radial neural tract narrows but produces more lateral branches into the neighboring tissue (arrows). Scale bars: **a, b** - 40 µm; **c** - 100 µm; **d** - 20 µm; **e** - 30 µm; **f** - 50 µm; **g** - 30 µm.

There are no neurons or neurites outside the radial neural tracts, in the subumbrella within the swim circular striated muscle layer. There are also no smooth radial muscle fibers among the swim circular striated muscles outside those radial pathways (Fig. 8c).

**Figure 8.**
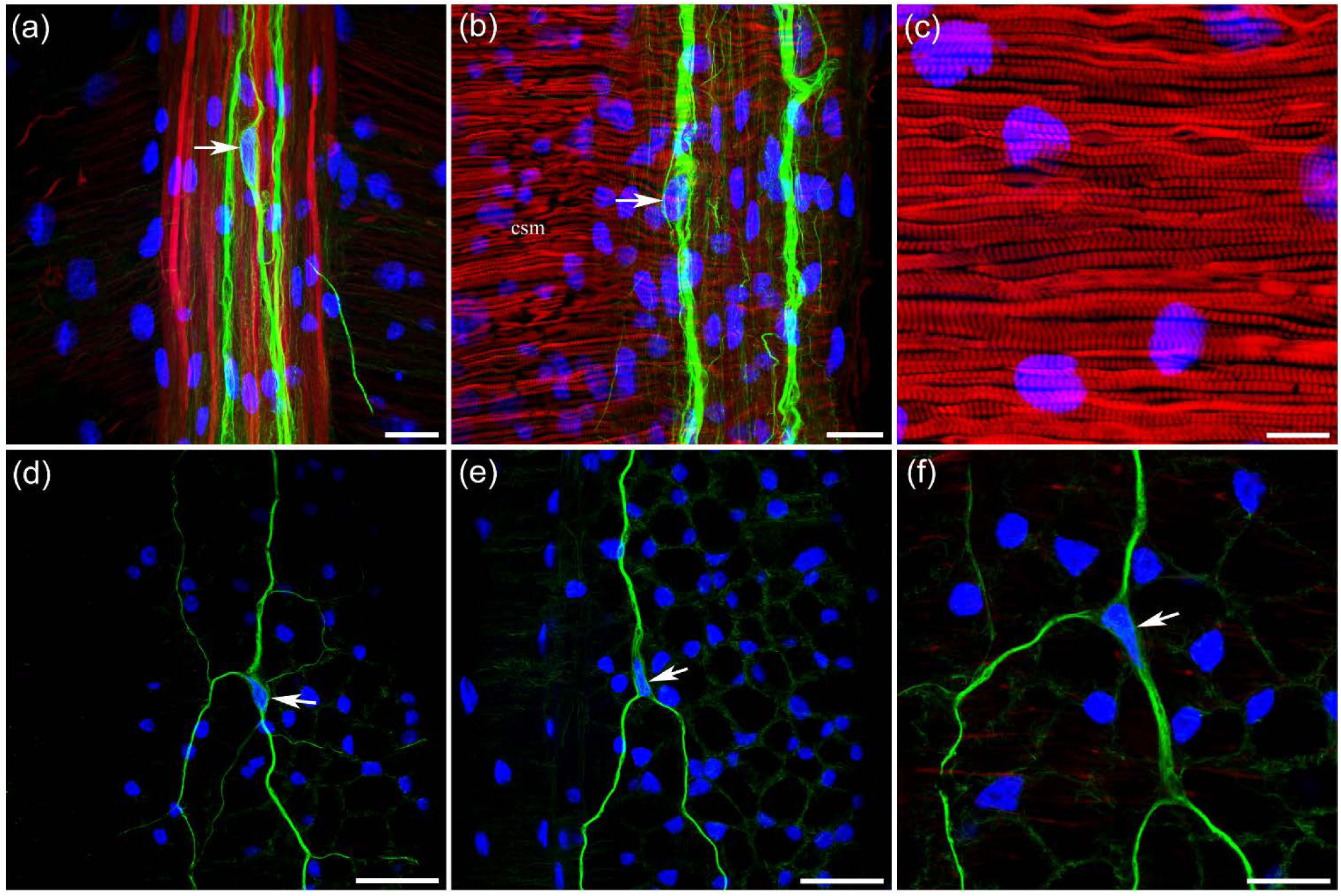
Neural elements in the *Euphysa* radial tract and radial canals. **a,b** - Bipolar neurons (arrows) along the radial neural tract form some of the longitudinal axons. **c** - The circular striated muscle layer labeled with phalloidin. **d, e, f** - Below the radial neural tract and smooth radial muscle bundle, in the walls of the radial canals (in different focal planes), a few large, separate tripolar neurons (arrows) produce very long neurites that travel longitudinally along the radial canals. Scale bars: **a, b** - 20 µm; **c** - 10 µm; **d, e** - 40 µm; **f** - 20 µm.

Below the radial neural tracts and smooth radial muscle bundles, within the walls of the radial canals, there are large tripolar neurons with very long neurites that run longitudinally along the radial canals (Fig. 8d-f). They closely resemble the tripolar neurons in the walls of the radial canals of *Aglantha digitale* (Norekian and Moroz, 2020b).

### *Bougainvillia multitentaculata* (order Anthoathecata, family Bougainvilliidae)

The umbrella is short and round, typically about 1 cm in diameter. There are four radial canals. The tentacle arrangement is highly distinctive compared with other species. There are only four large tentacle bulbs, which are brightly stained with tubulin AB (Fig. 9a). Each tentacle bulb bears many small tentacles. A relatively thin nerve ring, brightly labeled with tubulin AB, encircles the bell margin (Fig. 9a). At higher magnification, numerous individual neurites are visible within the nerve ring (Fig. 9e).

**Figure 9.**
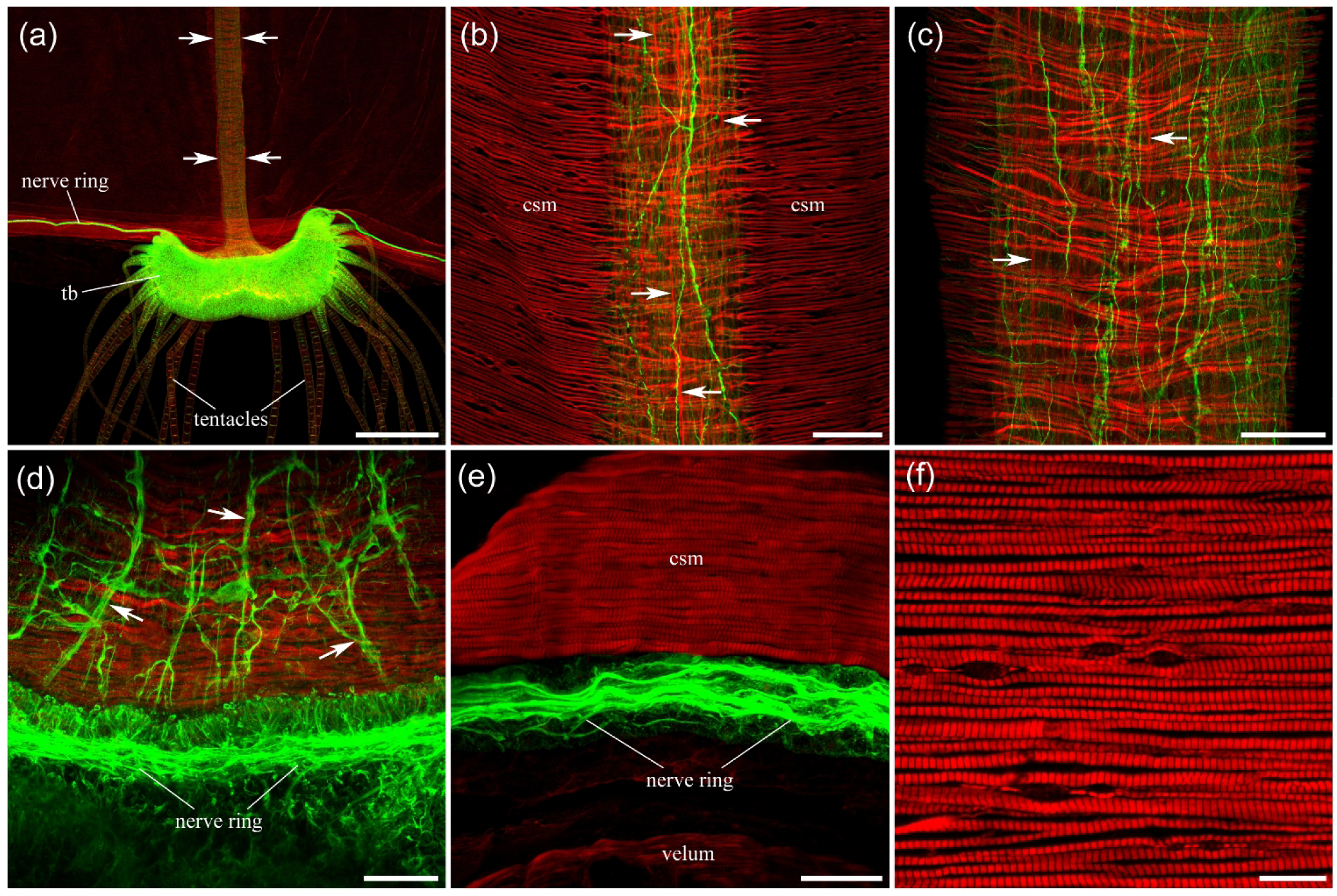
The structure of the *Bougainvillia* neural and muscle systems in subumbrella. **a** - A large tentacle bulb (*tb*), which bears a group of thin tentacles, is labeled with tubulin AB. A radial neural and muscular pathway (outlined by arrows) runs from the marginal nerve ring near the tentacle bulb toward the apical end of the umbrella. **b, c** - The radial pathway contains smooth radial muscle fibers (some shown by arrows) and several mostly longitudinal thin neurites labeled with tubulin AB (green). **d** - The radial neural pathway (some neurites indicated by arrows) originates at the nerve ring in the bell margin. **e** - The nerve ring is brightly labeled with tubulin AB and consists of several individual axons clearly visible at higher resolution. **f** - Circular striated muscles at higher magnification, labeled with phalloidin. Abbreviations: *tb* – tentacle bulb, *csm* – circular striated muscles. Scale bars: **a** - 500 µm; **b** - 50 µm; **c** - 40 µm; **d** - 30 µm; **e** - 20 µm; **f** - 10 µm.

Along each of the four radial canals, a band of smooth radial muscles crosses the circular striated muscle layer from the tentacle bulb at the margin to the apical end of the bell (Fig. 9a). This pathway of smooth radial muscles also contains a neural tract composed of several longitudinally oriented neural processes (Fig. 9b-c). Although most neurites are longitudinally oriented, some small processes cross the tract to connect neighboring longitudinal neurites. This neural pathway originates from the nerve ring (Fig. 9d).

There are no neurons or neurites in the swim circular striated muscle layer outside the narrow radial neural tracts along the radial canals. There are also no smooth radial muscle fibers among the swim circular striated muscles outside those radial tracts (Fig. 9b, e, f).

The mouth structure in *Bougainvillia* is highly distinctive. A wide peduncle supports a short manubrium, which bears several branching oral tentacles (Fig. 10a-b). Each branch typically bifurcates 3 or 4 times along its length (Fig. 10b-c). At the tip of each branch, a cluster of nematocytes is present (Fig. 10d-e). Each nematocyte has a single nucleus stained with DAPI, a cell body labeled with tubulin AB, and a ring of phalloidin-labeled stereocilia surrounding a single sensory cilium (cnidocil) stained with tubulin AB (Fig. 10e). All oral branches contain a dense neural network labeled with tubulin AB, with neurites oriented mostly longitudinally along the entire length of each branch, while some processes cross them perpendicularly and connect neighboring longitudinal neurites (Fig. 10d).

**Figure 10.**
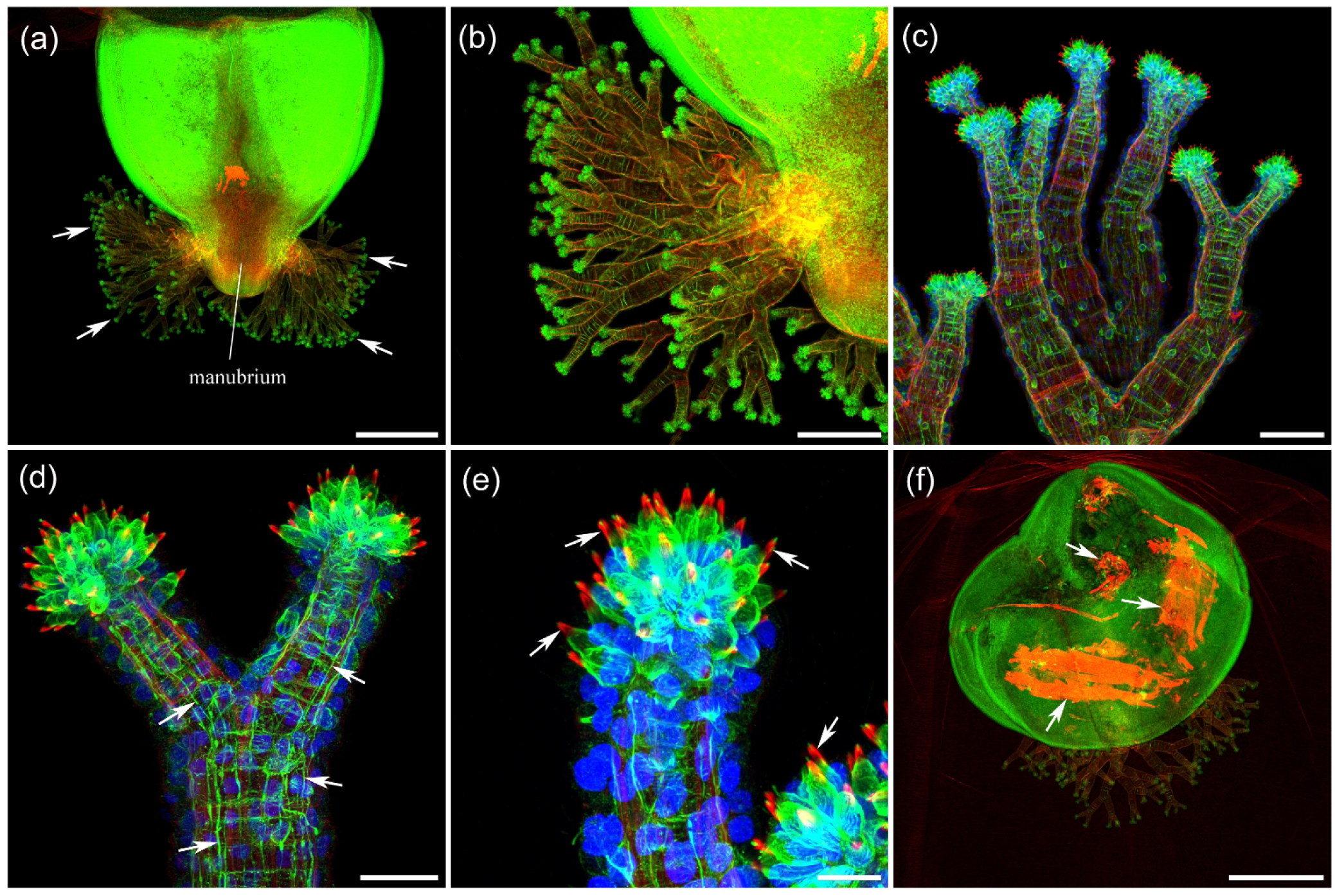
The structure of oral tentacles unique to *Bougainvillia*. **a, b** - A short manubrium bears several branching oral tentacles (arrows). **c** - Each tentacle bifurcates 3-4 times along its length. d - A dense neural network covers the walls of all tentacles and their branches and consists of many longitudinal neurites (arrows) labeled with tubulin AB, as well as shorter circular neural processes that connect them. **e** - Each tentacle branch at its end has a cluster of nematocysts, whose cell bodies are stained with tubulin AB. A single short sensory cilium at the end of each nematocyte is surrounded by a ring of stereocilia labeled with phalloidin in red (arrows). DAPI reveals a single large nucleus associated with each nematocyte. **f** - Remnants of half-digested crustaceans (most likely barnacle nauplii) are brightly labeled with phalloidin (arrows) and are frequently seen inside the stomach. Scale bars: **a, f** - 500 µm; **b** - 200 µm; **c** - 50 µm; **d** - 20 µm; **e** - 10 µm.

The numerous nematocytes at the tips of the oral tentacles and the dense innervation are presumably instrumental in feeding behavior, since Bougainvillia feeds on small crustaceans, especially barnacle nauplii, which must be captured and immobilized. The remnants of those crustaceans, brightly labeled with phalloidin, are frequently observed in the stomach (Fig. 10f).

#### *Stomotoca atra* (order Anthoathecata, family Pandeidae)

There are four radial canals and only two large tentacles, plus a number of rudimentary tentacles around the margin. The subumbrella does not reach the apical end of the bell. Halfway through the bell, it ends – the tissue folds and reverses direction back to the margin, forming a wide peduncle that holds a large manubrium extending well outside the bell.

The nerve ring lies at the margin of the bell and consists of many large, thin axons (Fig. 11a, b, f). From the bell margin, four bundles of smooth radial muscles run along each radial canal all the way to the manubrium. Each radial pathway along the radial canals also includes a radial neural tract originating at the nerve ring (Fig. 11a, b). This tract contains a pronounced, fully formed polygonal network consisting of thick threads, brightly labeled with anti-tubulin AB (Fig. 11b-d). Radial smooth muscle fibers are present only in the four radial pathways along the radial canals (Fig. 11d) and are absent from the rest of the subumbrella. Outside the four radial neural tracts, there are no neurons or neurites in the area occupied solely by the circular striated muscles. On very rare occasions, a single neuronal cell body without developed neurites can be seen near a radial canal, with one or two in the entire subumbrella region (Fig. 11e).

**Figure 11.**
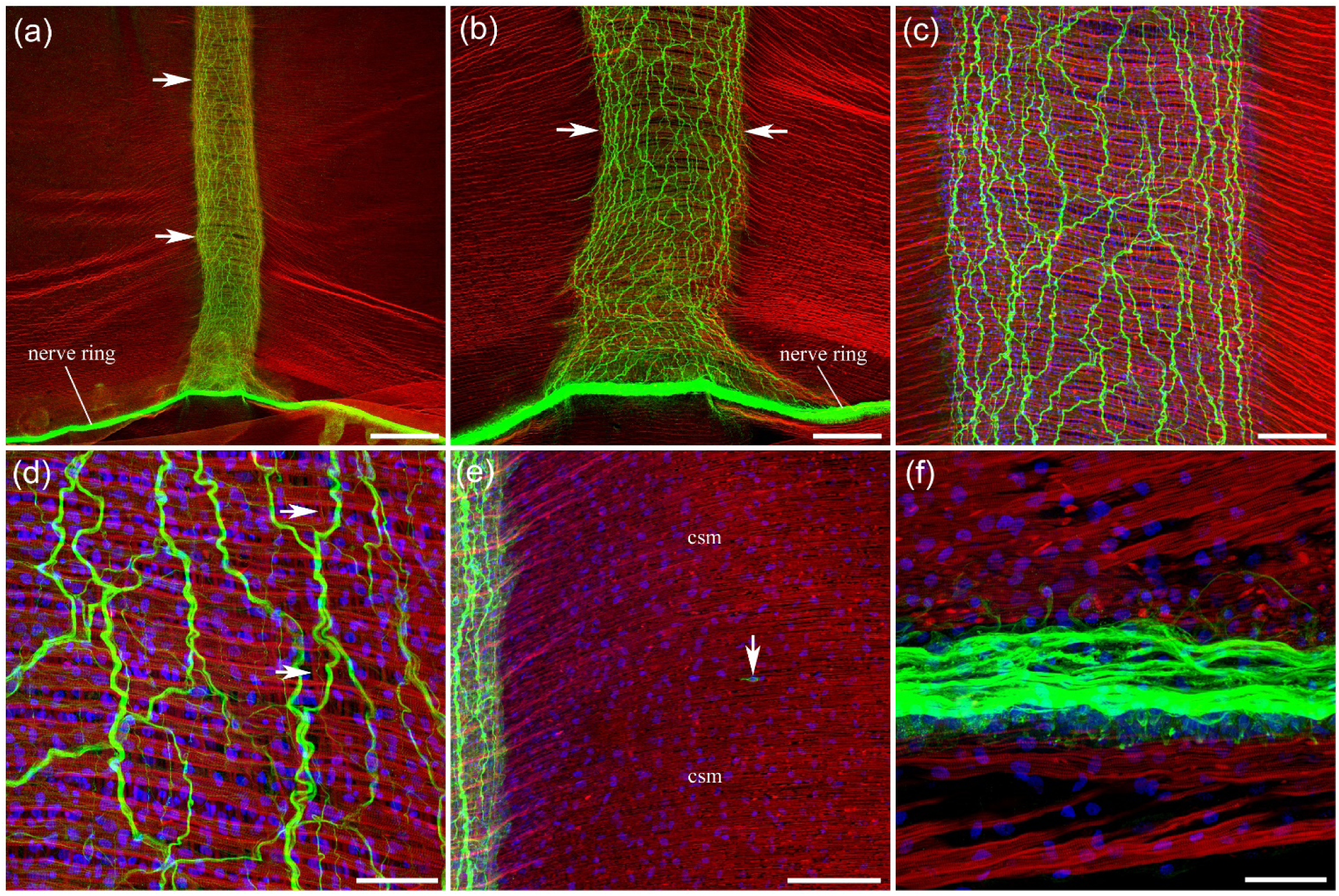
Neural and muscle systems in *Stomotoca*. **a, b** - A dense radial neural pathway (arrows) extends from the nerve ring at the margin of the umbrella toward the peduncle. **c, d** - The radial neural tract consists of a pronounced polygonal network of thick threads, brightly labeled with anti-tubulin AB. Smooth radial muscle fibers (arrows) are present within the radial pathway, though they are thinner and less dense than in other species. **e** - No neurons or neurites are found outside the radial neural pathways within the circular striated muscle (*csm*) layer. On very rare occasions, a single neuronal cell body (arrow) without developed neurites is observed not far from the radial canal. **f** - The nerve ring consists of numerous thick and thin neural processes. Scale bars: **a** - 500 µm; **b** - 200 µm; **c** - 100 µm; **d** - 40 µm; e - 100 µm; **f** - 30 µm.

The circular striated muscle layer is present in the subumbrella only halfway through the bell, up to the fold where a wide peduncle begins to form (Fig. 12a, e). That is where it is functionally needed to contract the bell for swimming. Notably, the polygonal neural network that forms the radial neural tract runs only from the nerve ring up to this fold, where the circular striated muscle layer ends (Fig. 12a). After the fold, and after the polygonal network ends, there are only numerous, mostly longitudinal thin neurites,and neural cell bodies remain in the pathway running along the smooth radial muscle bundles into the wide peduncle (Fig. 12a, b). This suggests that the polygonal neural network forming a neural tract in the subumbrella is not the neural pathway carrying a signal from the nerve ring to the manubrium but was rather important for the bundle of smooth radial muscles in the subumbrella, halfway through the bell before the peduncle, or maybe even for the circular striated muscle layer.

The walls of the radial canals themselves (in a different focal plane than the radial neural tract) contain a network of multipolar neurons (Fig. 12c). These neurons produce long neurites that run mostly longitudinally along the canals, but also establish side connections, forming a well-developed neural network.

**Figure 12.**
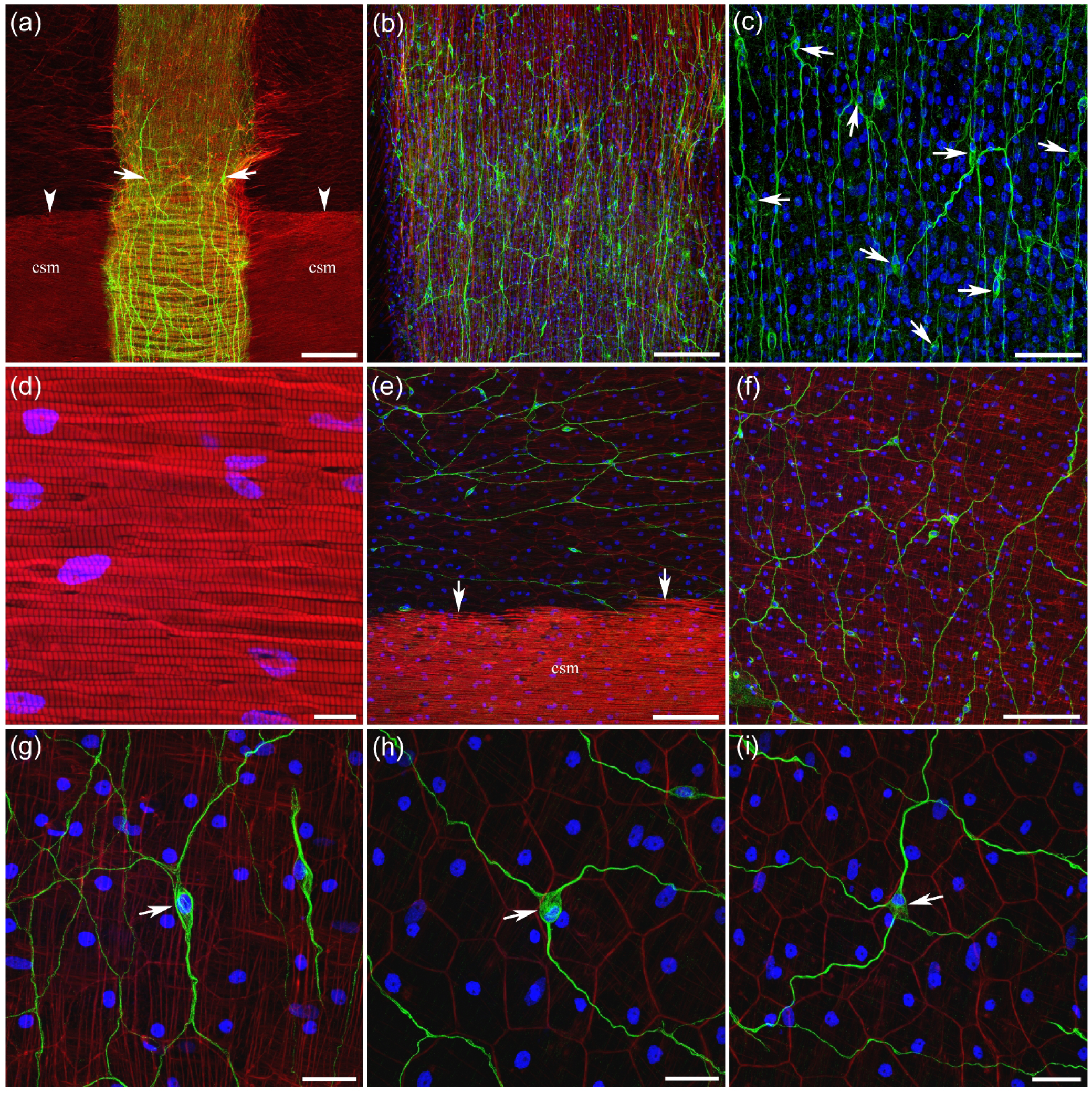
Neural network in the wide peduncle and radial canals of *Stomotoca*. **a** - Halfway through the bell, subumbrella ends; the tissue folds and reverses direction back to the margin, forming a wide peduncle. The circular striated muscle (*csm*) layer is present in the subumbrella only up to that fold (arrowheads). The thick polygonal neural network (arrows) in the radial neural pathway also extends only up to that fold. **b** - After the fold, when the polygonal network disappears, only numerous mostly longitudinal thin neurites and neural cell bodies remain in the pathway along the radial smooth muscle fibers traveling further toward the manubrium. **c** - The walls of the radial canal next to the radial neural pathway (in a different focal plane) contain a network of multipolar neurons (arrows) and long neurites labeled with tubulin AB. **d** - Circular striated muscle layer labeled with phalloidin. **e** - After the circular striated muscle (*csm*) layer ends (arrows) and the subumbrella transforms into a wide peduncle, numerous neurites and neural cell bodies appear and begin to form a loose network. Near the striated muscle layer and the fold, the neurites have a mostly circular orientation. **f** - Further from the fold and deeper into peduncle, their orientation is more radial. **g, h, i** - Neurons (arrows) that form this network in the peduncle are mostly bipolar (g) and tripolar (h) cells, while a few have four neurites (i). Scale bars: **a** - 200 µm; **b** - 100 µm; **c** - 40 µm; **d** – 10 µm; **e, f** - 100 µm; **g, h, i** - 20 µm.

In the peduncle itself, where the circular striated muscle layer and the subumbrella polygonal neural network do not extend, numerous neurons are distributed throughout the entire peduncle area (Fig. 12e, f). Their long branches visibly connect to each other, forming a network. Closer to the striated muscle layer and the peduncle fold, the neurites are mostly circular in orientation (Fig. 12e). Further from the fold, their orientation is more radial (Fig. 12f). The neurons forming the network in the peduncle are mostly bipolar and tripolar, although some cells produce four neurites (Fig. 12g-i).

#### *Neoturris breviconis* (order Anthoathecata, family Pandeidae)

There are four wide radial canals with the jagged outlines. Numerous small tentacles are attached to the margin of the bell.

The nerve ring at the margin is clearly defined (Fig. 13a). The entire subumbrella area is covered with a fully formed polygonal neural network (Fig. 13a, b; see also Mackie and Meech, 2008). Long neurites join to create thicker multi-neurite threads, forming a fully developed network of irregular polygonal units of varying sizes, elongated in the radial direction and perpendicular to the circular striated muscle fibers (Fig. 13b, c). Smooth radial muscle fibers in the subumbrella are evenly distributed across the entire subumbrella area (Fig. 13b, c). The neural network area matches the area of smooth radial muscle distribution. The polygonal neural network is formed by a number of neurons with long neurites. Most neurons are bipolar or tripolar, while some have four processes extending from the cell body (Fig. 13d-f).

**Figure 13.**
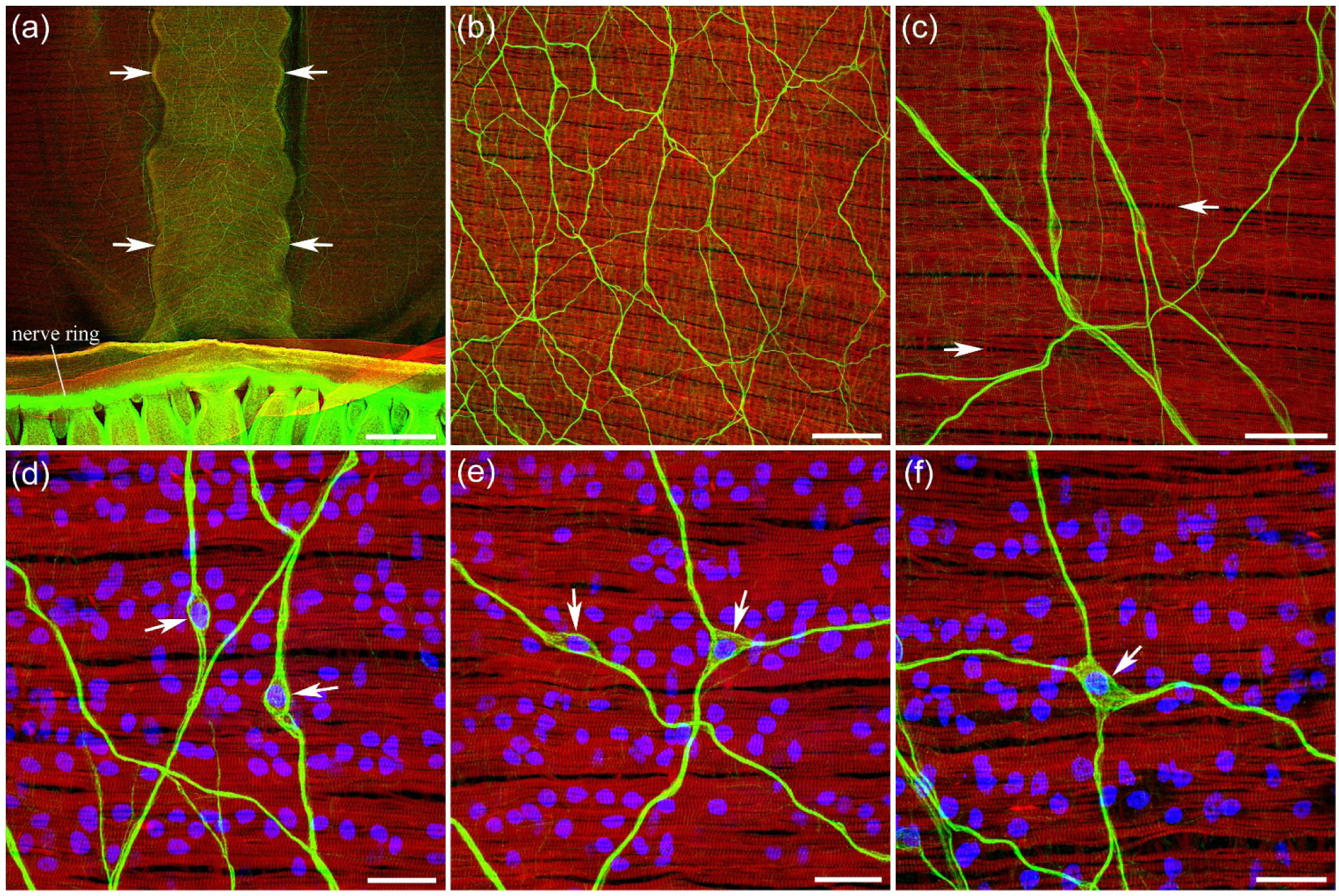
Neural network in the *Neoturris* subumbrella. **a** - A wide radial canal with jagged outline is indicated by arrows. **b** - The entire subumbrella region is covered with the neural network, which consists of irregular polygonal units of different sizes, elongated in the radial direction and perpendicular to the circular striated muscle fibers. **c** - Long, thin neurites merge to form thicker multi-neurite threads and create a fully developed network. Thin radial smooth muscle fibers (arrows) are evenly distributed across the entire subumbrella area. **d, e, f** - The neural network is formed by a number of neurons (arrows) with long neurites. Most neurons are bipolar or tripolar, while some have four processes extending from the cell body. Scale bars: **a** - 500 µm; **b** - 100 µm; **c** - 40 µm; **d, e, f** - 20 µm.

The polygonal neural network crosses the radial canal area without noticeable change in its structure (Fig. 14a). However, the network is slightly denser along the radial canals, as if it were more compressed there (Fig. 14b, c).

**Figure 14.**
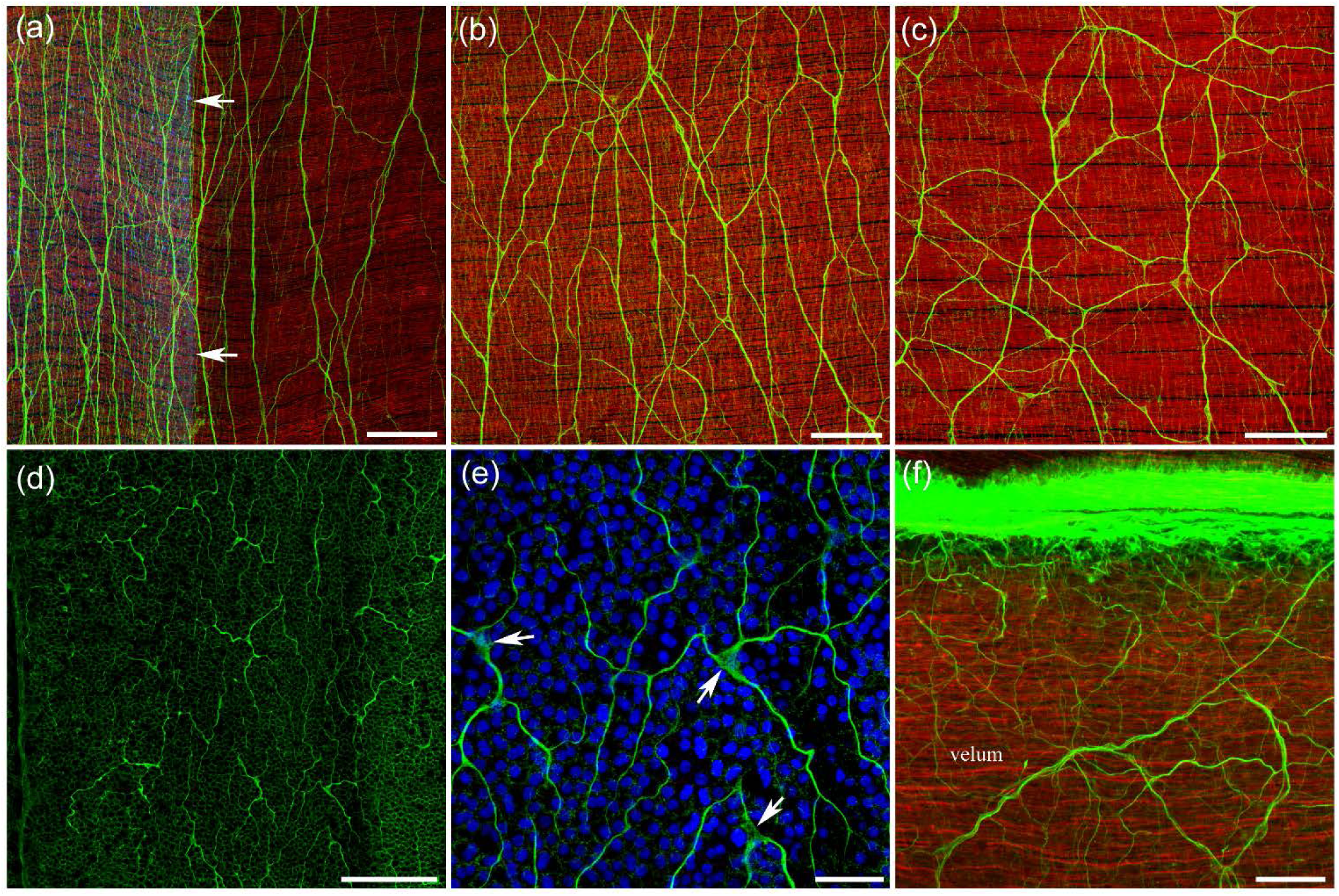
*Neoturris* neural networks in the subumbrella, radial canals, and velum. **a** - A polygonal neural network extends into the radial canal area (arrows) without noticeable change in its structure. **b** - However, the network is slightly denser along the radial canals, as if it were more compressed from the sides. **c** - Outside the radial canal area, the polygonal units are wider. **d, e** - The radial canal walls contain neurons with processes, many of which are oriented longitudinally along the radial canal. These are clearly located in a different focal plane than the subumbrella neural network. Some neuronal cell bodies are indicated by arrows. **f** - There is also a network of neurites attached to the nerve ring, which innervates the velum. Scale bars: **a, b, c, d** - 100 µm; **e, f** - 20 µm.

The radial canal walls contain numerous multipolar neurons with processes, many of which are oriented longitudinally along the radial canal pathway, forming a neural network (Fig. 14d, e). They are clearly in a different focal plane than the subumbrella polygonal neural network.

There is also a network of neurites and neural cell bodies attached to the nerve ring that innervates the velum (Fig. 14f).

#### *Leuckartiara longicalcar* (order Anthoathecata, family Pandeidae)

There are four radial canals. A few dozen long tentacles are attached to the margin. *Leuckartiara* is a close relative of *Neoturris* and looks similar, except for a gelatinous, cone-shaped projection on the top of the bell.

The nerve ring at the margin is brightly labeled with the tubulin AB (Fig. 15a). A neural network in subumbrella structurally resembles the network in Neoturris – a slightly elongated, radial, polygonal neural net (Fig. 15a-c). It consists of thick, multi-neurite threads that form irregular polygonal units and are produced by a number of mostly bipolar and tripolar neurons (Fig. 15c, d). The main difference is that the polygonal network does not cover the entire subumbrella area. It is restricted to a wide neural tract that runs along each radial canal, about three times the width of the canal itself (Fig. 15a, b). This radial pathway, which contains the neural network, also contains radial smooth muscle fibers, clearly present among the circular striated muscles (Fig. 16c, d). The neighboring areas lack neural elements and radial smooth muscle fibers - only circular striated muscles are present.

**Figure 15.**
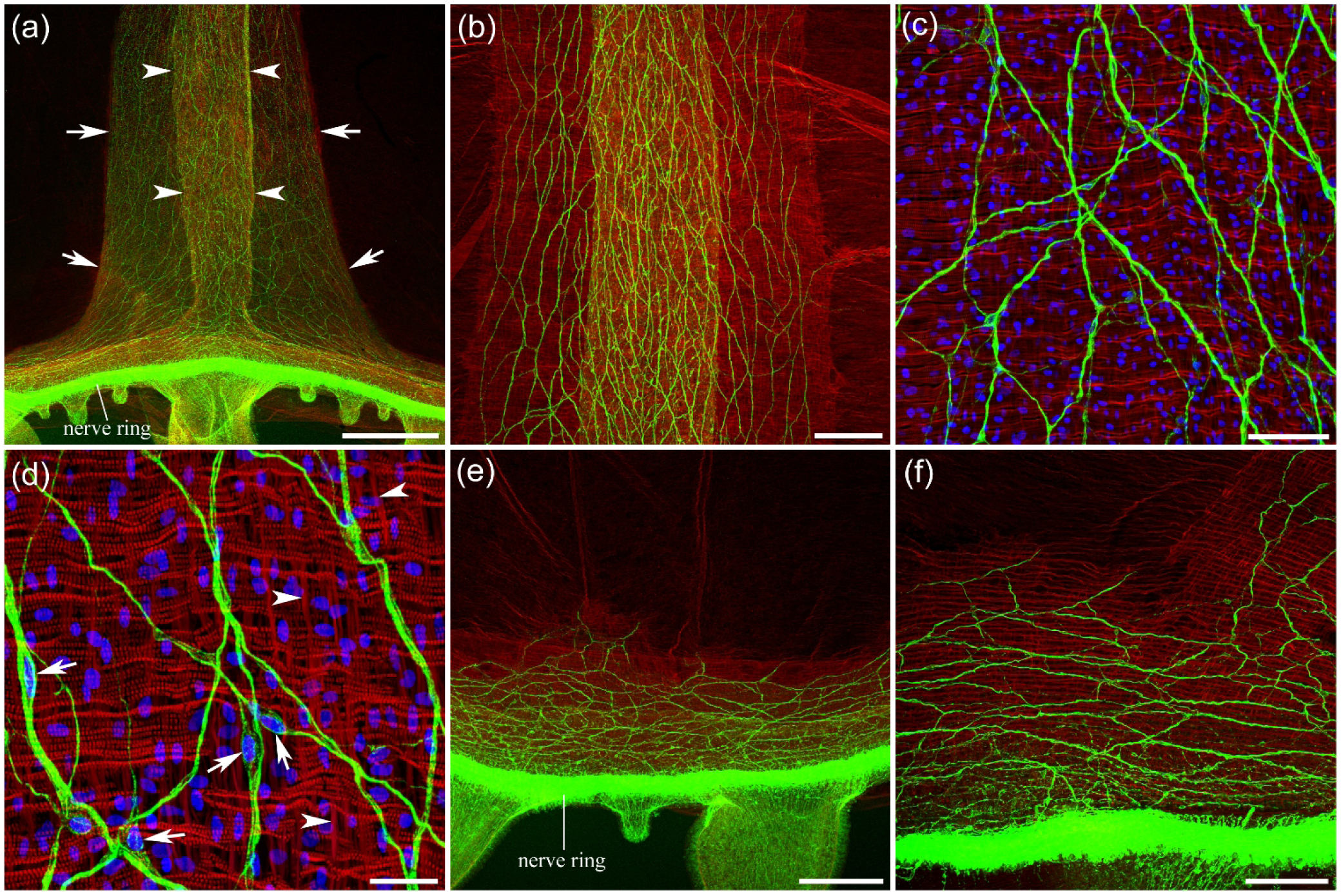
Neural and muscle systems in *Leuckartiara*. **a** - Along each radial canal (outlined by arrowheads), there is a wide radial pathway (arrows), which contains radial smooth muscle fibers and a neural tract. **b, c** - The radial neural tract consists of a network of irregular, elongated, polygonal units, very similar to the *Neoturris* neural network. **d** - The neural cell bodies forming the polygonal network are clearly visible at high magnification with DAPI staining (arrows). Arrowheads point to radial smooth muscle fibers. **e, f** - A wide band of the same polygonal network near the nerve ring connects neighboring radial neural tracts. Scale bars: **a** - 500 µm; **b** - 200 µm; **c** - 50 µm; **d** - 20 µm; **e** - 200 µm; **f** - 100 µm.

**Figure 16.**
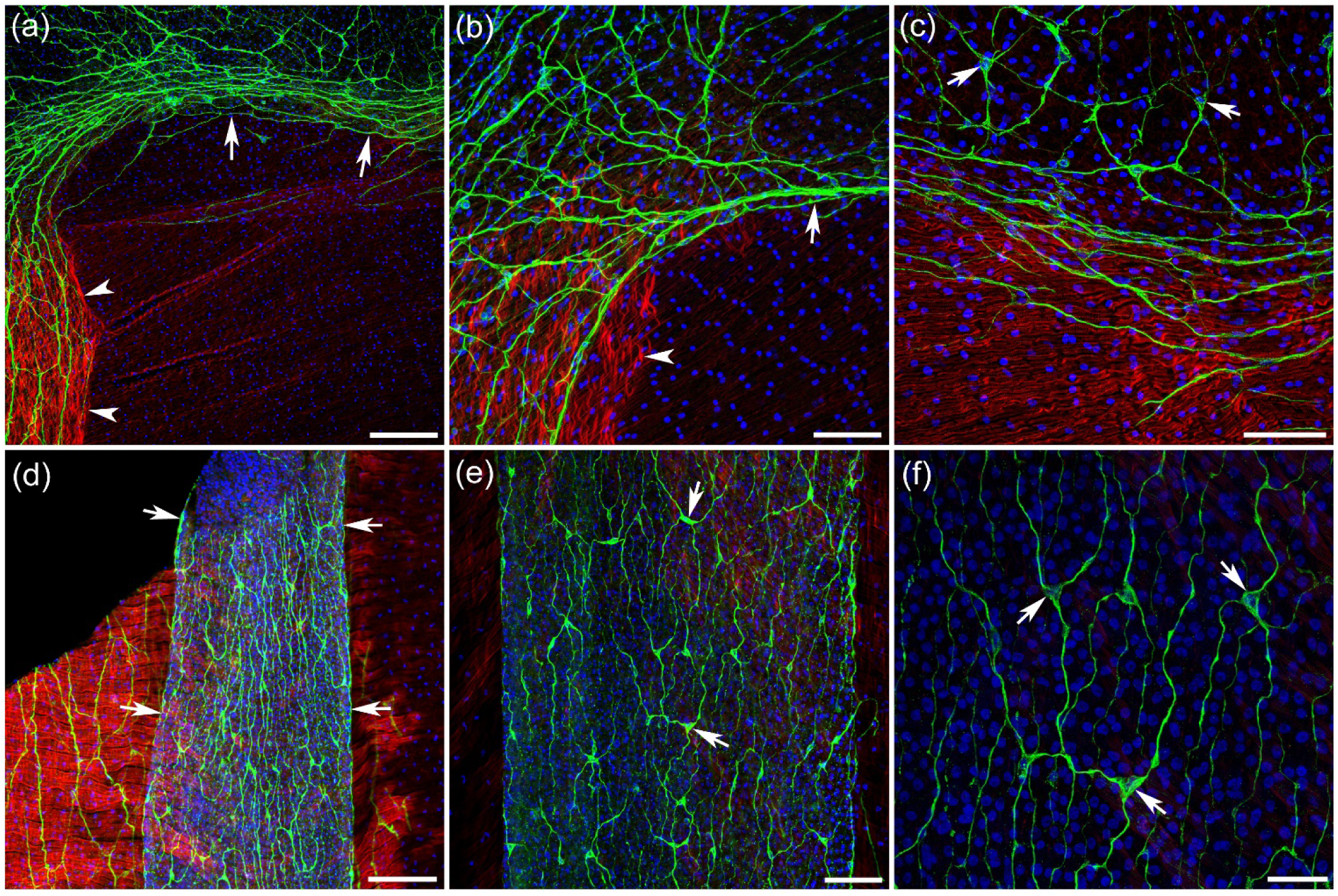
*Leuckartiara* neural network in the apical subumbrella and radial canals. a, b - At the apical end of the subumbrella, where the circular striated muscle layer ends at the base of the peduncle, the polygonal neural network of a radial neural tract turns and continues as a circular neural band (arrows) connecting the neighboring radial pathways. The wide radial muscle tract (arrowheads) also ends at the peduncle base, at the edge of the striated muscle layer. **c** - While connecting neighboring radial neural tracts, this neural band also forms connections with multipolar neurons (arrows) in the peduncle area. **d** - The diagonal cut through the subumbrella (in the upper left corner) provides a clearer view of a radial canal (arrows) and its position relative to the myoepithelial layer. **e, f** - The walls of the radial canals contain a network of neurons (arrows) with multiple processes labeled with tubulin AB. Scale bars: **a** - 100 µm; **b** - 50 µm; **c** - 40 µm; **d** - 100 µm; **e** - 50 µm; **f** - 20 µm.

In addition to the polygonal network in the radial pathways, there is a band of the same network along the ring nerve, connecting neighboring radial neural tracts (Fig. 15e, f). At the apical end of the subumbrella, a neural network lane also connects neighboring radial neural tracts and appears to be a continuation of the same polygonal network (Fig. 16a-c). Where the subumbrella reaches the top of the bell and connects to the peduncle, the circular striated muscle layer ends. The wide radial pathway, which contains radial smooth muscles, also ends at the fold, at the edge of the striated muscle layer. The neural network, however, turns and continues to the neighboring radial neural tract (Fig. 16a-c). It also connects to multipolar neurons with long neurites in the peduncle area (Fig. 16c).

The walls of the radial canals contain a network of neurons with multiple processes labeled with tubulin AB (Fig. 16d-f). They form a neural network very similar to the neural network in the walls of radial canals in *Neoturris*.

#### *Aequorea victoria* (order Leptothecata, family Aequoreidae)

The bell can be up to 10 cm in diameter. There are many radial canals running across the subumbrella, a distinctive feature of this species. Numerous thin tentacles are attached to the bell margin. The manubrium is short and elastic – the mouth can open wide, leading to a large stomach chamber. It consumes large prey such as other jellyfish and ctenophores. We saw it eat and digest a pteropod mollusc, *Clione limacina*.

The nerve ring labeled with tubulin AB encircles the entire umbrella at the margin (Fig. 17a). It consists of the inner and outer nerve rings, which are clearly separated (Fig. 17b; 18a). Many neurites exit the nerve ring to innervate each tentacle bulb (Fig. 17i). A narrow network of neurons and neurites also extends above the nerve ring in the subumbrella area (Fig. 17b, i; 2a).

**Figure 17.**
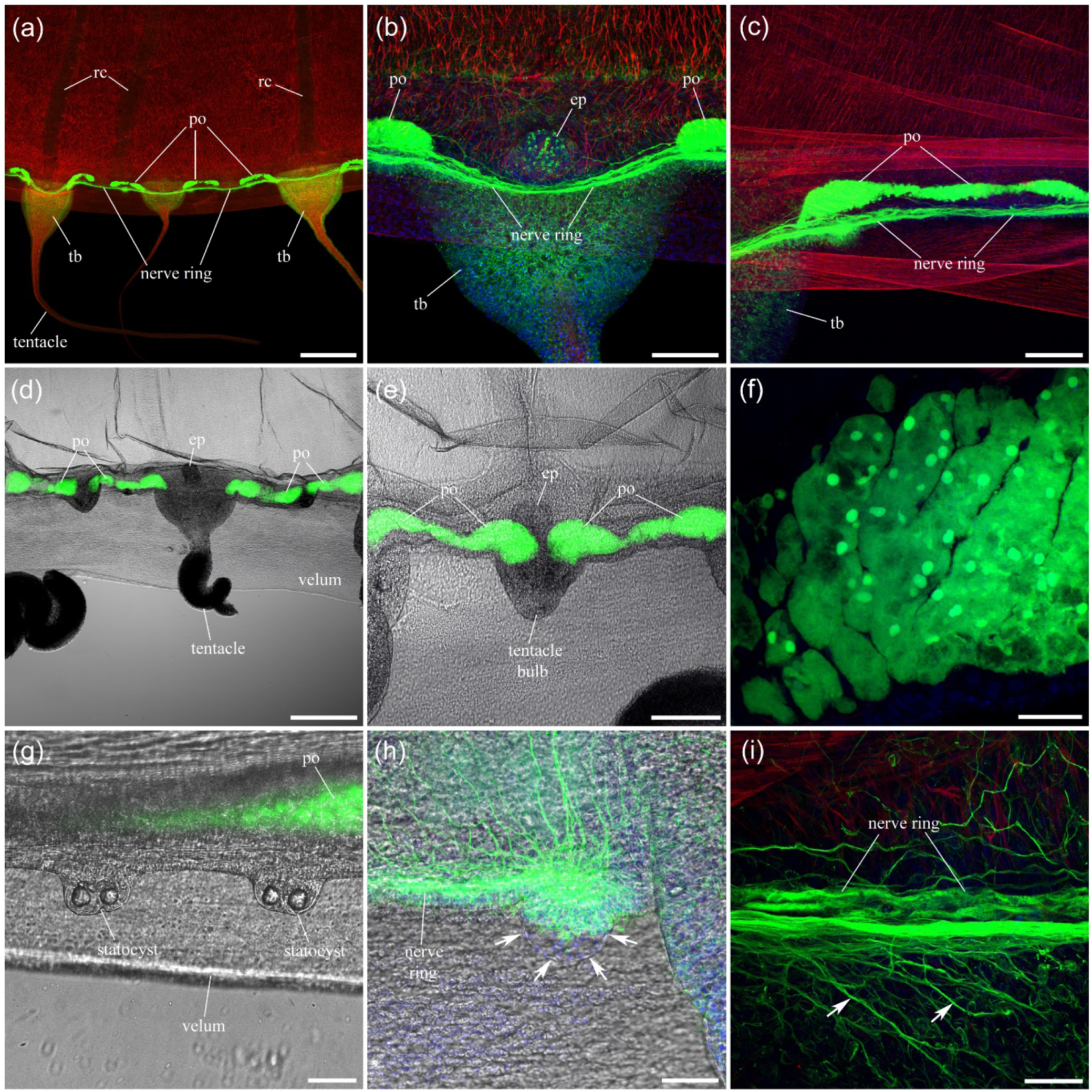
Neural and sensory elements labeled with tubulin AB at the margin of the bell in *Aequorea*. **a** - The relatively thin nerve ring encircles the margin of the bell, which bears many small tentacles. Each tentacle bulb (*tb*) has two photo-organs (*po*) next to the nerve ring, which contain GFP and produce bright green fluorescence. Note also many radial canals (*rc*), which are characteristic of this species. **b** - In addition to photo-organs (*po*), one on each side, each tentacle bulb (*tb*) has a single large excretory papilla (*ep*) near its center line on the adaxial side of the tentacle bulb, pointing into the subumbrella. **c** - Each photo-organ is elongated and runs parallel to the nerve ring. **d, e** - Margin of the bell in the confocal DIC light without any immuno- or phalloidin labeling in the preparation. Photo-organs (*po*) showing bright green fluorescence are located along the margin between the tentacle bases. Note also the position of the excretory papillae (*ep*) at the midline of each tentacle base, including both fully developed tentacles and newly growing tentacle bulbs. **f** - Photo-organs consist of a number of individual lobes, clearly visible at higher magnification. There are also many small granules or vacuoles inside the lobes that have much brighter fluorescence than the rest of the lobe tissue. **g** - There are usually two small statocysts (sometimes 3 or 4) between neighboring tentacles, each containing mostly two statoliths (sometimes 3 or 4). This is a live (not-fixed and not-labeled) preparation. **h** - Each statocyst (arrows) is attached to the bell margin very close to the nerve ring (green). There is always a noticeable swelling of the nerve ring, with significantly more neural processes at the point of attachment. Statoliths always disappear (dissolve?) after the immuno protocol, making it more difficult to see statocysts. **i** - Many neurites (arrows) exit the nerve ring and innervate the tentacle bulb. Scale bars: **a, d** - 500 µm; **b, c** - 100 µm; **e** - 200 µm; **f** - 20 µm; **g, h** - 50 µm; **i** - 25 µm.

*Aequorea victoria* was the original source of the Green Fluorescent Protein (GFP), which is widely used as a marker in many areas of genetics and molecular biology (Shimomura et al., 1962; Shimomura 1979, 2005; Morise et al., 1974). GFP is concentrated in specialized, elongated, gland-like photo-organs located at the margin of the bell between the tentacle bases (Fig. 17a, b). Harvey (1921) described, under the microscope examination, the “oval masses of yellow tissue corresponding in position with the luminous areas at night” - “spots along the edge of the umbrella at the base of the tentacles”. The photo-organs exhibit bright-green fluorescence in live or fixed animals without any labeling – mostly in the green channel (slightly in blue), with no signal in the red spectrum (Fig. 17d, e). The photo-organs are thin and relatively long, running along the nerve ring and positioned on both sides of each tentacle bulb (Fig. 17c-e). The photo-organs lie in the focal plane of the exumbrella rather than on the subumbrella side, and there is an obvious gap between the nerve ring and the photo-organs, with no noticeable neurites connecting them. As seen at higher magnification, the photo-organs consist of distinct lobes that also contain numerous granules or vacuoles with much brighter fluorescence than the lobes themselves (Fig. 17f).

A number of small statocysts (about 80 µm in diameter) are attached to the bell margin, usually two but sometimes three or four between neighboring tentacles or tentacle bulbs (Fig. 17g). Each statocyst contains mostly two, sometimes three or four, statoliths (Fig. 17g). Statocysts are attached to the bell margin very close to the nerve ring, which thickens noticeably at the point of attachment and contains more short neurites running from it (Fig. 17h).

Radial neural tracts are present along the radial canals in *Aequorea*, as in other hydrozoans (Fig. 18a). These tracts consist of several mostly longitudinally oriented neurites from bipolar or tripolar neurons, extending along the entire length of the tracts (Fig. 18b). The neural tracts are not dense but are visually distinct from the rest of the subumbrella network. Numerous neurons and neurites are also found throughout the subumbrella between the radial neural tracts (Fig. 18; see also Satterlie, 2008). In small adults, 2-3 cm in diameter, these neurons do not form a fully developed neural network but rather represent a collection of neurons with processes spread across the subumbrella (Fig. 18c). Most neurons are bipolar cells (Fig. 18c, d), although neurons with three or four branches are not uncommon. Throughout the subumbrella, these neurons and their neurites are mostly radially oriented (Fig. 18c, d). In large *Aequorea*, 7-8 cm in diameter, the subumbrella neurons have longer, more extensively branching neurites that establish multiple connections with one another and form a fully developed neural network (Fig. 18e, f; see also Satterlie, 2008). The neural network covers the entire subumbrella region and is slightly elongated in the radial direction.

**Figure 18.**
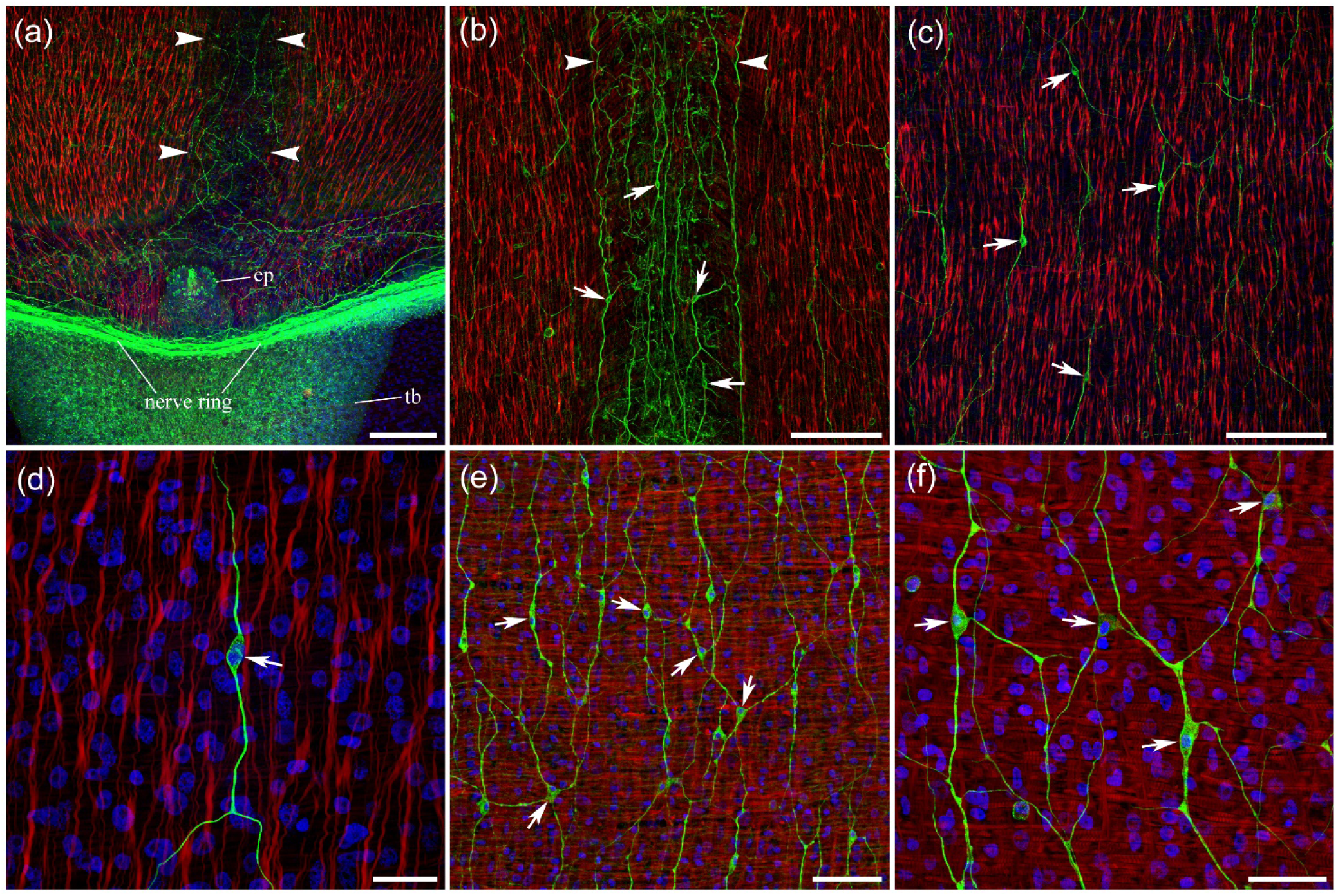
The structure of the *Aequorea* neural and muscle systems in the subumbrella. **a** - The radial neural tract (outlined by arrowheads) runs along the radial canal and originates at the nerve ring, predominantly at the line crossing the center of the tentacle bulb (*tb*) and the excretory papilla (*ep*). **b** - Each radial neural tract (arrowheads) consists of several mostly longitudinally oriented neurites produced by bipolar and tripolar neurons (arrows), spread along its entire length. **c, d** - In small adults (2-3 cm), numerous individual neurons (arrows) are distributed throughout the subumbrella without forming a neural network. Most of these subumbrella neurons are bipolar cells (arrows), with neurites oriented radially, parallel to the smooth radial muscle fibers (labeled with phalloidin in red). **e, f** - In large adults (7-8 cm), the subumbrella neurons (arrows) form a fully developed neural network. Note the circular striated muscles and smooth radial muscle fibers running perpendicular to them. Scale bars: **a, b, c** - 100 µm; **d** - 20 µm; **e** - 50 µm; **f** - 30 µm.

*Aequorea victoria* has conspicuous excretory papillae on the adaxial side of the tentacle bulbs, pointing into the subumbrella. Each tentacle bulb has a single long excretory papilla at its midline (Fig. 17d, e). Excretory papillae are clearly visible in stained preparations because tubulin AB brightly labels a number of cone-shaped cells on their surface that resemble nematocysts (Fig. 17b; 18a). Each excretory papilla also has a dense neural network in its walls (Fig. 19a). The gonads, which are attached to the radial canals and run along the radial pathways, are long and massive in large *Aequorea* and extend almost to the margin of the bell. Numerous individual neurons with neurites are labeled with tubulin AB within the gonads (Fig. 19b, c). Attached to the subumbrella at its apical end, a wide peduncle has a fully formed neural network covering its entire surface in both small and large adult *Aequorea* (Fig. 19d-f). However, in large animals, the network is much denser and contains significantly more neurons (Fig. 19e, f).

**Figure 19.**
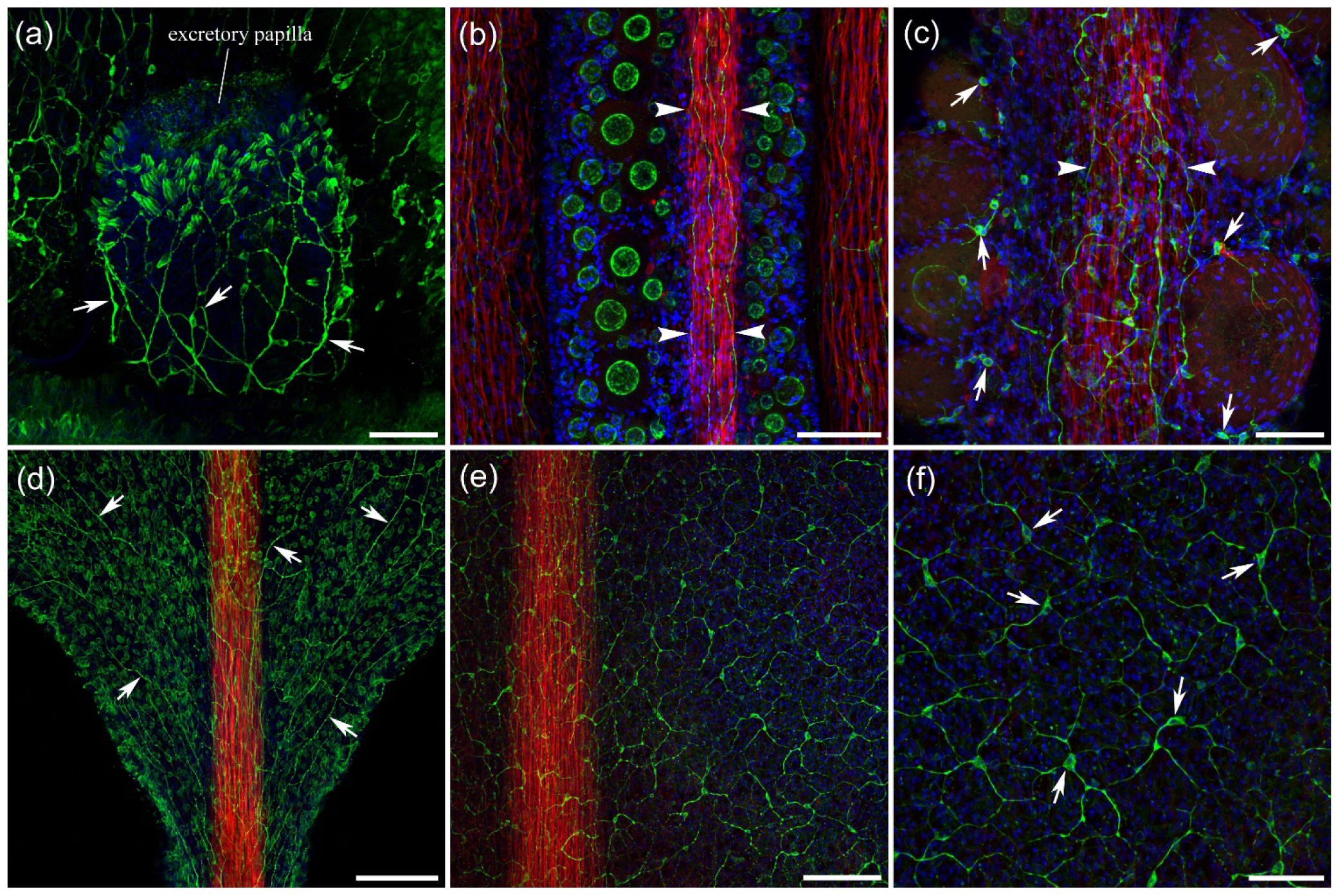
Innervation of the peduncle, gonads, and excretory papillae in *Aequorea*. **a** - Each excretory papilla (*ep*) is densely innervated by branching neurites (arrows) that form a network in its body wall. Note also numerous cone-shaped cells outlined by tubulin IR, which are located mostly at the distal end – these are nematocysts. **b, c** - Gonads are attached to the radial canal and run along the radial neural and muscle pathway (outlined by arrowheads). Arrows point to neurons inside gonads. **d** - The peduncle area has a fully formed neural network (arrows) covering its entire surface, even in small, 2-3 cm animals. The radial smooth muscle fibers in the peduncle area form a dense radial muscle tract that runs through peduncle to manubrium (labeled with phalloidin in red). **e, f** - In large adults, there are significantly more neurons (arrows) that form a much denser neural network in the peduncle. Scale bars: **a, c, f** - 50 µm; **b, d, e** - 100 µm.

Radial smooth muscle fibers are distributed throughout the subumbrella region (Fig. 18). They are consistently visible alongside the subumbrella neurons and neurites (Fig. 18d). Near the bell margin and the nerve ring, there are no distinct radial muscle tracts – instead, smooth radial muscle fibers are evenly distributed across the entire subumbrella area. However, away from the margin, the radial smooth muscle fibers become much denser along the radial canals, forming clear, very dense radial smooth muscle tracts that run through the subumbrella and then the entire peduncle to manubrium (Fig. 19b-e).

#### *Mitrocoma cellularia* (also known as ***Halistaura cellularia***; order Leptothecata, family Mitrocomidae)

The umbrella can reach up to 10 cm in diameter – this is a large jellyfish, comparable in size to *Aequorea*. There are four radial canals, each with a long, firmly attached gonad along most of its length. The wide, prominent peduncle has a very short manubrium with four long, ruffled lips. The bell margin bears numerous thin tentacles. It exhibits strong bioluminescence localized to a narrow band around the margin.

*Mitrocoma* (*Halistraura*) was among the first hydrozoan species, along with *Aequorea*, used to study bioluminescence (Harvey, 1921; Shimomura et al., 1963). As Harvey (1921) noted, unlike in *Aequorea*, “in *Mitrocoma*, the yellow masses” responsible for bioluminescence and located at the edge of the umbrella “are much closer together forming an almost continuous line”. Indeed, the photo-organs in *Mitrocoma* appear as a continuous, narrow band running uninterrupted next to the nerve ring and encircling the bell margin (Fig. 20a, b). Short, thin branches presumably connect the photo-organ to the nerve ring, as revealed by thin optical sections through the middle of the margin and photo-organ (Fig. 20c). Most of the photo-organ fluorescence is in the green spectrum, with significantly weaker fluorescence in the blue (Fig. 20d). In preparations without immuno- or phalloidin labeling (without fluorescent tracers), photo-organs are clearly visible, showing natural bright green fluorescence along the bell margin (Fig. 20e, f_i_). Separating the green and blue channels shows weaker photo-organ fluorescence in the blue spectrum, in addition to the strong green fluorescence (Fig. 20 f_ii_ and f_iii_). There is no signal in the red spectrum.

**Figure 20.**
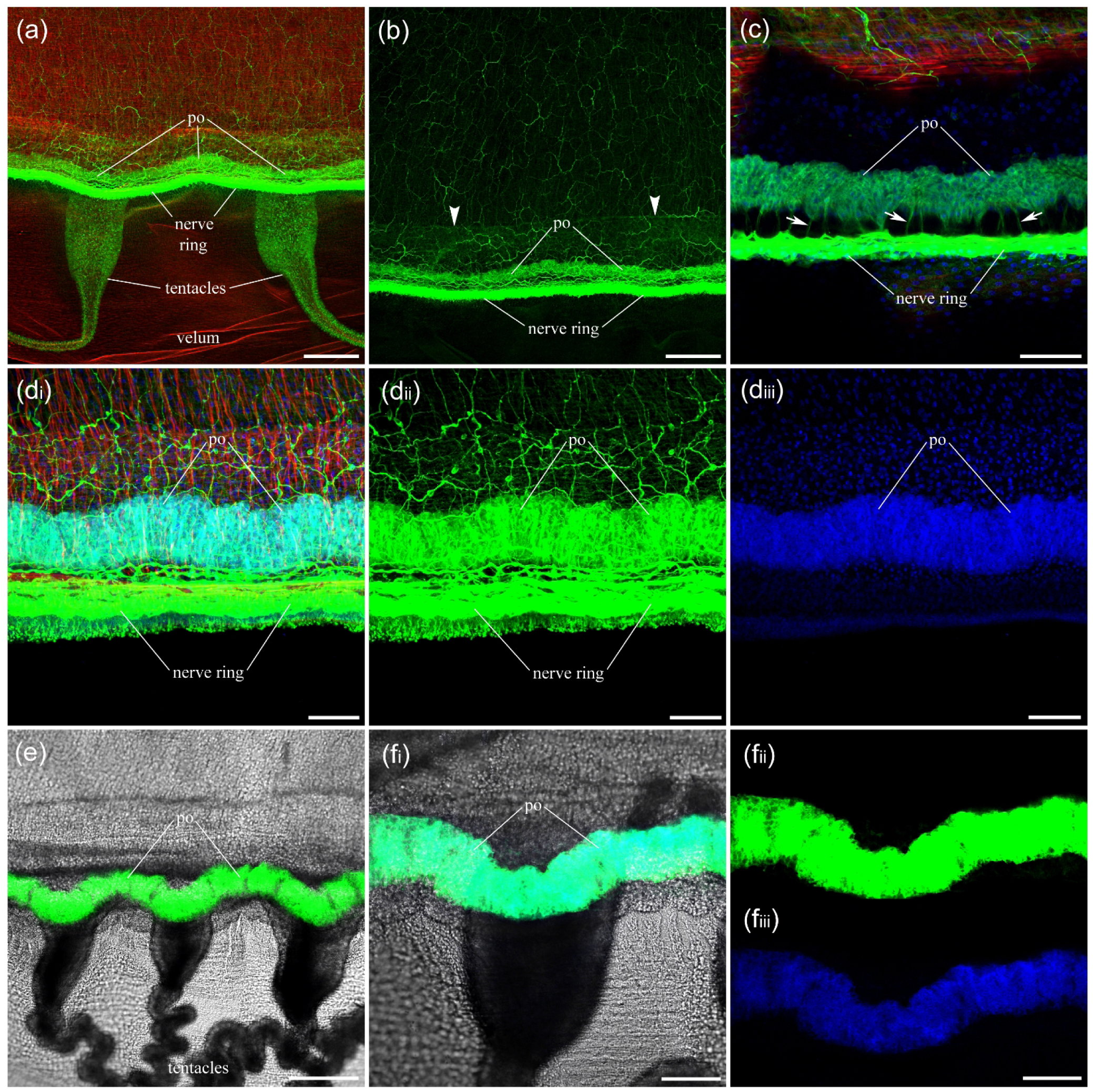
The nerve ring and photo-organs at the margin of the bell in *Mitrocoma*. **a** - The nerve ring, clearly labeled with tubulin AB in green, encircles the margin of the bell. The photo-organ (*po*), which contains a fluorescent peptide and produces bright green fluorescence, appears as a narrow, continuous band adjacent to the nerve ring and runs uninterrupted around the bell margin. **b** - Similar to (a), but in the green channel only, which makes the subumbrella neural network more visible above the nerve ring and photo-organ (*po*). Arrowheads indicate the circular canal at the margin of the bell. **c** - An optical section through the middle of the photo-organ reveals numerous short, thin branches that connect the photo-organ to the nerve ring (arrows). **d_i_** - The photo-organ (*po*) next to the nerve ring at higher magnification. Note also the smooth radial muscles labeled with phalloidin (red) in the subumbrella above the nerve ring. **d_ii_** - The green channel of the same preparation shows that most of the photo-organ fluorescence is in the green spectrum. **d_iii_** - The blue channel of the same preparation. There is some fluorescence from the photo-organ in the blue spectrum, although not as bright. **e, f_i_** - Margin of the bell with the confocal transmitted light detector in preparations without any immuno- or phalloidin labeling. Photo-organs (*po*) showing natural bright green fluorescence are located along the bell margin. **f_ii_** and **f_iii_** - The green and blue channels of the same preparation are shown separately and reveal that there is weaker photo-organ fluorescence in the blue spectrum in addition to the strong green fluorescence. Scale bars: **a, b** - 200 µm; **c, d** - 50 µm; **e** - 200 µm; **f** - 100 µm.

The polygonal neural network covers the entire subumbrella area above the nerve ring (Fig. 20b; 21a). The neural network is denser near the nerve ring and has more circular or neutral orientation near the bell margin (Fig. 21b). Away from the margin, across the rest of the subumbrella region, the polygonal network is stretched in the radial direction (Fig. 21c, d). Radial smooth muscle fibers are also distributed throughout the entire subumbrella area and do not form radial bundles along the radial canals (Fig. 21c, d). The distinctive wide peduncle is entirely covered with a fully formed polygonal neural network, which is also stretched in the radial direction (Fig. 21e). Radial smooth muscle fibers cover the entire peduncle area, as the neural network does. The radial canals in *Mitrocoma* contain numerous tiny non-neuronal cells on their surface, brightly labeled with tubulin IR, which completely mask any possible radial neural pathways in the subumbrella. However, focusing the optical sectioning away from the surface and deeper into the canal walls reveals that each radial canal contains several longitudinally oriented neural processes (arrows) produced by small bipolar neurons (Fig. 21f).

**Figure 21.**
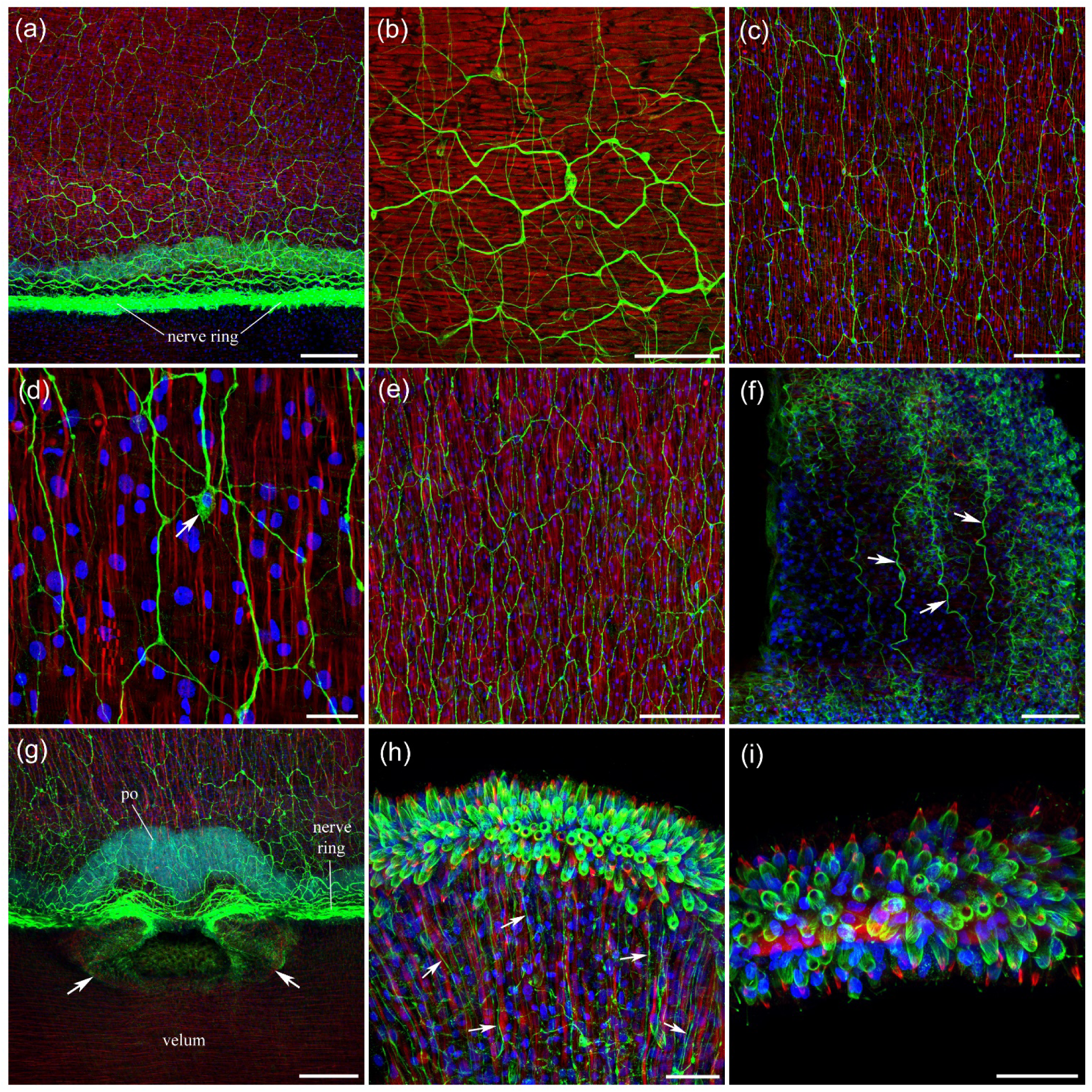
Neural networks and muscles in the subumbrella of *Mitrocoma*. **a** - A polygonal neural network covers the entire subumbrella region. **b** - The neural network is denser and has more circular orientation near the nerve ring and bell margin. **c** - Away from the margin and across the entire subumbrella, the polygonal neural network is stretched radially. Note the radial smooth muscle fibers (labeled with phalloidin in red), which are distributed throughout the subumbrella. **d** - The neural network is formed by multipolar neurons. The cell body of one such neuron is indicated by an arrow in this higher-magnification image, which also reveals radial smooth muscle fibers and some circular striated swim muscles. **e** - The short, wide peduncle has a fully formed polygonal neural network covering its entire surface and stretched radially. The radial smooth muscle fibers (labeled with phalloidin in red) cover the entire peduncle. **f** - The walls of each radial canal contain several longitudinally oriented neural processes (arrows) produced by small bipolar neurons. **g** - One of several vesicles (arrows), which are evenly apread and attached to the bell margin. Note the connection to the nerve ring. **h** - The long lips at the end of the manubrium contain, at their very edge, a tightly packed band of nematocysts. Tubulin IR reveals a number of neural processes innervating the nematocyst area. They run parallel to smooth muscle fibers labeled with phalloidin in red. **i** - Nematocysts also cover the entire surface of the numerous, long tentacles at the bell margin. The general structure of nematocysts and their labeling appear to be highly conserved and similar across all studied species. Scale bars: **a, c, e, g** - 100 µm; **b, f** - 50 µm; **d, h, i** - 20 µm.

The four long lips surrounding the mouth contain tightly packed groups of nematocysts along their edges (Fig. 21h). Tubulin IR reveals many neural processes approaching and presumably innervating those nematocyst groups (Fig. 21h). These processes run in parallel with a number of smooth muscle fibers in the lips. Nematocysts also cover the surface of the numerous tentacles at the margin of the bell (Fig. 21i). Several marginal vesicles are evenly distributed around the umbrella and appear to be connected to the nerve ring (Fig. 21g).

#### *Clytia gregaria (*formerly known as *Phialidium gregarium*; order Leptothecata, family Campanulariidae)

There are four radial canals, each with a thin, firmly attached gonad along its longitudinal side. The manubrium is very small and located at the top of the subumbrella. Numerous very short tentacles are attached to the bell margin.

The relatively thin nerve ring encircles the margin of the bell (Fig. 22a). A clear separation is visible between the inner and outer nerve rings (Fig. 22f). Small statocysts are attached to the margin of the bell next to the nerve ring between the tentacles (Fig. 23f). Radial neural tracts are present and run along the radial canals. However, they do not form tight nerves or dense, fully formed networks, butrather a collection of neurons and neurites running mostly in radial orientation (Fig. 22a-c). A narrow stretch of a network of neurons and neurites is also attached to the ring nerve in the margin area, connecting neighboring radial neural tracts (Fig. 22d-f).

**Figure 22.**
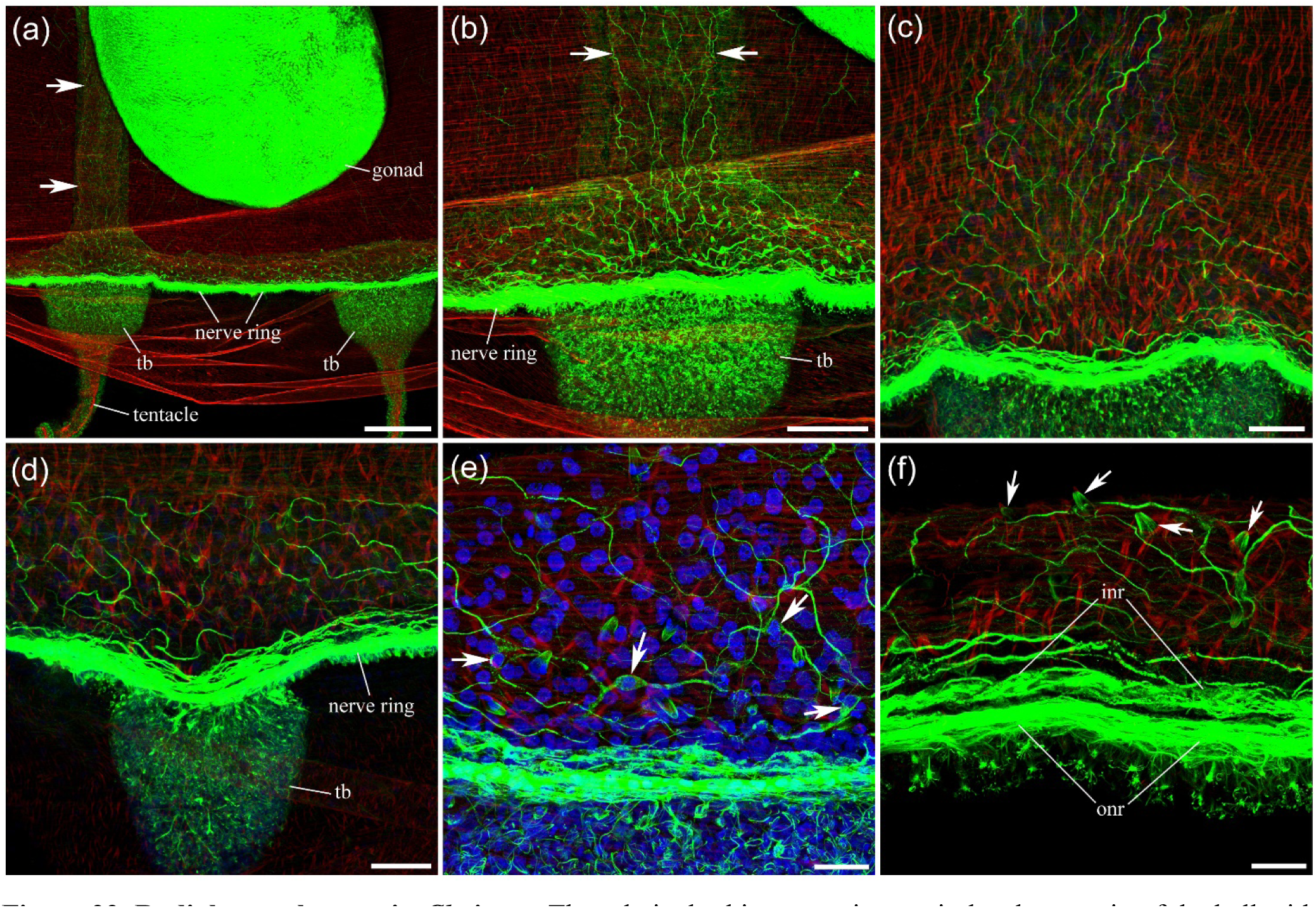
Radial neural tracts in *Clytia*. **a** - The relatively thin nerve ring encircles the margin of the bell with numerous short tentacles (*tb* – tentacle base). Radial neural tracts (arrows) are clearly present, originating from the nerve ring next to the tentacle base and running along the four radial canals. **b, c** - The radial neural tract (arrows) does not consist of fully formed nerves or networks but rather represents a collection of neurons and neurites oriented mostly radially. **d, e** - There are no radial neural tracts next to the bases of other tentacles that are not near the radial canals; only a narrow stretch of a network of neurons (arrows) and neurites, attached to the nerve ring and connecting neighboring radial neural tracts, is present. **f** - A narrow band of a network next to the nerve ring between the tentacles. Note the clear separation between the inner nerve ring (*inr*) and the outer nerve ring (*onr*). Arrows point to nematocysts. Scale bars: **a** - 200 µm; **b** - 100 µm; **c, d** - 50 µm; **e, f** - 20 µm.

**Figure 23.**
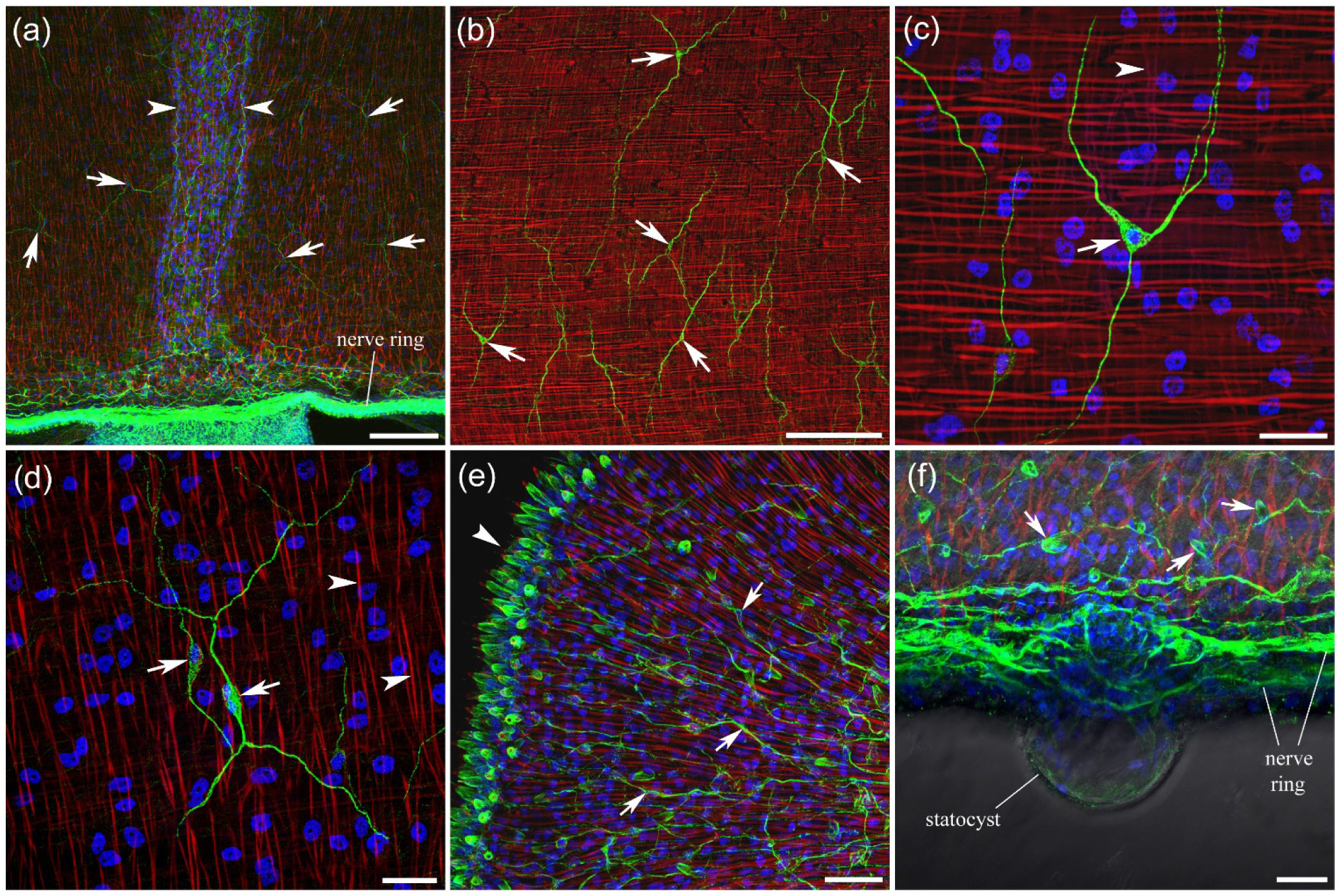
Neural elements in *Clytia* subumbrella. **a,b** - In addition to the radial neural tracts (arrowheads), numerous neurons (arrows) are present throughout the subumbrella region. They form a mesh of neurons with multiple long processes, distributed throughout subumbrella, some overlapping and others more isolated. **c, d** - Most neurons are bipolar or tripolar cells (arrows), although neurons with four branches are not uncommon. Note that radial smooth muscle fibers (arrowheads) are located throughout the entire subumbrella region. **e** - The lips of the manubrium have many neurites (arrows) forming a dense neural network. Note the numerous nematocysts at the edge of the lips (arrowhead). **f** - The statocyst attached to the nerve ring is clearly visible in DIC light. Arrows point to nematocysts at the bell margin. Scale bars: **a, b** - 100 µm; **c, d, e, f** - 20 µm.

In addition to the radial neural tracts and a narrow band of neural network along the nerve ring, numerous neurons and neurites are distributed throughout the subumbrella region (Fig. 23a, b). These neural elements form a loose mesh, with multiple long processes extending across the subumbrella area. Some neighboring neurons have overlapping processes, suggesting possible contact, whereas others are more isolated (Fig. 23b). Most neurons are bipolar or tripolar cells (Fig. 23c-d), although neurons with four branches are not uncommon.

The neural network is also present in the manubrium and its lips, where neurons and their long neurites form a dense meshwork that covers the entire area (Fig. 23e). The edges of the lips are densely populated with nematocysts (Fig. 23e).

There are no tight bundles of radial smooth muscle fibers in *Clytia*. Instead, individual radial smooth muscle fibers are distributed throughout the subumbrella region (Fig. 23a-d). Thus, the distribution of neural elements in the subumbrella again correlates with the radial smooth muscle distribution.

Natural bright-green fluorescence is observed in *Clytia* without labeling, predominantly in their numerous small tentacles (Fig. 24). The photo-organs are small, round or slightly elongated structures packed along the entire length of the tentacles and are clearly visible in both live (Fig. 24a-b) and fixed animals (Fig. 24d-f).

**Figure 24.**
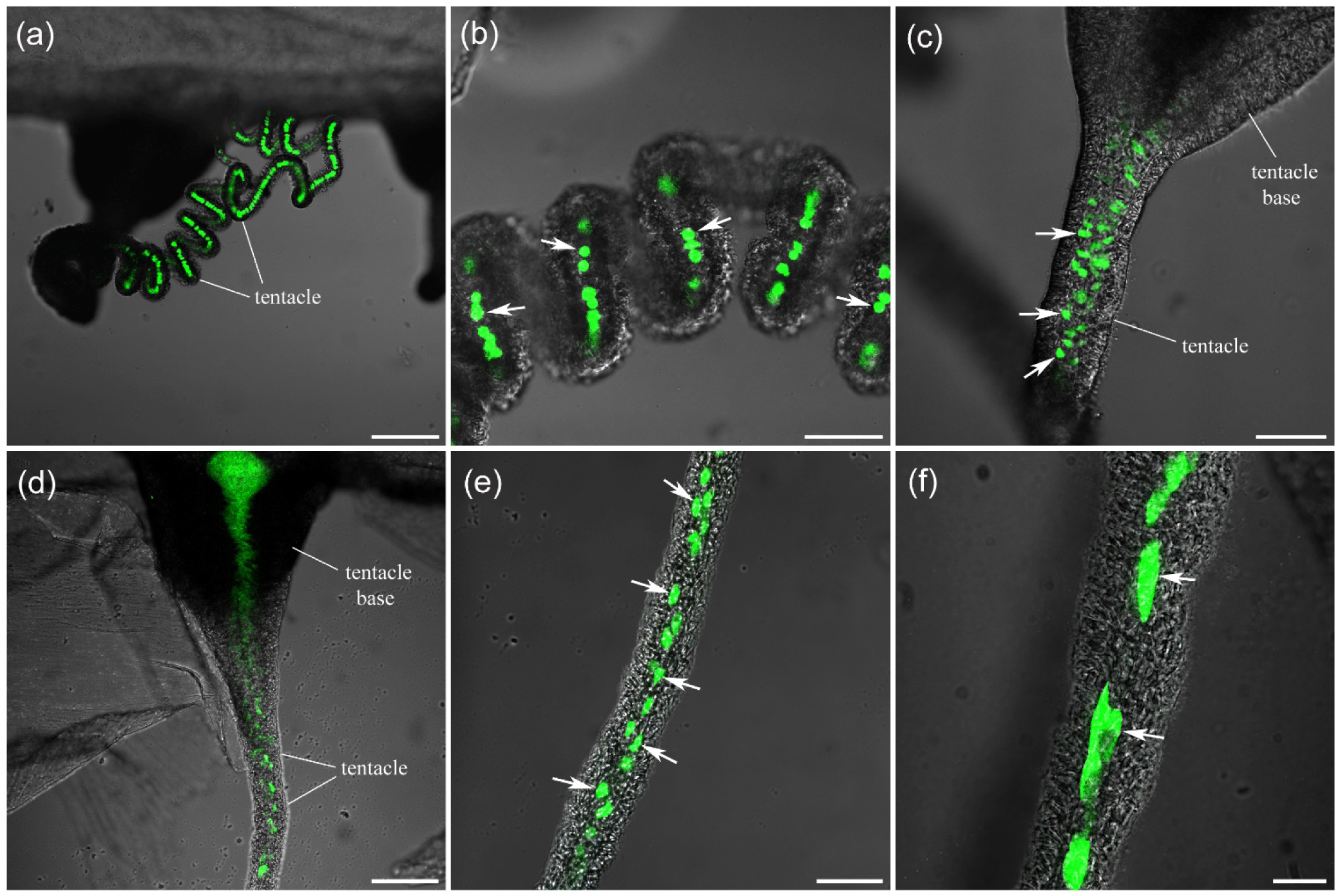
Natural green fluorescence in *Clytia* tentacles. All preparations are unlabeled and scanned with the confocal transmitted light detector. **a, b, c** – live preparations; **d, e, f** – preparations fixed in 4% paraformaldehyde and washed in PBS. Arrows point to some of the numerous round or slightly elongated structures along the tentacles that show bright green fluorescence. Scale bars: **a** – 200 µm; **b, c, e** – 50 µm; **d** – 100 µm; **f** – 20 µm.

#### *Eutonina indicans* (order Leptothecata, family Eirenidae)

A short manubrium with four lips is attached to the long, transparent peduncle and hangs below the bell margin. Four radial canals each have a long, sinuous gonad attached to them. Many short, thin tentacles are located along the bell margin.

The nerve ring, brightly labeled with the tubulin AB, encircles the margin of the bell (Fig. 25a, b). Four radial neural tracts originate at the nerve ring and extend across the subumbrella region along the radial canals to the manubrium (Fig. 25a, b). Each radial neural tract consists of several longitudinally oriented neurites produced by mostly bipolar and tripolar neurons located along the entire length of the tract (Fig. 25b, c). A distinct neural network also exists in the subumbrella between the radial neural tracts (Fig. 25b). In young, small adults (about 1-1.5 cm in diameter), subumbrella neurons are more individually identifiable, although neurites from neighboring neurons do overlap and apparently form connections (Fig. 25d). In large animals (up to 4 cm in diameter), a fully formed neural network consists of irregular polygonal units elongated in the radial direction (Fig. 25e). The bipolar, tripolar, and multipolar neurons that form the threads of this neural network are more clearly identified with DAPI labeling (Fig. 25f).

**Figure 25.**
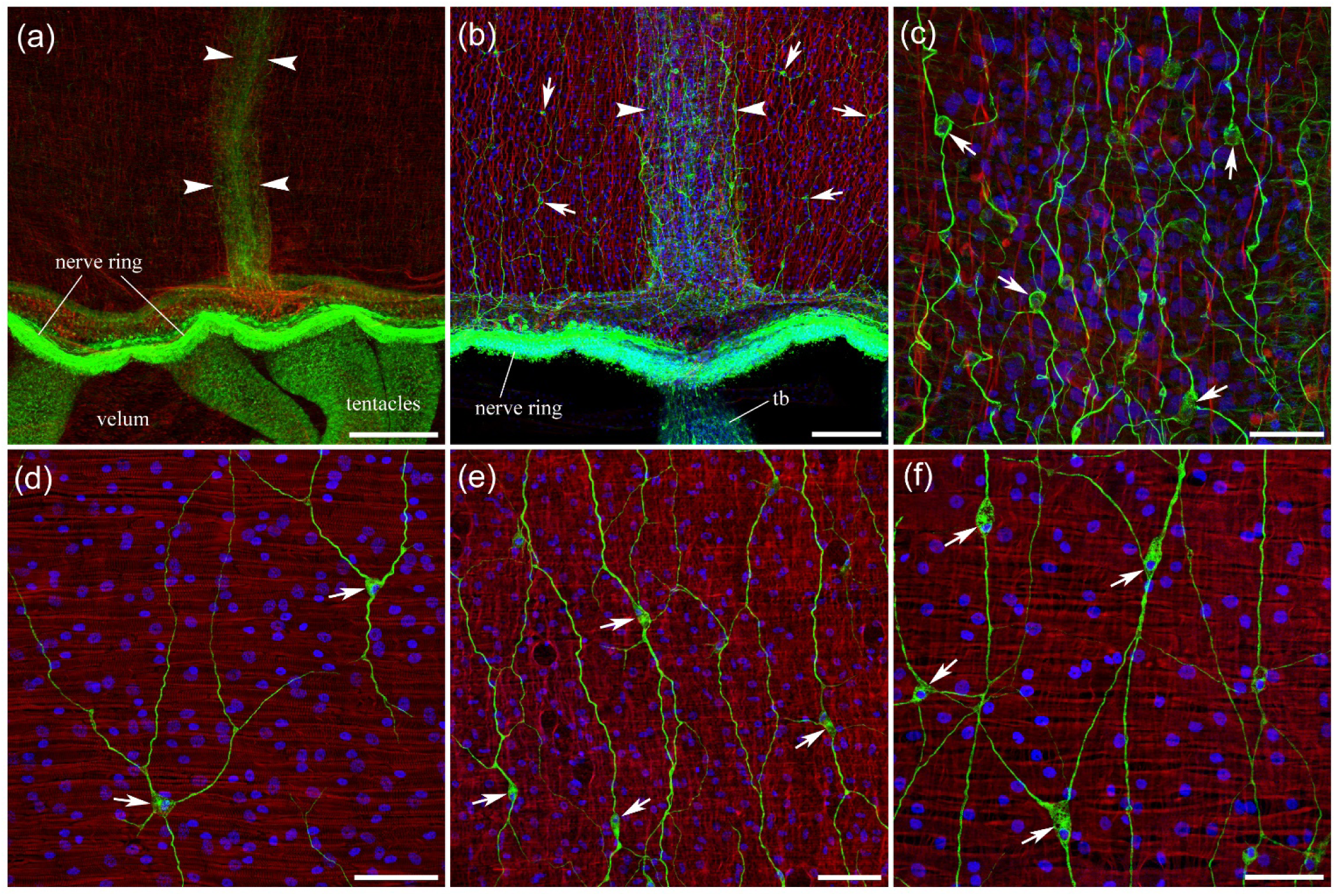
The structure of the *Eutonina* neural and muscle systems in the subumbrella. **a** - The radial neural tract (outlined by arrowheads) originates at the nerve ring and extends across the subumbrella to the apical end of the bell. **b** - The radial neural tract (arrowheads) consists of several mostly longitudinally oriented neurites. In addition, numerous neurons (arrows) are present throughout the entire subumbrella region, forming a neural network. **c** - High magnification of the radial neural tract shows many longitudinally oriented neurites produced by bipolar and tripolar neurons (arrows). **d** - In small adults (1-1.5 cm), subumbrella neurons (arrows) that are distributed throughout the subumbrella are more individually identifiable, although they appear to make visible connections with some of their neighbors. **e, f** - In large adults (up to 4 cm), the subumbrella neurons (arrows) form a fully developed neural network. Note the smooth radial muscle fibers across the entire subumbrella region, running perpendicular to the circular striated muscles. Scale bars: **a** - 200 µm; **b** - 100 µm; **c** - 30 µm; **d, f** - 40 µm; **e** - 60 µm.

Smooth radial muscle fibers are distributed throughout the subumbrella region. They run perpendicular to the circular striated swim muscles (Fig. 25f). The radial smooth muscle fibers do not form tight muscle bundles along the radial canals – as seen in some other hydrozoans (Fig. 25a, b).

The long, conical peduncle in *Eutonina* also contains a fully formed neural network that extends along its entire length (Fig. 26a, b). The network consists of multipolar neurons and their neurites (Fig. 26a, b). Four long gonads along the radial canals contain many small neural cells distributed throughout the length of each gonad (Fig. 26c, d).

**Figure 26.**
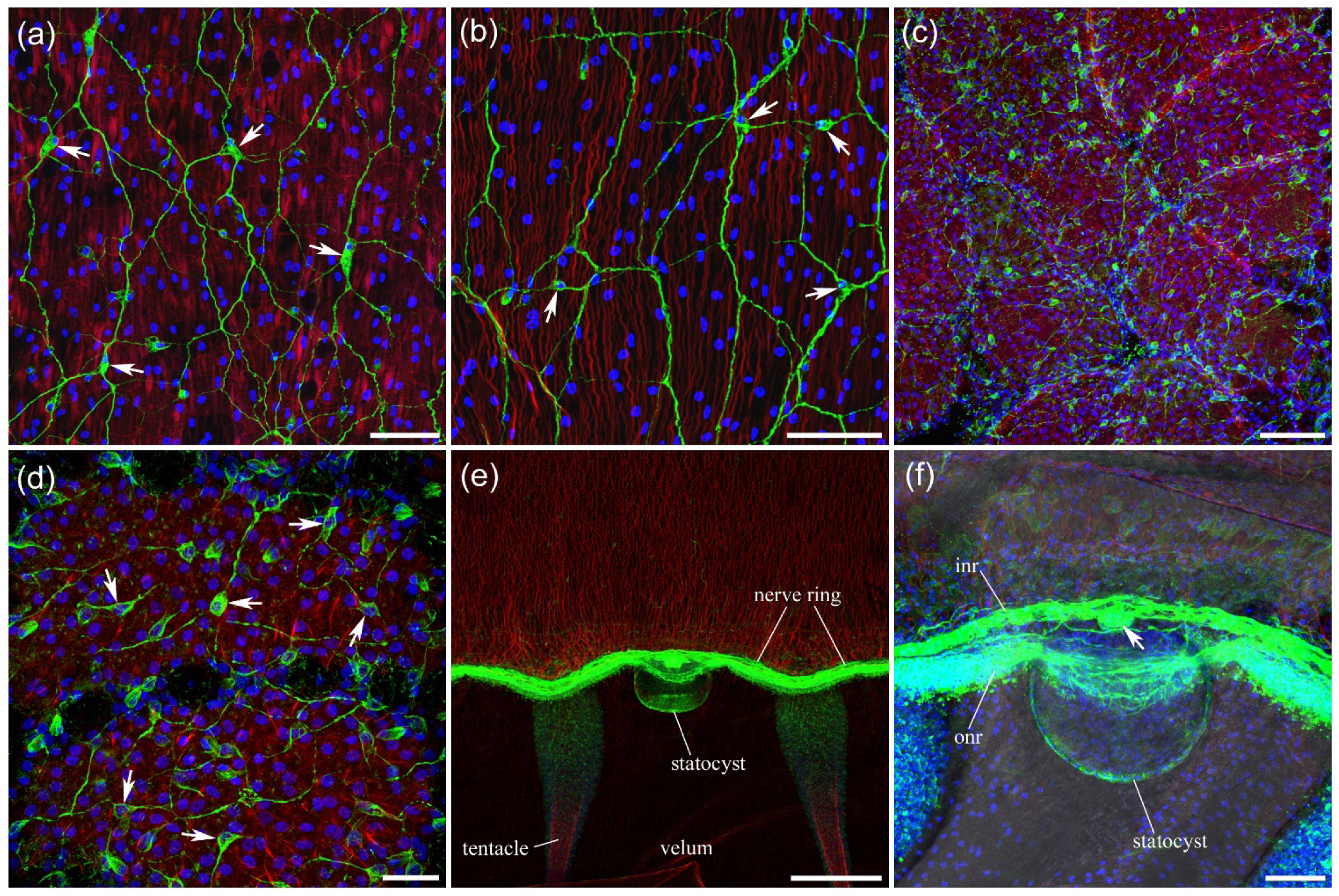
Innervation of the peduncle, gonads, and statocysts in *Eutonina.* **a, b** - The entire peduncle contains a fully developed neural network composed of multipolar neurons (arrows) and their long neurites. **c, d** – Long gonads along the radial canals contain numerous small neurons (arrows) with neurites distributed throughout the length of each gonad. **e, f** - Eight statocysts are located around the bell margin between tentacles protruding on the subumbrella side of the bell. The statocyst vesicle is connected to the inner nerve ring (*inr*), while the outer nerve ring (*onr*) wraps around the vesicle, forming a noticeable arc near the exumbrella surface and away from the statocyst. The arrow in (f) points to the bulging of the inner nerve ring (*inr*) where the connection to the statocyst occurs. The statocyst vesicle is clearly visible in DIC light in (f). Scale bars: **a, b, c, f** - 50 µm; **d** - 20 µm; **e** - 200 µm.

*Eutonina* has eight statocysts located around the bell margin, between the tentacles and protruding from the subumbrella toward the center of the bell (Fig. 26e). The statocyst vesicle is connected to the inner nerve ring, which shows a noticeable bulge at the point of connection (Fig. 26f). The outer nerve ring runs around the vesicle, closer to the exumbrella side of the bell. The vesicle itself contains no labeled neurites.

#### *Eperetmus typus* (order Limnomedusae, family Olindiidae)

Numerous, relatively long tentacles are located along the margin of the bell. Four radial canals have four firmly attached gonads on the subumbrella surface.

The nerve ring is very prominent and brightly labeled with the tubulin AB (Fig. 27a, d). Along the margin of the bell, next to the nerve ring, there is a wide bundle of thick, circular, smooth muscle fibers (Fig. 27a-c). Other hydrozoans also have a bundle of smooth circular muscle fibers at the bell margin next to the nerve ring. However, *Eperetmus* has the largest and thickest bundle among all studied hydrozoans. This layer of smooth circular muscle is extensively innervated, with numerous thin and thick neurites running mostly in the circular direction (Fig. 27a, c).

**Figure 27.**
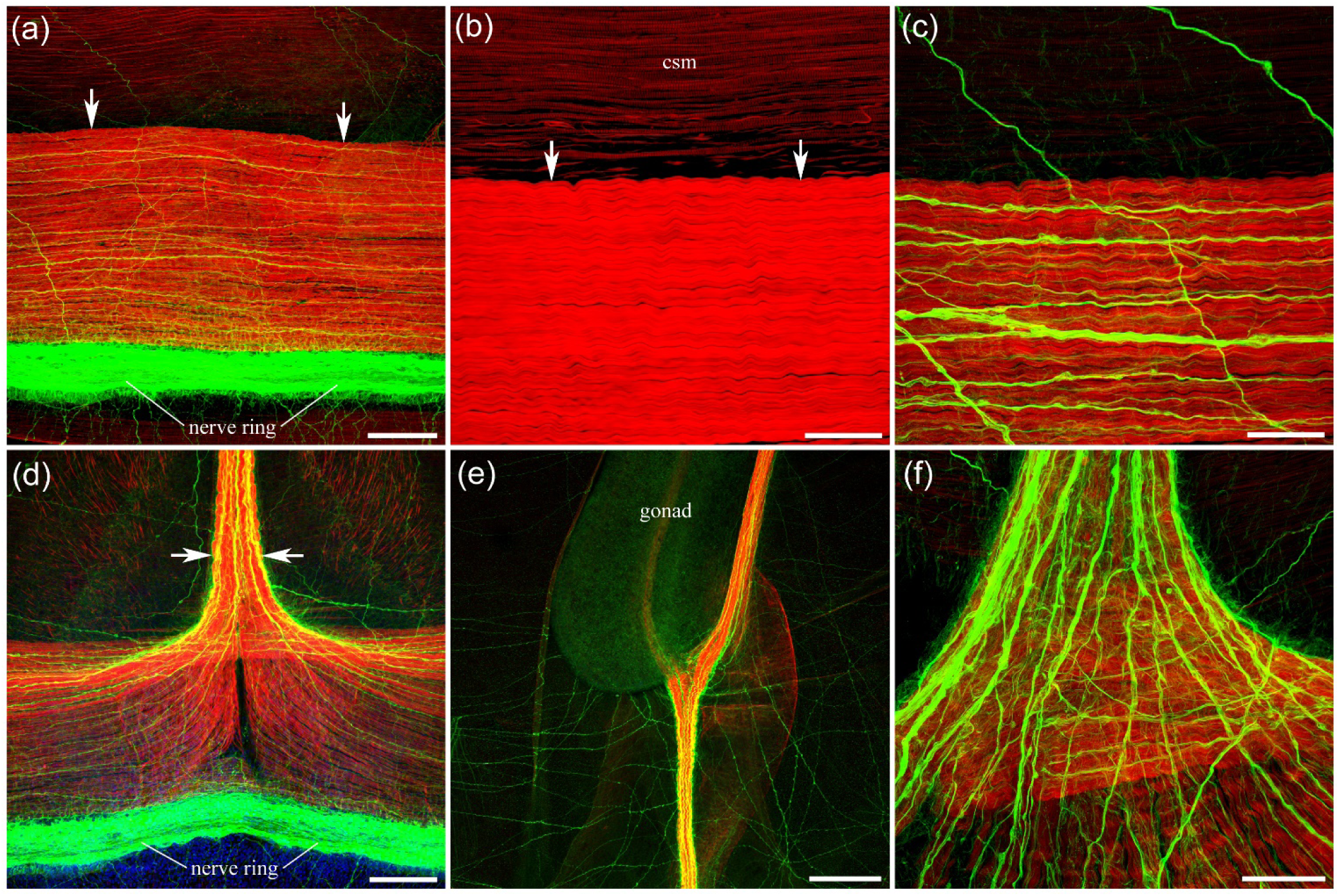
The muscular and neural systems in *Eperetmus*. **a** - A broad band of thick circular smooth muscles (arrows) runs along the margin, adjacent to the nerve ring. **b** - Phalloidin-only labeling (without tubulin AB) of the circular smooth muscle bundle (arrows). Note the striation of the circular striated muscles (*csm*) controlling swimming. **c** - Numerous thick and thin neurites are embedded in this band of circular smooth muscles, running mostly in the same circular direction. **d** - Upon approaching a radial canal from both sides, the circular smooth muscle fibers turn 90° and join to form a radial smooth muscle pathway (arrows) running from the margin to the apical end of the bell. The neural tract within the radial pathway is very pronounced. Some neural fibers that form that tract come directly from the nerve ring, while many others that join the radial neural tract come from the circular smooth muscle band area. **e** - Each radial muscular and neural tract runs to the gonad, which is firmly attached to the subumbrella, where it bifurcates into two branches – each running farther to the apical end of the umbrella on each side of the gonad. **f** - The radial neural tract consists of numerous thick and thin neurites running mostly in the longitudinal, radial direction. Scale bars: **a** - 100 µm; **b, c** - 30 µm; **d** - 100 µm; **e** - 200 µm; **f** - 40 µm.

Four radial pathways run along the radial canals from the bell margin to the apical end of the bell. Each pathway includes a bundle of smooth radial muscles and a radial neural tract (Fig. 27d-f). The circular smooth muscle fibers from the bell margin make a 90° turn and join from two sides to form a radial muscle pathway along the radial canal (Fig. 27d). The radial neural tract, embedded within the bundle of radial smooth muscles, is prominent and consists of numerous thick and thin neurites (Fig. 27f). Some neurites that form the tract arise directly from the nerve ring, while many others that join the radial neural tract follow the circular smooth muscle band as it turns into the radial pathway (Fig. 27d). Each radial pathway runs to the gonad attachment, where it bifurcates into two branches that extend farther to the apical end of the subumbrella on each side of the gonad (Fig. 27e).

Many long neural processes traverse the subumbrella over long distances (Fig. 28a). They run in different directions, though many are circularly oriented. Some long neural processes clearly originate from the radial neural tracts (Fig. 28b). However, neural cell bodies are also present throughout the subumbrella, apparently producing many of these long neural fibers at a distance from the radial neural tracts (Fig. 28c, d). These cell bodies are exclusively bipolar neurons. Importantly, there are no smooth radial muscles in the subumbrella outside the radial pathways.

Many thin and thick neurites branch from the nerve ring and extend into the velum, providing its innervation (Fig. 28e). The tentacles also have a very dense innervation along their entire length. Most neurites run longitudinally, parallel to each other, toward the tip of the tentacle (Fig. 28f). The regularly spaced disks, resembling doughnuts, carry numerous nematocytes (Fig. 28f).

**Figure 28.**
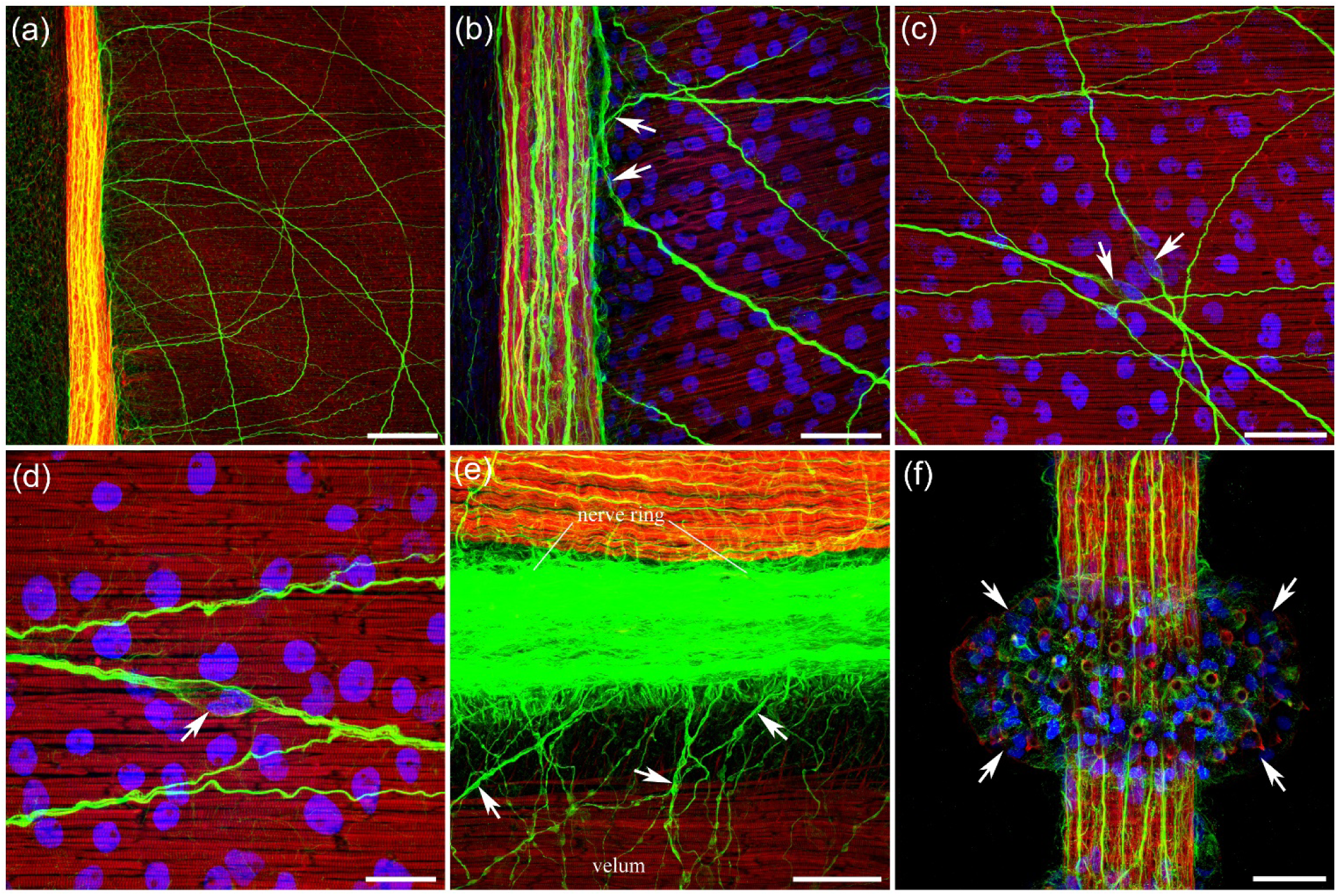
Neural system in *Eperetmus*. **a** - Numerous long neural processes extend outside the radial neural tracts, crossing the entire subumbrella in different directions. **b** - Many of them originate (arrows) from the radial neural tracts. **c, d** - Neural cell bodies that produce these long fibers are also found throughout the subumbrella, at a distance from the radial neural tracts. These cell bodies are exclusively bipolar neurons (arrows). **e** - Numerous neurites (arrows) originating from the nerve ring innervate the velum. The circular muscles in the velum are striated. **f** - The tentacles have very dense innervation throughout their length: mostly longitudinal neurites running in parallel toward the end of the tentacle. The regularly spaced doughnut-shaped disks (arrows) carry nematocytes. Scale bars: **a** - 100 µm; **b, c** - 40 µm; **d** - 20 µm; **e** - 40 µm; **f** - 30 µm.

#### *Aglantha digitale* (order Trachymedusae, family Rhopalonematidae)

There are eight radial canals and eight thin gonads attached to the subumbrella along the canals in the apical part of the bell. Numerous thin tentacles are located along the margin of the bell.

*Aglantha* has a unique system of giant axons that play a key role in swimming behavior (Donaldson et al., 1980; Mackie & Meech, 1985; Roberts & Mackie, 1980; Singla, 1978b). Eight motor giant axons run along the radial canals from the nerve ring to the apex of the bell and are brightly labeled with tubulin AB (Fig. 29a, b; Norekian and Moroz, 2020b). In large animals, the motor giant axons reach up to 35 μm in diameter at their base near the nerve ring. Closer to the apex, they are significantly narrower, in the range of 5–15 μm (Fig. 29c; Norekian and Moroz, 2020b). Near the nerve ring, each motor giant produces short, thick branches that enter the inner nerve ring and then connect via gap junctions to the entire system of rootlet interneurons, which spread around the nerve ring on either side of each motor giant to make contact with their neighbors (Mackie & Meech, 2000; Weber et al., 1982). The nerve ring is the source that drives the activity of motor giant axons during both slow and fast swimming in *Aglantha* (Meech & Mackie, 1993a, 1995; Roberts & Mackie, 1980). In addition to motor giant axons, the subumbrella area contains a number of lateral neurons electrically coupled to axons (Weber et al., 1982). Lateral neurons run parallel to the circular striated muscles and perpendicular to motor giants at more or less evenly spaced intervals (Fig. 29c; Norekian and Moroz, 2020b). The lateral neurons branch extensively and sometimes overlap, covering and innervating much of the circular striated muscle sheet, as they are in the same focal plane where striated muscle fibers are located (Fig. 29d; Norekian and Moroz, 2020b). Giant motor axons, directly and via a population of lateral motor neurons, activate striated muscle cells of the subumbrella myoepithelium (Kerfoot et al., 1985).

**Figure 29.**
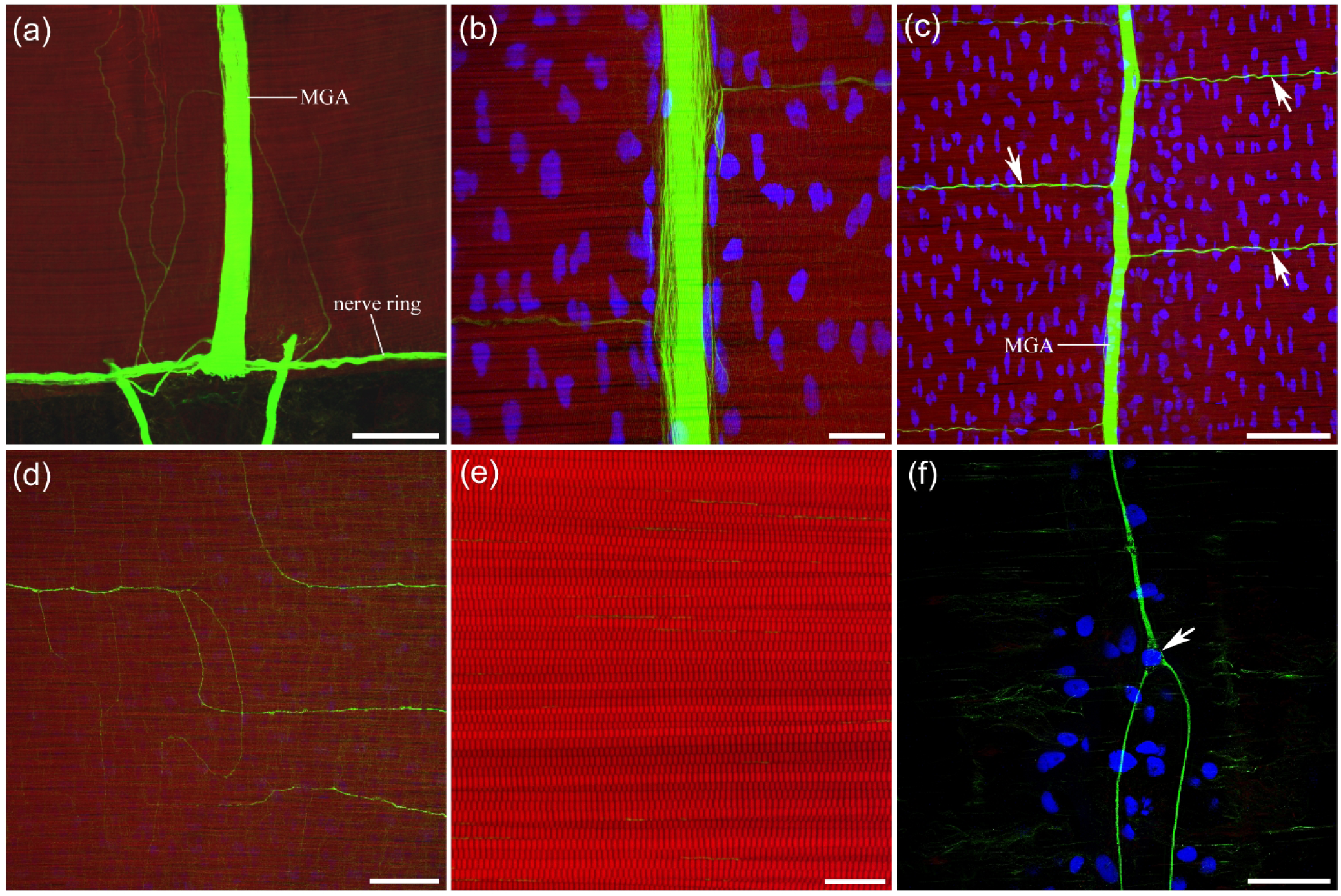
Neural and muscle systems in the subumbrella of *Aglantha*. **a** - Motor giant axons (*MGA*s) run across the entire umbrella along radial canals from the nerve ring at the margin to the apex. **b** - Each motor axon is a giant syncytium, with tubulin AB labeling numerous thin tubulin filaments inside the axon, visible at higher magnification. **c** - Along their entire length, MGAs connect to many lateral neurons (arrows) that run perpendicular to the giants, in parallel with the striated muscle fibers and more or less evenly distributed across the subumbrella. **d** - Lateral neurons branch extensively in the striated muscle layer, with neurons from neighboring MGAs sometimes having overlapping innervation fields. **e** - The striation of the circular striated muscles controlling swimming is clearly visible at high magnification. **f** - The walls of the radial canals contain a few large tripolar neurons (arrow) with long processes that run longitudinally along the radial canal. Scale bars: **a** - 100 µm; **b** - 20 µm; **c** - 50 µm; **d** - 100 µm; **e** - 10 µm; **f** - 40 µm.

The walls of the radial canals contain a few large tripolar neurons (Fig. 29f; Norekian and Moroz, 2020b). These neurons produce long neurites that run radially along the length of the radial canal. Usually, the radial canal and these tripolar neurons, with their long neurites, are masked by the bright tubulin immunostaining of the motor giant axons. However, in some preparations, the motor giant axons can be slightly displaced, revealing these radial canal neurons.

The nerve ring consists of an inner nerve ring located closer to the subumbrella surface and an outer nerve ring with a giant axon located closer to the tentacles (Mackie & Meech, 1995a, 1995b; Roberts & Mackie, 1980; Singla, 1978b). Both the inner and outer nerve rings are visible in tubulin-immunolabeled preparations. In addition to the eight giant motor axons that connect to the nerve ring, the nerve ring gives rise to many thin neural processes that form a basal plexus around the ring.

#### *Solmundaegina nematophora* (previously described as *Aegina citrea*; order Narcomedusae, family Solmundaeginidae)

*Solmundaegina* has four thick, muscular tentacles located not at the margin of the bell, as in other hydrozoans, but are elevated closer to the apical end of the bell. Each tentacle has a thick, curved “root” deeply embedded in the umbrella. At the midsection, there is a row of 8 wide pouches involved in digestion. It is a very active, fast swimmer and an aggressive predator, hunting other jellies of different sizes, even as big as *Solmundaegina* itself. We saw it attack a *Stomotoca* of equal size, grabbing it with all four tentacles and looking like a spider on top of its prey.

The nerve ring is thick, clearly defined, and brightly labeled with tubulin AB, as in other hydrozoan species (Fig. 30a, b). The main difference, which makes *Solmundaegina*’s nervous system particularly unique, is the presence of four very thick branches (the same thickness as the nerve ring itself) that run from the nerve ring to the base of each tentacle – tentacular nerves (Fig. 30a, b). Several large axons and smaller neurites exit the nerve ring on both sides of the nerve, make a 90° turn toward the apical end of the bell, and merge to form a thick tentacular nerve (Fig. 30b, ci). A tight bundle of smooth radial muscle fibers, intensely labeled by phalloidin, is associated with the tentacular nerve and runs with it up to the tentacle base (Fig. 30cii, d). A similar narrow bundle of smooth muscle also runs along the nerve ring around the bell margin.

**Figure 30.**
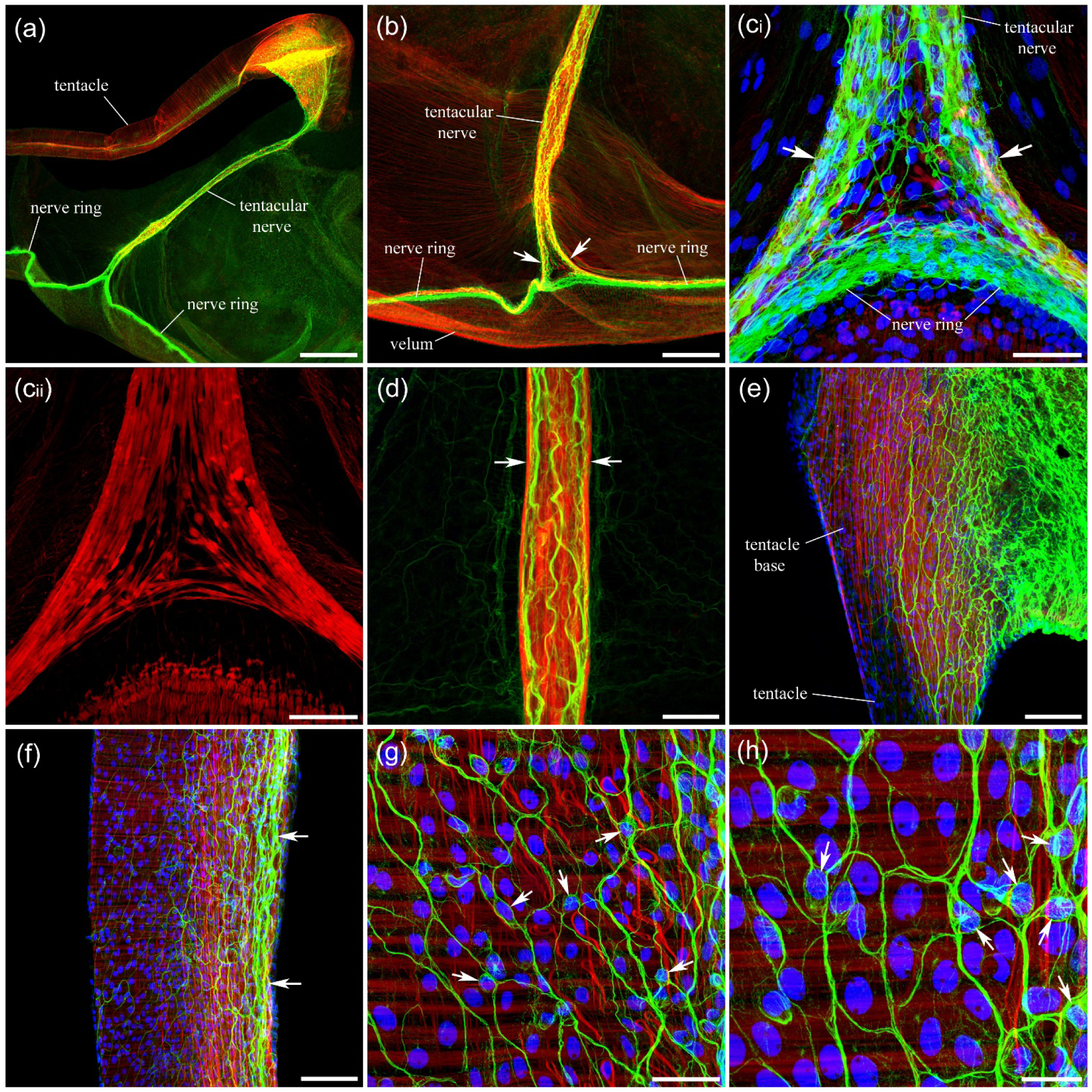
Tentacular nerves and innervation of the tentacles in *Solmundaegina*. **a** - A thick tentacular nerve runs perpendicular to the bell margin from the nerve ring across the umbrella to the tentacle base, where it extensively branches, forming a dense neural network. **b, c_i_** - Each tentacular nerve is formed by two thick bundles of neurites (arrows) that exit the nerve ring, make a 90° turn toward the tentacle base, and merge. **c_ii_** - Phalloidin labels smooth muscle fibers in the tentacular nerve and nerve ring (this is the red phalloidin channel from **c_i_**). **d** - The tight smooth muscle bundle runs within the tentacular nerve (arrows) all the way to the tentacle base. **e** - A very dense neural network is present at the tentacle base, which then extends into the tentacle. **f** - The neural network runs the entire length of the tentacle, concentrating mostly on the inward side (arrows) of the tentacle facing the umbrella. Note that longitudinal smooth muscle fibers are also localized to the same inward side of the tentacle. **g, h** - The neural network in the tentacles is formed by numerous multipolar neurons, whose cell bodies (arrows) are identified by DAPI labeling of their nuclei. Scale bars: **a** - 500 µm; **b** - 200 µm; **c, h** - 40 µm; **d** - 50 µm; **e, f** - 100 µm; **h** - 20 µm.

At the base of each tentacle, the tentacular nerve extensively branches to form a dense neural network (Fig. 30a, e). This network then extends into the tentacle itself (Fig. 30f). The tentacle musculature consists of circular muscles and longitudinal smooth muscle fibers (Fig. 30f-h). Most of the smooth longitudinal muscle fibers are located on the inward side of the tentacles, facing the bell (Fig. 30f). This arrangement apparently causes the tentacles to bend and grab the prey. The neural network is also much denser on that inward side, where the longitudinal muscle fibers are located (Fig. 30f). The network is formed by numerous multipolar neurons with cell body sizes 10-15 µm (Fig. 30g, h). It extends along the entire length of the tentacle, becoming narrower and less dense toward the tentacle end.

The circular striated muscle layer is relatively narrow and lies between the bell margin and a row of 8 thick digestive pouches (Fig. 31a). Thin radial smooth muscle fibers are also present in this area. A noticeable neural mesh extends from the middle part of the tentacular nerve into the band of striated circular muscles in its upper region (Fig. 31b). The striated muscle area closer to the pouches has a relatively dense network of neural fibers, many of which are circularly oriented and presumably originate from the tentacular nerve (Fig. 31b). In addition, a loose mesh of neural cell bodies and their long neurites is distributed throughout the entire striated muscle band area (Fig. 31c). This mesh is not very dense but is well pronounced. The long neurites crossing in all directions are produced mostly by tripolar and bipolar neurons (Fig. 31c, d).

**Figure 31.**
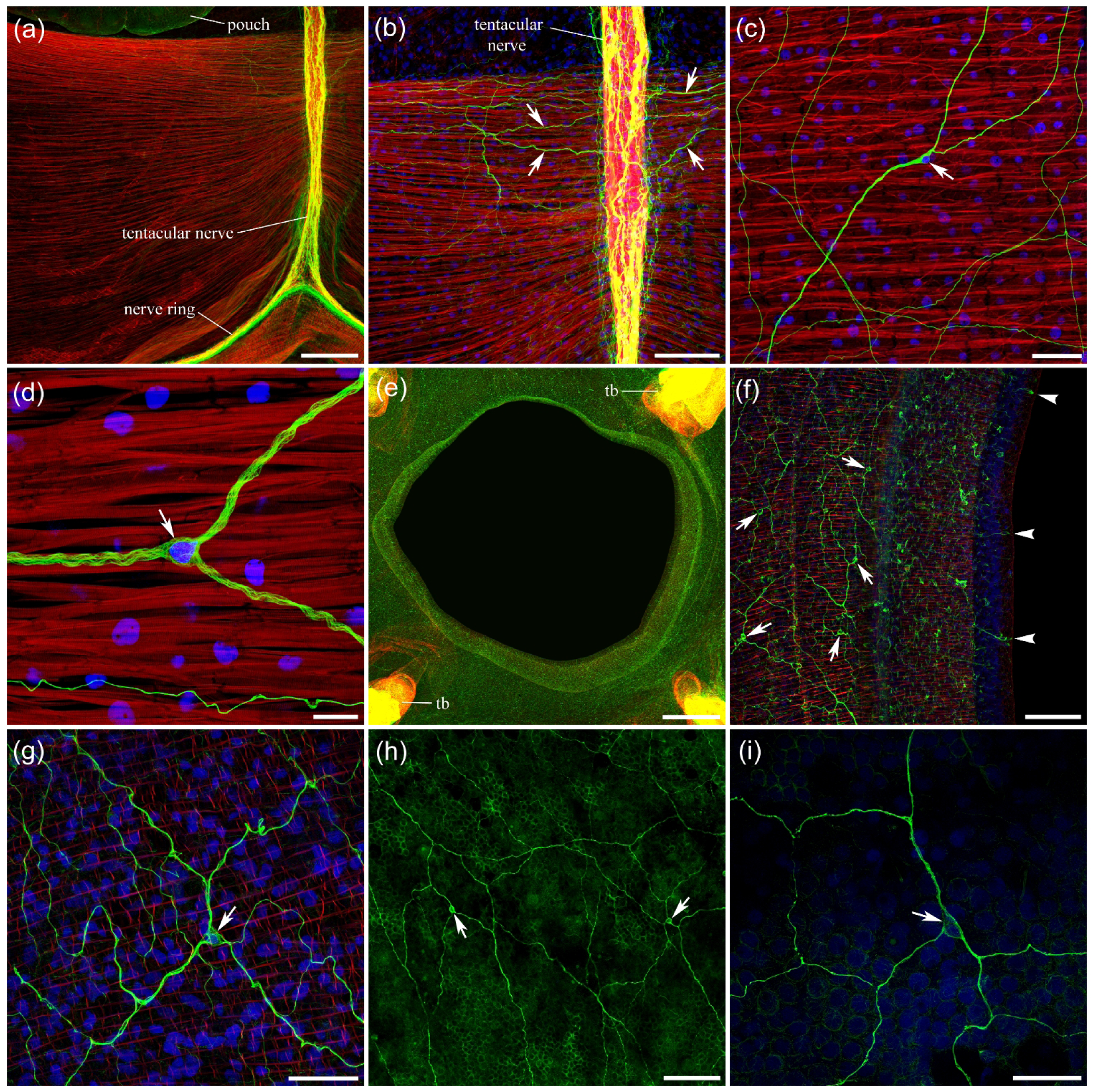
Neural and muscle systems in the subumbrella of *Solmundaegina*. **a** - The circular striated muscle layer is confined to the region between the digestive pouches and the nerve ring at the bell margin, running around the lower part of the subumbrella from one tentacular nerve to another. **b** - Long neurites (arrows) extend from the tentacular nerve to the band of circular striated muscles, mostly at its upper side, closer to the digestive pouches. **c** - A scattered mesh of neurons (arrow) with very long neurites running in all directions is located within the striated muscle area. **d** - These neurons are mostly tripolar (arrow) and bipolar cells. **e** - A large mouth that resembles a diaphragm is located inside the bell at the level of the tentacle bases (*tb* - one at each corner of the image). **f** - The mouth area is densely innervated by a network of neurons (arrows) labeled with tubulin AB. Arrowheads point to the presumably sensory neurons at the margin of the mouth. **g** - Neurons that form the network in the mouth area are mostly multipolar cells (arrow) with extensive branching. **h, i** - The wall of the digestive pouches contains neurons (arrows) with long, branching neurites forming a neural network, although at much lower density than in the mouth area. Scale bars: **a** - 200 µm; **b** - 100 µm; **c** - 50 µm; **d** - 20 µm; **e** - 500 µm; **f, h** - 100 µm; **g, i** - 40 µm.

*Solmundaegina* lacks a manubrium hanging from the apical roof of the subumbrella, as in other hydrozoans. It has a large mouth at the level of the tentacle bases, resembling a diaphragm – a large opening surrounded by a flat muscular ring (Fig. 31e). The mouth leads to a large cavity, apparently where prey is placed for digestion. This large central cavity extends into all 8 smaller digestive pouches. The entire mouth area has a dense neural network labeled with tubulin AB (Fig. 31f). This neural network consists of numerous small multipolar neurons with long, extensively branching neurites (Fig. 31f, g). There are also small, elongated, presumably sensory neurons at the margin of the mouth (Fig. 31f).

The neural network is also present in the walls of the digestive pouches (Fig. 31h). It is less dense than the neural network in the mouth area and is more prevalent in the upper parts of the pouches, closer to the mouth. The network consists of small multipolar neurons with very long, branching neurites (Fig. 31h, i).

Narcomedusae have unusual round nematocysts, characteristic of this order (Mackie and Mackie 1963; Mills and Miller 1984). Even the name of the current species, *Solmunaegina nematophora*, alludes to the nematocyst patches on the exumbrella (phora = bearing in Latin; Lindsay et al., 2017). Clusters of 15 to 50 round nematocysts are distributed across the exumbrella surface and are clearly visible under DIC light (Fig. 32a). Each nematocyst has a ring of microvilli, brightly labeled with phalloidin (Fig. 32b). In the center of this microvilli ring is a sensory cilium called the cnidocil, which is weakly labeled with the tubulin AB (Fig. 32c).

**Figure 32.**
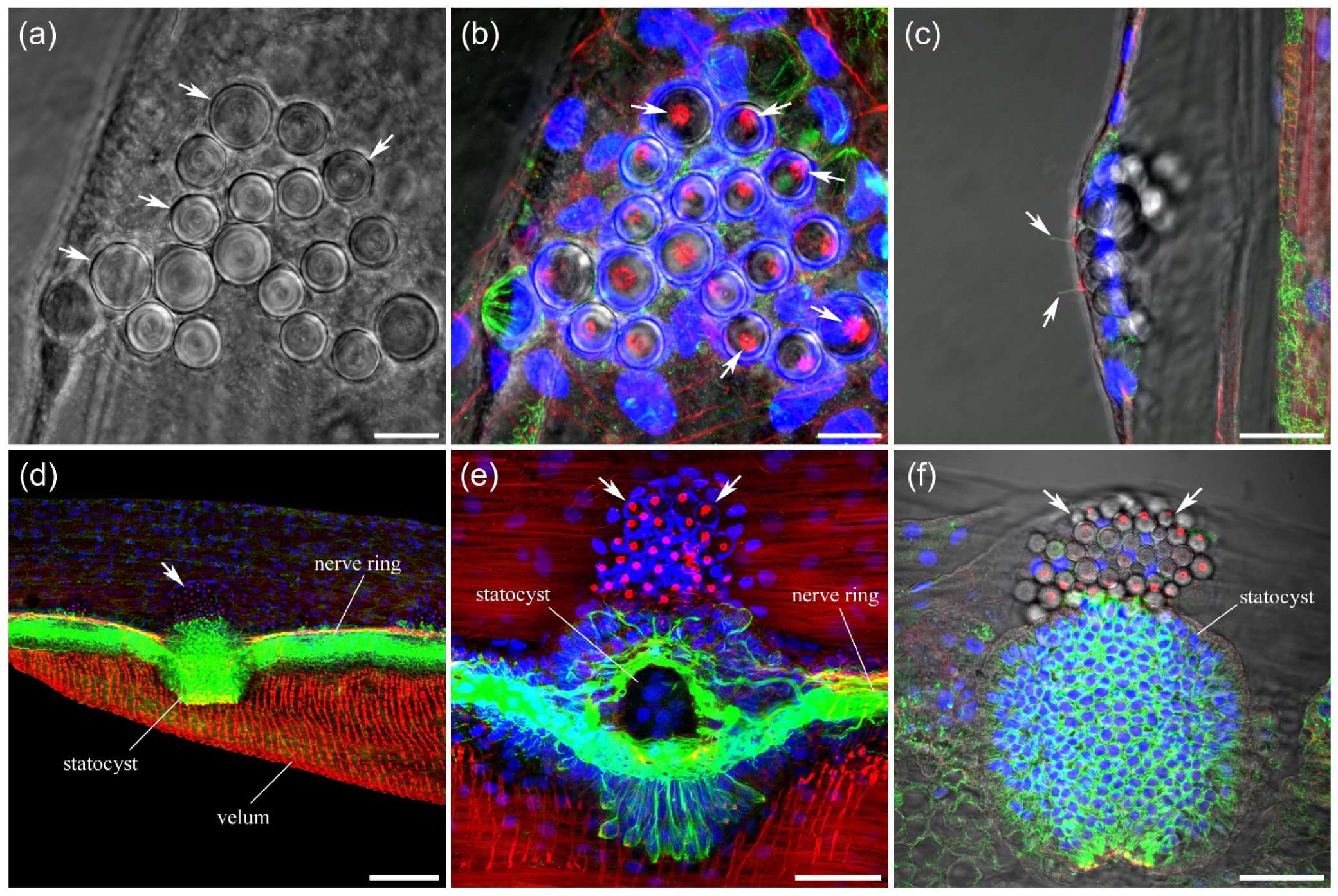
Nematocysts in *Solmundaegina*. **a** - A cluster of round nematocysts (arrows) in the exumbrella under DIC light. **b** - Phalloidin labels a ring of short stereocilia (arrows) on each nematocyst. Note a single large nucleus associated with each nematocyst, labeled with DAPI. **c** - Each nematocyst has a single sensory cilium, the cnidocil (arrows), which is weakly labeled with the tubulin AB. **d** - A cluster of nematocysts (arrow) is always associated with each statocyst (view from the bottom of the bell up). Several statocysts are located around the bell margin and connected to the nerve ring. Note that the velum contains radial and circular muscle fibers. **e** - An optical section through the statocyst reveals its tight connection to the nerve ring. A cluster of nematocysts (arrows) is always present next to the statocyst. **f** - A statocyst with a cluster of nematocysts (arrows) under DIC light. Scale bars: **a, b** - 10 µm; **c** - 20 µm; **d** - 100 µm; **e, f** - 40 µm.

There are several statocysts in *Solmunaegina* attached to the bell margin, next to the nerve ring (Fig. 32d). Each statocyst has a cluster of nematocysts associated with it on the exumbrella side of the margin (Fig. 32d-f). Statocysts are linked to the nerve ring, and some neurites appear to branch from the nerve ring to encircle the statocyst, presumably contributing to its innervation (Fig. 32e).

## 4 DISCUSSION

Satterlie (2008) suggested that in *Aequorea*, swim contractions are controlled not only by synaptic input from the motor network of the inner nerve ring and electrotonic depolarization from adjacent active muscle cells, but also by synaptic inputs from the subumbrella nerve net, which further conducts excitation from the inner ring motor network to the swim muscles. However, comparative investigations in a number of other hydromedusae suggest that similar subumbrella neural networks, which show RFamide immunoreactivity, are associated with radial smooth muscles rather than the circular swimming system (Mackie, Singla & Stell, 1985). We have not done electrical recordings to prove or disprove this theory. Nevertheless, all our morphological data strongly suggest a close link between the smooth radial muscles and the subumbrella neural networks. These radially oriented smooth muscles are used for widespread defensive responses and local radial feeding and righting responses, while only striated muscles with a circular orientation produce swim contractions. In almost all species studied, the distribution of smooth radial muscles matched the distribution of neural networks in the subumbrella. For example, *Polyorchis, Sarsia, Euphysa, Stomotoca*, and *Bougainvillia* all have bundles of radial smooth muscles along the radial canals and no radial smooth muscles in the subumbrella outside those radial bundles (Table 1, Fig. 33). And they all have only the radial neural tracts along the radial canals in the same area, with no neural elements between those radial pathways. At the same time, *Neoturris, Aequorea, Mitrocoma, Clytia*, and *Eutonina* all have radial smooth muscle fibers spread throughout the entire subumbrella region (Table 1, Fig. 33). And they all have neural networks present throughout the entire subumbrella, not only along the radial canals. *Leuckartiara* is particularly interesting, as it further confirms this pattern. *Leuckartiara* does not have a tight bundle of radial smooth muscles as other species do, but has a wide lane around each radial canal that contains smooth radial muscle fibers with large voids between those lanes. Its neural network is restricted exactly to these wide lanes. This strong correlation between the location of radial smooth muscles and diffuse neural networks in the subumbrella suggests that these diffuse subumbrella neural networks control radial smooth muscles in hydrozoans.

**Figure 33.**
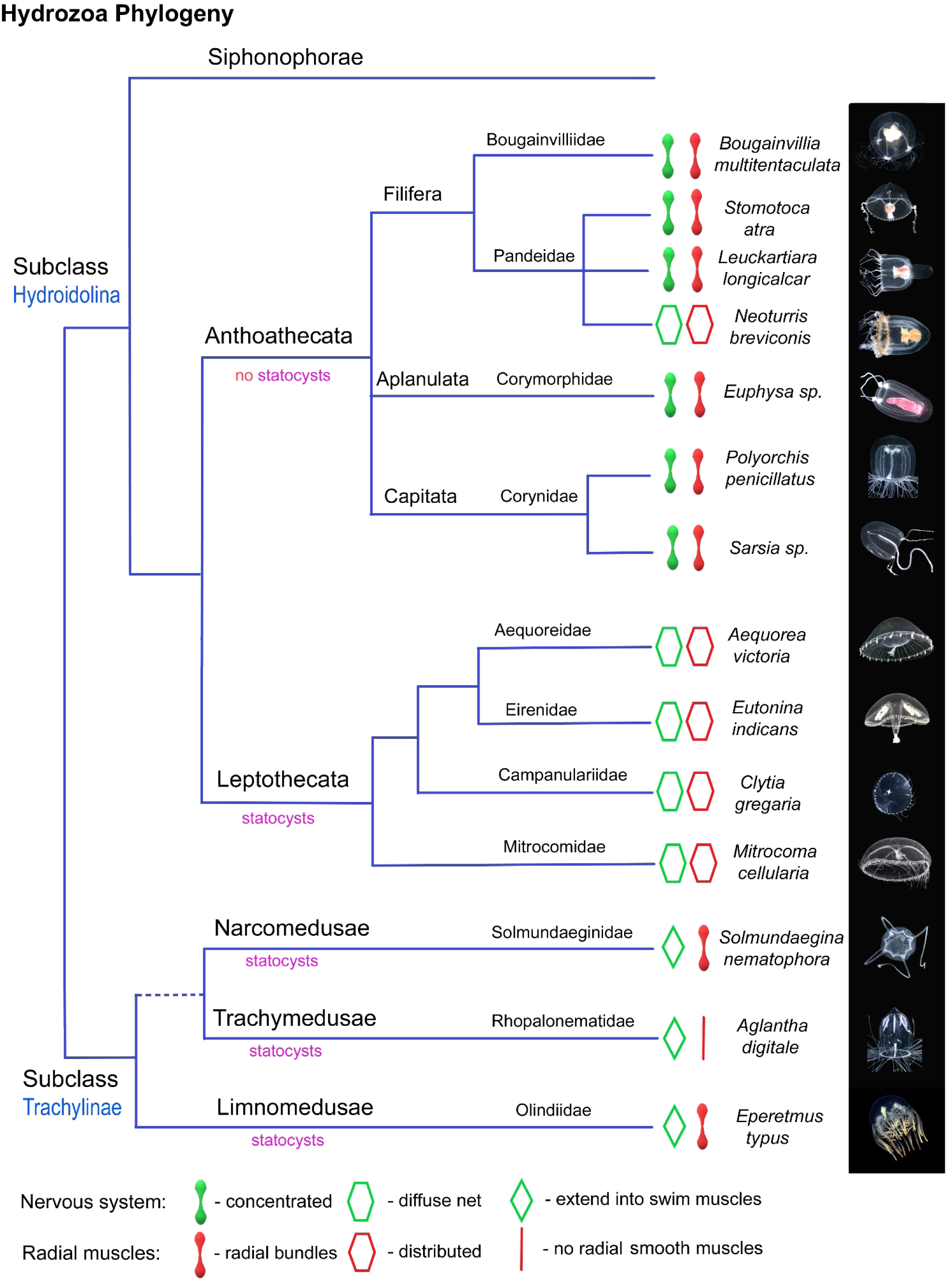
Phylogenetic relationships of studied hydrozoans and architecture of their neural and muscle systems in the subumbrella. Nervous systems can be concentrated in the narrow radial tracts along the radial canals, or cover the entire subumbrella as a uniform diffuse neural network. Smooth radial muscles can be located only in tight radial bundles along the radial canals, or be distributed throughout the subumbrella region. Note that there is a close match between the distribution of neural systems and distribution of radial smooth muscles. There is also a clear tendency to have a diffused and uniformly distributed neural and radial muscle systems in the order Leptothecata, and concentrated distribution of neural and radial muscle systems in the radial tracts along the radial canals in the order Anthoathecata (*Neoturris* being a noticeable exception). In *Aglantha*, the radial smooth muscles are absent, and the neural system specifically targets the swimming striated muscles. In *Eperetmus* and *Solmundaegina*, neural processes also extend into subumbrella regions between radial canals where only swim muscles are located.

Three species from the list stand out and differ from the others. First is *Aglantha*, which is unique among hydrozoans. *Aglantha* lacks radial smooth muscles in the subumbrella, so it does not perform defensive "crumpling" behavior. Its subumbrella nervous system is entirely dedicated to controlling the swim striated muscles and includes giant motor axons along the radial canals (instead of radial neural tracts) and lateral neurons that innervate the swim muscles. *Solmundaegina* is also very different in its anatomy and behavior, with four very thick, muscular tentacles located not at the bell margin but on the top of the bell, used to physically grab large prey. Its radial neural pathways also function as tentacular nerves that connect the marginal ring nerve to the densely innervated tentacles. These radial neural pathways/tentacular nerves are tightly intertwined with bundles of thick smooth radial muscles and presumably innervate them, as in other hydrozoan species. However, the neural cells and processes are also present in the subumbrella while radial muscles are not (Table 1, Fig. 33). *Eperetmus* is similar to the first group of hydrozoan species, with radial smooth muscles concentrated only in the radial pathways, associated with radial neural tracts (Table 1, Fig. 33). However, *Eperetmus* also has numerous neural processes that cross the entire subumbrella, where smooth radial muscles are not present. Notably, these processes are very long, span large distances, and do not resemble a local neural network. Why do *Solmundaegina* and *Eperetmus* have neural processes in the subumbrella regions that lack smooth radial muscles? It is possible that those processes target the swim striated muscles in these two species, similar to *Aglantha*. It is important to stress that *Aglantha* is the fastest swimmer among all species studied. *Solmundaegina* is also very agile and shows fast directional swimming. *Eperetmus* is also very fast. This fast swimming may be linked to neural processes in the subumbrella, which could provide an additional boost to the swimming system in these two species, as in *Aglantha*.

Why do some hydrozoan species have smooth radial muscles and neural elements concentrated only around radial canals, forming narrow radial pathways, while others have smooth radial muscles and diffuse neural networks spread throughout the entire subumbrella region? Satterlie (2018) suggested that the appearance of the subumbrella network might be correlated with the oblate (broad and shallow) versus prolate (elongated and narrow) umbrella type in hydrozoan medusae. For example, oblate species such as *Aequorea, Clytia*, and *Eutonina* have a diffuse neural network in the subumbrella, whereas prolate species, such as *Polyorchis*, lack a subumbrella network. It appears that many studied species follow that pattern. However, there are notable examples that clearly fall outside this theory. In addition to prolate *Aglantha*, which can be considered a distinct and special arrangement, a fully developed polygonal neural network is found in the subumbrella of prolate *Neoturris*. Thus, the umbrella type is probably not the determining factor. What we can conclude, however, is that the transition between the dispersion and concentration of radial smooth muscles and neural networks in the subumbrella region is a fluid process and was not set in stone evolutionarily long ago. Even closely related species from one family can have completely different types of neural and smooth muscle architecture. Take, for example, three species from the family Pandeidae (order Anthoathecata): *Stomotoca, Neoturris*, and *Leuckartiara*. *Stomotoca* has a concentrated architecture type with radial neural tracts and bundles of smooth muscles along the radial canals only, with no neural elements or radial smooth muscles outside those pathways. *Neoturris* has a diffuse polygonal network and smooth radial muscles spread throughout the entire subumbrella region. And *Leuckartiara* is positioned somewhere in between – with a wide area around the radial canals that contains a neural network and smooth radial muscles, with large voids in between. In general, however, many species from the order Anthoathecata show a tendency to have a concentration of neural elements and smooth muscles only in the radial pathways along the radial canals and lack neural networks and radial smooth muscle fibers in the rest of the subumbrella region (Fig. 33). All studied species from the order Leptothecata, by contrast, have diffuse neural networks and radial smooth muscle fibers spread throughout the entire subumbrella region (Fig. 33).

The advantage of having the smooth radial muscles and controlling neural networks spread over the entire subumbrella region is a higher flexibility and ability to produce small precise umbrella movements during defense or feeding responses – for example, local movements of small patches of the bell margin with tentacles that captured the prey. In the case of complete concentration of all smooth radial muscles and subumbrella neural elements in the radial pathways along the four radial canals such small deliberate responses are not possible. Only much larger and less precise umbrella movements are feasible. However, these hydrozoan species may rely on different strategies – escape swimming as a main defense behavior and using long manubrium for prey targeting instead of marginal flexibility. An extreme version is *Aglantha*, which lacks any smooth radial muscles in subumbrella and has the fastest escape response among known hydrozoans. This variety of different behavioral strategies finds its reflection in the diversity of neural network and radial smooth muscle distributions in the hydrozoan subumbrella region. Thus, having the standard body plan similar in all hydrozoans and very standard ring nerve system that initiates the swim rhythm and controls swim muscles, different hydrozoan species show a significant diversity in the subumbrella neural network and radial muscle architecture, which reflects the diversity of their behaviors and ecology.

## Acknowledgments

We thank FHL for its excellent microscopy facilities. We also thank Dr. Claudia Mills for assistance in identifying hydrozoan species. This research was supported by the National Science Foundation grant (IOS-2341882) and National Institutes of Health grant (5R01NS11449) to LLM.

## Data Availability Statement

The data that support the findings of this study are available from the corresponding author s upon request.

## Role of the authors

All authors take responsibility for the integrity of the data and the accuracy of the data analysis. Research design: TPN. Acquisition of data: TPN. Analysis and interpretation of data: TPN, LLM. Drafting of the article: TPN, LLM. Funding: LLM.

## REFERENCES

Anderson, P.A.V. (1979). Ionic basis of action potentials and bursting activity in hydromedusan jellyfish *Polyorchis penicillatus*. Journal of Experimental Biology, 78, 299–302.

Anderson, P.A.V. & Mackie, G.O. (1977). Electrically coupled, photosensitive neurons control swimming in a jellyfish. Science, 197 (*4299*), 186–188. http://www.jstor.org/stable/1744837

Arkett, S.A., Mackie, G.O., & Meech, R.W. (1988). Hair cell mechanoreception in the jellyfish *Aglantha digitale*. Journal Experimental Biology, 135, 329–342.

Arkett, S.A. & Spencer, A.N. (1986a). Neuronal mechanisms of a hydromedusan shadow reflex. I. Identified reflex components and sequence of events. Journal of Comparative Physiology A, 159, 201–213.

Arkett, S.A. & Spencer, A.N. (1986b). Neuronal mechanisms of a hydromedusan shadow reflex. II. Graded response of reflex components, possible mechanisms of photic integration, and functional significance. Journal of Comparative Physiology A, 159, 215–225.

Donaldson, S., Mackie, G.O., & Roberts, A. (1980). Preliminary observations on escape swimming and giant neurons in *Aglantha digitale* (Hydromedusae: Trachylina). Canadian Journal of Zoology, 58, 549–552. doi: 10.1139/z80-076

Eakin, R.M., & Westfal, l J.A. (1962). Fine Structure of photoreceptors in the hydromedusan, *Polyorchis penicillatus*. Proceedings of the National Academy of Sciences U S A, 48, 826–833. doi: 10.1073/pnas.48.5.826

Gladfelter, W.B. (1972). Structure and function of the locomotory system of *Polyorchis mont*ereyensis (Cnidaria, Hydrozoa). Helgoland Marine Research, 23, 38–79. doi: 10.1007/BF01616310

Gladfelter, W.G. (1973). A comparative analysis of the locomotory systems of medusoid Cnidaria. Helgolander Wiss. Meeresunters 25, 228–272. 10.1007/BF01611199

Harvey, E.N. (1921). Study on Bioluminescence. Biological Bulletin 41, 280–287.

Jha, R.K. & Mackie, G.O. (1967). The recognition, distribution and ultrastructure of hydrozoan nerve elements. Journal of Morphology, 123, 43–62.

Kerfoot, P.A, Mackie, G.O., Meech, R.W., Roberts, A., & Singla, C.L. (1985). Neuromuscular transmission in the jellyfish *Aglantha digitale*. Journal of Experimental Biology, 116, 1–25.

Lin, Y.C., Gallin, W.J., & Spencer, A.N. (2001). The anatomy of the nervous system of the hydrozoan jellyfish, *Polyorchis penicillatus*, as revealed by a monoclonal antibody. Invertebrate Neuroscience, 4, 65–75. doi: 10.1007/s101580100008

Lindsay, D.J., Grossmann, M.M., Bentlage, B., Collins, A.G., Minemizu, R., Hopcroft, R.R., Miyake, H., Hidaka-Umetsu, M. & Nishikawa, J. (2017): The perils of online biogeographic databases: a case study with the‘monospecific’ genus *Aegina* (Cnidaria, Hydrozoa, Narcomedusae). Marine Biology Research, 13(*5*), 494–512. 10.1080/17451000.2016.1268261

Mackie, G.O. (1965). Conduction in the nerve-free epithelia of siphonophores. American Zoologist, 5, 439–453.

Mackie, G.O. (1970). Neuroid conduction and the evolution of conducting tissues. The Quarterly Review of Biology, 45(*4*), 319–332.

Mackie, G.O. (1971). Neurological complexity in medusae: A report of central nervous organization in *Sarsia*. Actas del 1’ Simposio Internacional de Zoofilogenia, Salamanca, pp. 269–280.

Mackie, G.O. (1975). Neurobiology of *Stomotoca*. II. Pacemakers and conduction pathways. Journal of Neurobiology, 6(*4*), 357–378.

Mackie, G.O., & Meech, R.W. (1985). Separate sodium and calcium spikes in the same axon. Nature, 313, 791–793. doi: 10.1038/313791a0

Mackie, G., & Meech, R. (1995a). Central circuitry in the jellyfish *Aglantha*. I: The relay system. Journal of Experimental Biology, 198, 2261–2270.

Mackie, G., & Meech, R. (1995b). Central circuitry in the jellyfish *Aglantha*. II: The ring giant and carrier systems. Journal of Experimental Biology, 198, 2271–2278.

Mackie, G.O., & Meech, R.W. (2000). Central circuitry in the jellyfish *Aglantha digitale*. III. The rootlet and pacemaker systems. Journal of Experimental Biology, 203, 1797–1807.

Mackie, G.O. & Meech, R.W. (2008). Nerves in the endodermal canals of hydromedusae and their role in swimming inhibition. Invertebrate Neuroscience, 8, 199–209. 10.1007/s10158-008-0082-6

Mackie, G.O., Meech, R.W., & Spencer, A.N. (2012). A new inhibitory pathway in jellyfish *Polyorchis penicillatus*. Canadian Journal of Zoology, 90, 172–181. doi: 10.1139/z11-124

Mackie, G.O. & Passano, L.M. (1968). Epithelial conduction in hydromedusae. The Journal of General Physiology, 52, 600–621.

Mackie, G.O. & Singla, C.L. (1975). Neurobiology of *Stomotoca*. I. Action systems. Journal of Neurobiology, 6(*4*), 339–356.

Mackie, G.O., Singla, C.L. & Stell, W.K. (1985). Distribution of nerve elements showing FMRFamide-like immunoreactivity in hydromedusae. Acta Zoologica, 66(*4*), 199–210.

Meech, R.W., & Mackie, G.O. (1993a). Ionic currents in giant motor axons of the jellyfish, *Aglantha digitale*. Journal of Neurophysiology, 69, 884–893. doi: 10.1152/jn.1993.69.3.884

Meech, R.W., & Mackie, G.O. (1993b). Potassium channel family in giant motor axons of *Aglantha digitale*. Journal of Neurophysiology, 69, 894–901. doi: 10.1152/jn.1993.69.3.894

Meech, R.W., & Mackie, G.O. (1995). Synaptic potentials and threshold currents underlying spike production in motor giant axons of *Aglantha digitale*. Journal of Neurophysiology, 74, 1662–1670. doi: 10.1152/jn.1995.74.4.1662

Morise, H., Shimomura O., Johnson F.H., Winant J. (1974). Intermolecular energy transfer in the bioluminescent system of *Aequorea*. Biochemistry. 13 (12): 2656–62. 10.1021/bi00709a028

Norekian, T.P. & Meech, R.W. (2020). Structure and function of the nervous system in nectophores of the siphonophore *Nanomia bijuga*. J Exp Biology, 223(24). 10.1242/jeb.233494

Norekian TP, Meech RW. (2026). Structure and function of the nervous system in the stem of the siphonophore *Nanomia septata*: its role in swimming coordination. J Exp Biology, 229, jeb251974. doi:10.1242/jeb.251974

Norekian, T.P., & Moroz, L.L. (2016). Development of neuromuscular organization in the ctenophore *Pleurobrachia bachei*. Journal of Comparative Neurology, 524, 136–151. 10.1002/cne.23830

Norekian, T.P., & Moroz, L.L. (2019a). Neural System and Receptor Diversity in the ctenophore *Beroe abyssicola*. Journal of Comparative Neurology, 527, 1986–2008. 10.1002/cne.24633

Norekian, T.P., & Moroz, L.L. (2019b). Neuromuscular organization of the Ctenophore *Pleurobrachia bachei*. Journal of Comparative Neurology, 527, 406–436. 10.1002/cne.24546

Norekian, T.P., & Moroz, L.L. (2020a). Comparative neuroanatomy of ctenophores: Neural and muscular systems in *Euplokamis dunlapae* and related species. Journal of Comparative Neurology, 528, 481–501. 10.1002/cne.24770

Norekian, T.P., & Moroz, L.L. (2020b). Atlas of the neuromuscular system in the trachymedusa *Aglantha digitale*: Insights from the advanced hydrozoan. Journal of Comparative Neurology, 528: 1231–1254. 10.1002/cne.24821

Passano, L.M. (1965). Pacemakers and activity patterns in medusae: Homage to Romanes. American Zoologist, 5, 465–481.

Passano. L.M. (1973). Behavioral control systems in medusae: A comparison between hydro- and scyphomedusae. Publications of the Seto Marine Biological Laboratory, 20, 615–645

Roberts, A., & Mackie, G.O. (1980). The giant axon escape system of a hydrozoan medusa, *Aglantha digitale*. Journal of Experimental Biology, 84, 303–318.

Satterlie, R.A. (1985a). Central generation of swimming activity in the hydrozoan jellyfish *Aequorea aequorea*. Journal of Neurobiology, 16, 41–55. doi: 10.1002/neu.480160105

Satterlie, R.A. (1985b). Control of swimming in the hydrozoan jellyfish *Aequorea aequorea*: direct activation of the subumbrella. Journal of Neurobiology, 16, 211–226. doi: 10.1002/neu.480160306

Satterlie, R.A. (2002). Neural control of swimming in jellyfish: A comparative story. Canadian Journal of Zoology, 80, 1654–1669. doi: 10.1139/z02-132

Satterlie, R.A. (2008). Control of swimming in the hydrozoan jellyfish *Aequorea victoria*: subumbrellar organization and local inhibition. Journal of Experimental Biology, 211, 3467–3477. doi: 10.1242/jeb.018952

Satterlie, R.A. (2011). Do jellyfish have central nervous systems? Journal of Experimental Biology, 214, 1215–1223. doi: 10.1242/jeb.043687

Satterlie, R.A. (2015a). Cnidarian Nerve Nets and Neuromuscular Efficiency. Integrative and Comparative Biology, 55, 1050–1057. doi: 10.1093/icb/icv067

Satterlie, R.A. (2015b). The search for ancestral nervous systems: an integrative and comparative approach. Journal of Experimental Biology, 218, 612–617. doi: 10.1242/jeb.110387

Satterlie, R.A. (2017). Cnidarian Neurobiology. The Oxford Handbook of Invertebrate Neurobiology. Edited by J.H. Byrne. doi: 10.1093/oxfordhb/9780190456757.013.7

Satterlie, R.A. (2018). Jellyfish Locomotion. Oxford Research Encyclopedia of Neuroscience. 10.1093/acrefore/9780190264086.013.147

Satterlie, R.A. & Spencer, A.N. (1983). Neuronal control of locomotion in hydrozoan medusae. A comparative study. Journal of Comparative Physiology, 150, 195–206.

Shimomura, O. (1979). Structure of the Chromophore of Aequorea Green Fluorescent Protein FEBS Letters 104(*2*), 220−222.

Shimomura, O. (2005). The discovery of aequorin and green fluorescent protein. Journal of Microscopy 217, 3–15. 10.1111/j.0022-2720.2005.01441.x

Shimomura, O., Johnson, F. H., & Saiga, Y. (1962). Extraction, Purification and Properties of Aequorin, a Bioluminescent Protein from the Luminous Hydromedusan, *Aequorea*. Journal of Cellular and Comparative Physiology, 59(*3*), 223–239. doi:10.1002/jcp.1030590302

Shimomura, O., Johnson, F. H., & Saiga, Y. (1963). Extraction, Purification and Properties of Halistaurin, a Bioluminescent Protein from the Hydromedusan *Halistaura*. Journal of Cellular and Comparative Physiology, 62(*1*), 9–15. 10.1002/jcp.1030620103

Singla, C.L. (1978a). Fine structure of the neuromuscular system of *Polyorchis penicillatus* (Hydromedusae, Cnidaria). Cell and Tissue Research, 193, 163–174. doi: 10.1007/bf00221609

Singla, C.L. (1978b). Locomotion and neuromuscular system of *Aglantha digitale*. Cell and Tissue Research, 188, 317–327. doi: 10.1007/bf00222640

Singla, C.L. (1983). Fine structure of the sensory receptors of *Aglantha digitale* (Hydromedusae: Trachylina). Cell and Tissue Research, 231, 415–425. doi: 10.1007/bf00222191

Spencer, A.N. (1975). Behavior and electrical activity in the hydrozoan *Proboscidactyla flavicirrata* (Brandt). II. The medusa. Biological Bulletin, 149, 236–250.

Spencer, A.N. (1978). Neurobiology of *Polyorchis*. I. Function of effector systems. Journal of Neurobiology, 9(*2*), 143–157.

Spencer, A.N. (1979). Neurobiology of *Polyorchis*. I. Structure of effector systems. Journal of Neurobiology, 10(*2*), 95–117.

Spencer, A.N. (1982). The physiology of coelenterate neuromuscular synapse. Journal of Comparative Physiology, 148, 353–363.

Spencer, A.N. & Arkett, S.A. (1984). Radial symmetry and the organization of central neurones in a hydrozoan jellyfish. Journal of experimental Biology, 110, 69–90.

Spencer, A.N. & Satterlie, R.A. (1980). Electrical and dye coupling in an identified group of neurons in a coelenterate. Journal of Neurobiology, 11(*1*), 13–19.

Spencer, A.N. & Satterlie, R.A. (1981). The action potential and contraction in subumbrella swimming muscle of Polyorchis penicillatus. Journal of Comparative Physiology, 144, 401–407.

Weber, C. (1982). Electrical activities of a type of electroretinogram recorded from the ocellus of a jellyfish, *Polyorchis penicillatus* (Hydromedusae). Journal of Experimental Zoology, 223, 231–243. doi: 10.1002/jez.1402230305

Weber, C., Singla, C.L., & Kerfoot, P.A. (1982). Microanatomy of the subumbrellar motor innervation in *Aglantha digitale* (Hydromedusae: Trachylina). Cell and Tissue Research, 223, 305–312. doi: 10.1007/bf01258491

Wehland, J., & Willingham, M.C. (1983). A rat monoclonal antibody reacting specifically with the tyrosylated form of alpha-tubulin. II. Effects on cell movement, organization of microtubules, and intermediate filaments, and arrangement of Golgi elements. Journal of Cell Biology, 97, 1476–1490. 10.1083/jcb.97.5.1476

Wehland, J., Willingham, M.C., & Sandoval, I.V. (1983). A rat monoclonal antibody reacting specifically with the tyrosylated form of alpha-tubulin. I. Biochemical characterization, effects on microtubule polymerization in vitro, and microtubule polymerization and organization *in vivo*. Journal of Cell Biology, 97, 1467–1475. 10.1083/jcb.97.5.1467

Wrobel, D. & Mills, C. (1998). Pacific Coast Pelagic Invertebrates: A guide to the common gelatinous animals. Sea Challengers Publication, Editor Hans Bertsch Dr., 108p. ISBN 0-930118-23-5

Wulf, E., Deboben, A., Bautz, F.A., Faulstich, H., & Wieland, T. (1979). Fluorescent phallotoxin, a tool for the visualization of cellular actin. Proceedings of the National Academy of Sciences U S A, 76, 4498–4502. 10.1073/pnas.76.9.4498

